# Memory traces bias new learning for hippocampal generalization

**DOI:** 10.1101/2025.11.24.690297

**Authors:** Fish Kunxun Qian, Guanchun Li, David Lipshutz, Sandro Romani, Jeffrey C. Magee

**Affiliations:** Howard Hughes Medical Institute, Baylor College of Medicine; Houston, TX, USA; Department of Neuroscience, Baylor College of Medicine; Houston, TX, USA; Neuroengineering Initiative, Rice University; Houston, TX, USA; Howard Hughes Medical Institute, Janelia Research Campus; Ashburn, VA, USA; Jan and Dan Duncan Neurological Research Institute, Texas Children’s Hospital; Houston, TX, USA

## Abstract

Generalizing prior experience to guide behavior in novel situations is a fundamental cognitive function, yet how this occurs remains poorly understood. Here we combined longitudinal imaging in mice with modeling and circuit perturbation to examine generalization in hippocampal CA1 across environments with shared task structure. Prior training accelerated behavioral adaptation and preserved cross-environment representations. We identified putative memory traces associated with generalization. At the cellular level, spatially informative residual activity reflecting generalizable information biased which neurons formed place fields (PFs) and where via behavioral timescale synaptic plasticity (BTSP). At the population level, BTSP was associated with rapid alignment of novel-environment network dynamics to familiar neural subspaces. Chemogenetic suppression of local CA1 during training reduced residual activity, BTSP induction, place cell recruitment and generalization. In sum, the influence of memory traces on new PF formation may allow past experience to guide new learning to form representations based on generalizable features.

## Introduction

Generalizing learned knowledge to navigate novel conditions represents a cornerstone of intelligent behavior(*1*). The mammalian hippocampus integrates spatial(*2*), temporal(*3*), self-motion(*4*), and task-related information(*5–9*) from experience into coherent cognitive maps(*10*–*15*). There exists a fundamental challenge here: how to maintain specificity for distinct episodes(*16*, *17*) while forming generalizable, transferable patterns(*18–26*). Classical theory(*18*) proposes that the hippocampus is responsible for rapid encoding of episodic memory while the neocortex slowly extracts regularities from experience for generalization. Yet mounting evidence suggests that the hippocampus itself may enable rapid generalization by extracting common structure across experiences and applying this structure to novel situations(*19*, *24–29*). The neural mechanisms underlying this process remain elusive.

Recent discoveries suggest two potentially interrelated mechanisms. First, neural population activity is often organized within low-dimensional neural manifolds(*30–34*), or subspaces, and hippocampal population geometry can encode task-relevant structure(*6*, *25*, *26*, *35*). In this framework, orthogonal subspaces separate distinct memories while shared dimensions capture generalizable information. Yet, how generalizable neural subspaces emerge in novel contexts remains unknown. Second, synaptic plasticity is believed to underlie adaptive behavior(*36–39*) and recent work has shown that behavioral timescale synaptic plasticity (BTSP) mediates rapid hippocampal learning(*40–43*). BTSP in CA1 neurons is triggered by dendritic calcium plateau potentials (plateaus) that rapidly, in one shot, potentiate CA3 inputs active within seconds of the plateau(*40*, *44–47*), allowing the linkage of overlapping experiences into coherent memories(*7*, *43*). It has been established that BTSP contributes to PF reorganization within a familiar environment and that CA1 place cells (PCs) flexibly reference spatial and reward-associated cues depending on experience(*7*). However, whether, and if so how, prior experience extracts common structure across multiple experiences to shape future plasticity in a novel context remained unknown. An intriguing property of the BTSP credit assignment procedure is that depolarization present in weak memory traces active at the beginning of new experiences biases which CA1 neurons will express PFs and at what particular location(*7*, *43*, *48–50*). This element of BTSP could allow past learning to influence new learning and in this way enable generalizable neural subspaces to emerge. We explore these issues below.

## Results

### Behavioral and neural generalization

To study how the hippocampus generalizes, we began with a re-analysis of our previously published data set(*7*), focusing on cross-environment generalization rather than the within-environment reference-frame dynamics examined previously. Here, head-fixed mice were trained on a hippocampal-dependent spatial learning task(*41*, *51*) where they ran in complete darkness on a 180-cm linear treadmill that featured a set of tactile cues for water rewards at a fixed location (50 cm; Fig. 1A). After several days of training in one environment (a cue-rich belt; 700±38 laps, in 5.4±0.4 days, n = 9 mice; fig. S1A), mice demonstrated spatial learning through anticipatory slowing and spatially selective licking before the reward location (Fig. 1B to 1D). Two-photon Ca^2+^ imaging of GCaMP6f-expressing pyramidal neurons in dorsal hippocampal CA1 revealed spatially tuned place cells (PCs) tiling the entire track (Fig. 1A; 1G, top; 1H, top), as shown previously(*7*).

**Fig. 1.**
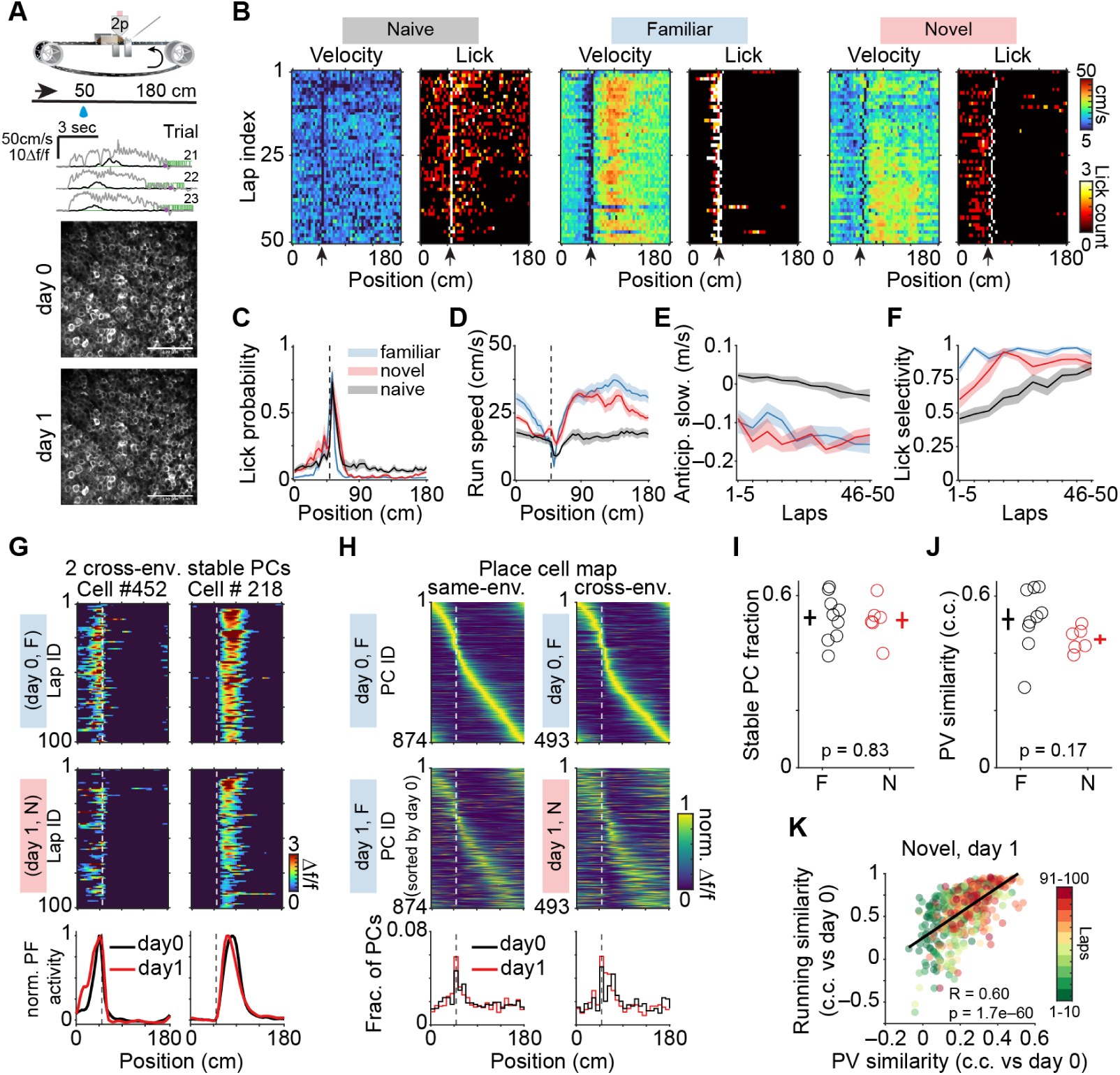
Behavioral and neural generalization. **(A)** Top, head-fixed mouse on a 180-cm linear treadmill with reward at 50 cm. Middle: Δf/f traces (black), velocity (grey), and licks (green) across 3 laps. Purple dots mark reward. Bottom: representative two-photon (2p) field of view showing GCaMP6f-expressing neurons tracked across days. Scale bar, 100 *μm*. **(B)** Velocity and lick heatmaps across space and laps for representative naïve (left), familiar (F, same-environment (Same), middle), and novel (N, cross-environment (Cross), right) mice. Naïve: first day over 50 laps/session; familiar/novel: day 1. Arrow: reward location. **(C)** Mean spatial lick probability for naïve (grey, n = 10 mice), novel (red, n = 6 mice), and familiar mice (blue, n = 9 mice). **(D)** Mean running speed (format as in (C)). **(E)** Anticipatory slowing across sections, with each section corresponding to mean across 5 laps. Lines and shading indicate mean±sem. n = 10, 6, 9 mice for naïve, novel, and familiar groups, respectively. Repeated two-way ANOVA with group as the between-mouse factor, and section as the repeated within-mouse factor. Novel versus naïve: group p = 1.3e–5. Novel versus familiar: group p = 0.64. **(F)** Licking selectivity (format and mice as (E)). Repeated two-way ANOVA, Novel versus naïve: group p = 0.015. Novel versus familiar: group p = 6e–3, group x section p = 0.044. Because naïve mice tended to run less per session, we took the first 50 laps for cross-condition comparisons for (B) to (F). **(G)** Two cross-environment stable cells (PF position difference ≤15 cm). Top/middle: spatial Δf/f activity across laps on day 0 (familiar env. A, (“F”)) and day 1 (novel env. B, (“N”)). Bottom: Peak-normalized (norm.) mean Δf/f across space for day 0 (black) and day 1 (red). **(H)** Left, peak-normalized spatial Δf/f for same-environment PCs for two days sorted by day 0 PF peak locations (n = 874 PCs, 9 mice), with PC fraction versus peak position below. Right, Cross-environment (n = 493 PCs, 6 mice). F/N: familiar/novel environments. **(I)** Stable PC fraction; familiar (“F”): 0.52±0.027 (n = 9 mice); novel (“N”): 0.51±0.029 (n = 6). p = 0.83, unpaired *t*-test. Circles: individual mice. Crosses: mean±s.e.m. **(J)** Population vector (PV) similarity (Pearson’s correlation coefficients (*c*. *c*.) between day 1 and day 0 PVs). Familiar (“F”): 0.52±0.037, (n = 9 mice); novel (“N”): 0.45±0.017 (n = 6 mice); p = 0.17, unpaired *t*-test, using reliable PCs on both days. **(K)** Trial-by-trial running versus PV similarity (both versus day 0 in familiar env. A) for novel env. B (day 1, n = 600 laps, 6 mice). Black line: linear fit. R = 0.6, p = 1.7e–60. PV similarity used reliable PCs from both days. Vertical dashed lines in (C,D,G,H) mark reward locations (50 cm).

To test for generalization, we exposed mice trained for several days (718±31 laps, in 5.7±0.3 days, n = 6 mice) in the familiar environment (belt A) to a novel environment (belt B with different tactile cues) while the reward structure remained constant (run laps for reward at particular location). Below we refer to this condition as “cross-environment” and compare these mice with “same-environment” controls that experienced the same familiar environment on subsequent days. The cross-environment mice exhibited stronger anticipatory slowing and licking selectivity than naïve controls across lap sections (anticipatory slowing: group effect, p = 1.3e–5, Fig. 1E; licking selectivity: group effect, p = 0.015, Fig. 1F; two-way repeated ANOVA). Moreover, cross-environment mice matched same-environment controls in anticipatory slowing (group effect, p = 0.64, Fig. 1E), while licking selectivity progressively approached same-environment levels across the session (group × section interaction, p = 0.044; Fig. 1F, two-way repeated ANOVA). Thus, mice exhibited robust behavioral generalization despite altered sensory cues.

To examine whether CA1 neural representations generalize across environments, we longitudinally tracked the same cells across days (Fig. 1A) and found that approximately half of the PCs maintained stable PFs across environments (familiar to novel; Fig. 1G; 1H, right), matching same-environment levels (familiar to familiar; Fig. 1H, left; 1I). Additionally, PCs exhibiting the same PF location (stable cells) concentrated near the reward (Fig. 1G; fig. S1B and S1D) with a three-fold higher density than non-rewarded zones (fig. S1D), whereas remapped cells tiled the environment more uniformly (fig. S1C and S1D). Consistent with this, cross-environment population vector (PV) correlations were similar to same-environment controls (Fig. 1J), with a strong correlation band along the entire diagonal of group-mean cross-environment correlation matrix (fig. S1E). The group-mean PV correlation peaked near zero shift (mean peak shift = −1.2 cm, p = 0.88, Wilcoxon signed-rank test versus zero; fig. S1H), and the signed optimal shift did not show a systematic lap-by-lap drift (Spearman ρ = −0.12, p = 0.61; fig. S1I), indicating that cross-belt stability primarily reflects preserved reward-relative organization rather than a global shift independent of reward anchoring. Critically, neural activity similarity across environments tightly coupled with running profile similarity trial-by-trial (Fig. 1K), as both improved with experience (fig. S1F, novel; fig. S1G for familiar environment control), consistent with previous theoretical work(*52*). Thus, stable task-relevant CA1 representations co-emerge with successful behavioral generalization.

### Preserved low-dimensional structure across environments

Single cell selectivity and population structure offer two distinct and complementary perspectives to describe neural code(*32*, *33*). We next asked whether the population-level structure of CA1 activity was preserved across environments. To test whether spatial information transferred across environments, we reanalyzed the data from our previous study and decoded the position of one set of mice from novel environment activity (cross-environment) or another set from the familiar environment activity (same-environment) using Bayesian decoders trained exclusively on the previous day’s activity in the familiar environment. Position predictions from cross-environment decoders were above chance (100-lap mean: 26.6±2.1cm versus 45cm chance, p = 3.1e–4, one-sample *t*-test, n = 6 mice). Further, cross-environment decoding accuracy was lower than same-environment controls (group, p = 1.1e–3; Fig. 2A) but this difference decreased with experience (group × section interaction, p = 4.4e–3; Fig. 2A), suggesting spatial information recovers through learning.

**Fig. 2.**
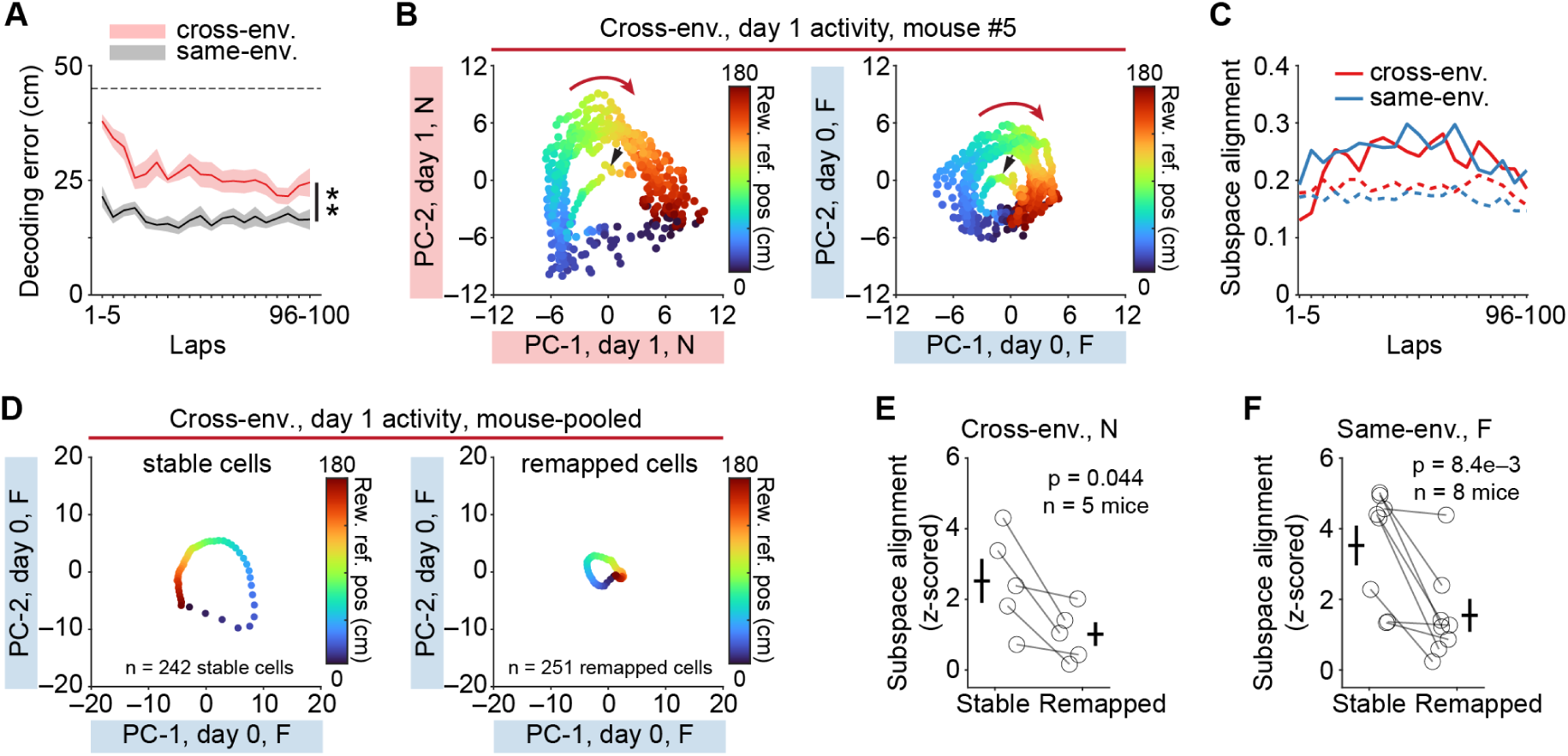
Preserved low-dimensional structure across environments supports fast generalization. **(A)** Mean decoding error across sections (5-lap mean). Cross: 26.6±2.1 cm (n = 6 mice); Same: 16.8±1.4 cm (n = 9); Dashed line: chance. Shading: S.E.M. Two-way repeated ANOVA: group, p = 1.1e–3; group × section interaction, p = 4.4e–3. **(B)** Example 10-lap mean day 1 (novel env. B) activity projected onto 2 principal components (PC-1, PC-2) from day 1 (left) or day 0 (right, familiar env. A), colored by reward distance. Black arrow: laps 1-10; red arrow: running direction; F/N: familiar/novel environments. **(C)** Mean subspace alignment across sections (5 laps/section). Cross: red, n = 6 mice; Same: blue, n = 9. Dashed lines: 95th percentile of shuffle distribution (1000 iterations of randomly selected 2D subspaces from day0 principal components), representing the upper bound expected by chance. Shading: SEM. **(D)** Day 1 novel session-mean activity from stable (left, n = 242 cells) and remapped (right, n = 251 cells) cells projected onto day 0 familiar subspaces (PC-1, PC-2), mouse-pooled. F: familiar environment. **(E)** Mean z-scored subspace alignment (Cross mice). Stable: 2.5±0.61; remapped: 1.0±0.34; p = 0.044, paired *t*-test, n = 5 mice (one excluded: <20 cells). Circles: mice; crosses: mean±sem. N: novel environment. z-scoring takes mean and standard deviation of subspace alignment calculated using randomly selected 2 subspaces (1000 times) from day 0 principal components. **(F)** Same-environment z-scored subspace alignment as (E). Stable: 3.5 ± 0.56; remapped: 1.6±0.47; p = 8.4e–3, paired *t*-test, n = 8 mice (one excluded: <20 cells). F: familiar environment.

Recent work demonstrates that hippocampal neural trajectories are confined to low-dimensional subspaces(*6*, *25*, *26*, *35*, *53*). To test if neural activity subspaces are involved in generalization, we identified leading dimensions (i.e., principal components) using principal component analysis (PCA; fig. S2A and S2B; see Methods). Neural trajectories in the novel environment formed continuous rings preserving circular task topology that were visualizable when color-coded by reward distance (Fig. 2B, left). This structure remained largely intact when projecting novel environment activity onto the principal components of neuronal activity recorded on the previous day in the familiar environment (Fig. 2B, right), even in the earlier laps (Fig. 2B, black arrow). Same-environment controls showed fully expanded rings by 10 laps of running (fig. S2C). Indeed, a subspace alignment index (5 laps per section) for cross environment sessions started low during laps 1-10, increased sharply by laps 11-15, and then plateaued at high levels (Fig. 2C, dashed lines mark the 95% upper bounds from random shuffles). Subspace alignment significantly exceeded chance level in both conditions (fig. S2D, p<0.05 in all sections for same-environment and from sections 3 to 20 for cross-environment mice). Principal angle analysis corroborated this rapid alignment (fig. S2E). Notably, pairwise neural co-activity levels in the familiar environment significantly predicted their corresponding co-activity levels in the novel environment during the first 20 laps (fig. S2F, left; S2F, right, same-environment controls). Additional representational similarity analyses were consistent with this preservation (fig. S2L, S2M). To characterize what the top 2 subspaces capture, we examined which cells dominated the loading weights of PC-1 and PC-2 from the day 0 PCA. For each mouse, we classified cells as high-weight (top third) and low-weight (bottom third) by their mean absolute loading weight across PC-1 and PC-2 (fig. S3A). High-weight cells exhibited substantially higher lap-to-lap place field reliability (fig. S3B), earlier place field onset (fig. S3C), and greater spatial information (fig. S3D) compared to low-weight cells, and their PFs tiled the environment broadly rather than clustering near reward (fig. S3E), suggesting that the top subspace reflects distributed spatial coding structure. Notably, high-weight cells also exhibited significantly higher cross-day PF similarity (fig. S3F), despite the day 0 PCA being blind to day 1 data, indicating that the familiar-environment covariance structure preferentially captures cells whose spatial tuning generalizes across environments. Together, preserved co-activity in a novel-context rapidly aligns with established task-relevant subspaces over a similar time-course as the behavioral adaptation to the novel environment (within 10-20 laps).

Finally, projecting the activity in the novel environment of only the stable set of PCs onto the top two components from the familiar environment also revealed visibly more expanded rings (Fig. 2D, left) than remapped cells (Fig. 2D, right; fig. S2H for controls). When projected onto their own same session components, however, both cell types exhibited well-organized trajectories (fig. S2G and S2I) with comparable variance within each day (fig. S2J and S2K). Ring expansion approaching same-day projection levels indicates preserved subspace structure. Indeed, stable cells showed significantly stronger subspace alignment than remapped cells (Fig. 2E and 2F). Thus, it appears that CA1 stable cells primarily support the subspace alignment for generalization, whereas remapped cells mediate subspace reorganization separating distinct memories.

### Reference frame switching actively supports generalization

What common information shapes generalization across familiar and novel experiences? Recent studies have shown that both spatial and goal referencing support CA1 hippocampal maps(*7*, *8*). To examine which reference frames were involved here, we reanalyzed the data from our previous study(*7*) where the reward location was shifted by 90 cm in both environments following the initial 100 trials of exploration. In the familiar environment, PCs initially maintained their original positions on the track despite the reward shift (fig. S4A and fig. S4E). Field position changes clustered tightly around zero (median: 18 cm, fig. S4A), yielding robust spatial-cue-aligned correlations but negative reward-aligned correlations (fig. S4G). Running profiles showed parallel patterns (fig. S4B, top; fig. S4H; fig. S4I, left). Following this initial spatial anchoring, both neural activity and running behavior modestly shifted in parallel to track the new reward location (half-maximum: 15 laps; fig. S4C; fig. S4J, left). In the novel environment, however, PCs immediately shifted following the reward location change (fig. S4A; fig. S4F). Field position changes clustered near 90 cm (median: 72 cm; fig. S4A), producing negative spatial-cue-aligned PF correlations but robust reward-aligned correlations (fig. S4G). Running profiles mirrored this immediate shift (fig. S4B, bottom; fig. S4H; fig. S4I, right), with remarkably fast behavioral adaptation (half-maximum: 2 laps; fig. S4D; fig. S4J, right). Population imaging of CA3 axons showed a similar environment-dependent reference frame reorganization that switched from spatial-cue alignment in the familiar environment to reward alignment in the novel environment (fig. S5). This suggests that the flexible place-cell referencing in CA1 could be inherited, at least partially, from CA3 input, consistent with previous work(*7*). It remains unclear, however, whether in our conditions CA3 inputs generalize across environments because the same axons were not tracked across environments. PCA analysis of concatenated neural data from five mice that experienced reward shifts in both environments (day 1, familiar A; day 2, novel B) suggested that CA1 simultaneously represents task-relevant (i.e., reward distance; fig. S4K and S4M) and context-specific (i.e., environment identity; fig. S4L and S4N) information but in separate subspaces. Together, the substantial reference-frame switching and the environment-specific activity indicate that cross-environment PF stability reflects active reorganization rather than rigid reward-distance coding, and the goal-referenced synaptic input could provide the generalizable information common to both current and past experiences.

### BTSP is involved in building population activity

Recent work has shown that BTSP, triggered by dendritic plateau potentials, rapidly forms new PFs in CA1 and new BTSP is required to produce robust PF representations (*40–42*, *45*, *54*), even during repeated experience in the same environment(*48*). Our previous study showed that BTSP rapidly forms new PFs during PC remapping in a familiar environment in response to a reward location switch(*7*). It remained unclear whether this plasticity is systematically influenced by prior experience. To answer this question, we first asked whether BTSP is the mechanism that builds the aligned population activity described above for generalization. To do this, we identified putative BTSP events through established signatures(*7*, *40–42*, *48*, *54*) (fig. S6B to S6D; summarized in fig. S6A) and found that most PCs exhibited BTSP regardless of stability (83% of stable cells and 84% of remapped cells with comparable event frequencies; Fig. 3A; fig. S6A). However, in both novel and familiar environments stable cells’ initial BTSP events were biased toward their previous session PF locations, while remapped cells showed no such bias (Fig. 3B). The stability between initial BTSP events and previous session PFs strongly predicted cross-environment PF stability (Fig. 3C), indicating that the spatial location of BTSP is a strong determinant of new PF location, consistent with previous studies(*41*, *42*, *45*). BTSP-positive PCs were enriched near the reward location on the novel belt (fig. S6E), with early-onset PFs and first BTSP events clustering near reward during laps 1-20, followed by fewer and spatially broader recruitment of PFs over later laps (fig. S6F; S6G).

**Fig. 3.**
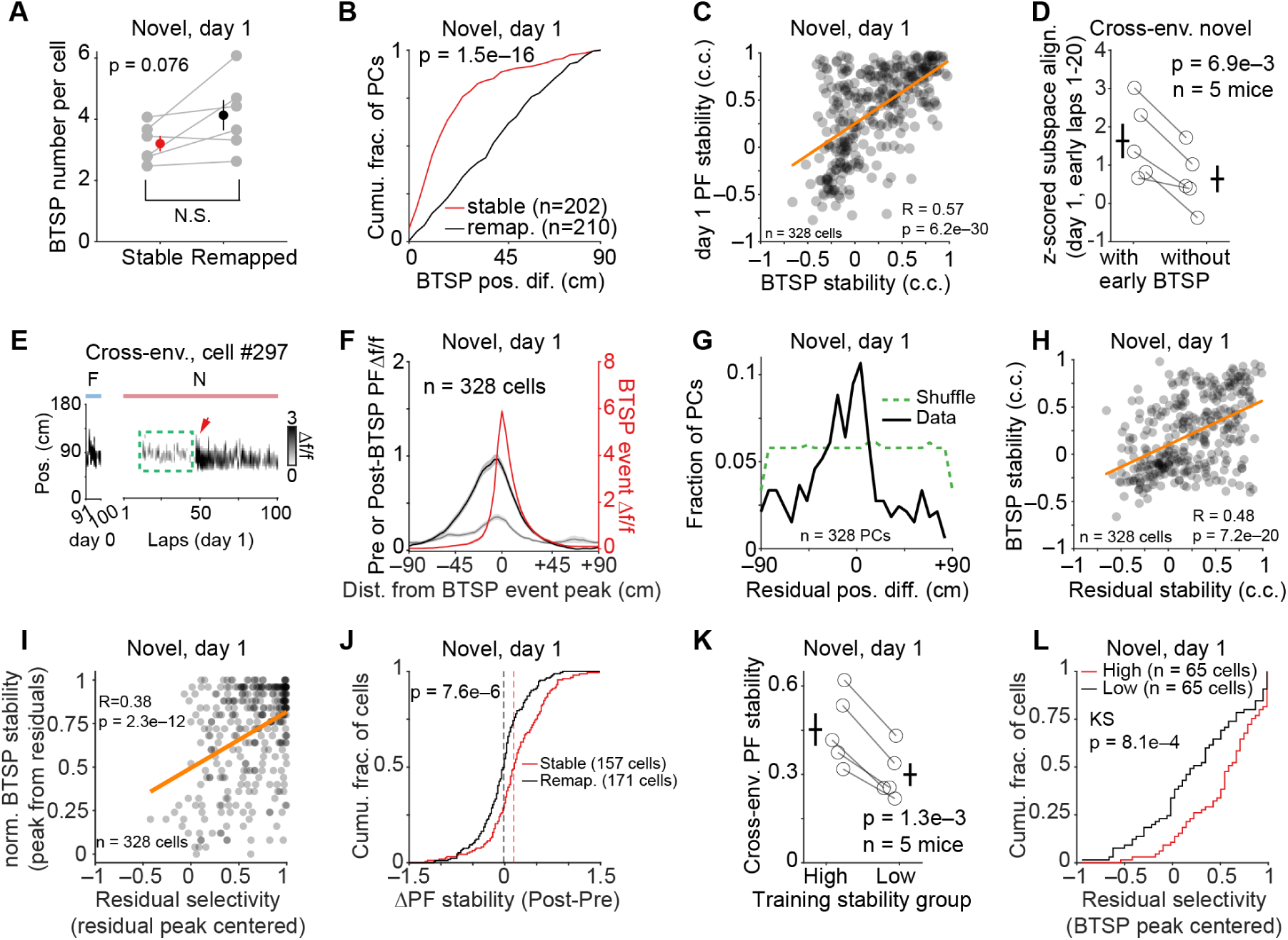
Residual activity guided BTSP for generalization. **(A)** BTSP events per cell (100 laps, day 1, novel env. B). Stable versus remapped cells: 3.2±0.25 versus 4.1±0.5; p = 0.076, paired *t*-test, n = 6 mice for each group. **(B)** Cumulative fraction versus position difference between day 1 (novel env. B) first BTSP event peak and day 0 (familiar env. A) PF peak. Stable: n = 202 cells, 6 mice; remapped: n = 210 cells, 6 mice. p = 1.5e–16, two-sample Kolmogorov-Smirnov test. **(C)** Day 1 (novel env. B) PF stability versus initial BTSP stability (both *c*. *c*. versus day 0 PF). Orange line: linear fit; R = 0.57, p = 6.2e–30, n = 328 cells, 6 mice. Note, 25 of 412 cells in (B) lacked usable lap data (e.g., first-lap plateaus), and 59 of the remaining 387 lacked significant pre-BTSP events, see fig. S6A and Methods. **(D)** Z-scored subspace alignment between day 1 (novel env. B, early laps 1-20) and day 0 (familiar env. A) population activity, for cells with versus without early BTSP (earlier vs later half of first BTSP laps within each mouse; Methods) in cross-environment mice. Z-scores were computed relative to 1000 shuffles of randomly paired subspaces from day 0 (computed separately for each subpopulation), enabling comparison across groups with different population structure. With: 1.63±0.45; without: 0.64±0.35; p = 6.9e–3, paired *t*-test, n = 5 mice. One excluded (fewer than 20 cells per either group). **(E)** An example Δf/f heatmap from a cross-environment stable cell. Green box: residual activity near day 0 PF. Red arrow: subsequent BTSP event. F/N: familiar/novel environments. **(F)** Population-averaged spatial tuning aligned to the first BTSP event peak (position 0) for all residual-active PCs on day 1 (novel env. B). For each cell, the mean Δf/f spatial profile from the 5 laps preceding the first BTSP event (Pre, grey) the 5 laps following (Post, black), and the BTSP event lap (red) were circularly shifted so that the BTSP event peak was centered. Lines and shading: mean±sem. n = 328 cells, 6 mice. **(G)** Cell fraction versus position difference between day 1 (novel env. B) residual activity peak (mean, 5 laps pre-BTSP) and day 0 PF peak. Dashed: top 5% from random model. n = 328 cells. **(H)** Residual stability (*c*. *c*. versus day 0 PF) and initial BTSP event stability (*c*. *c*. versus day 0 PF) in novel environment B (day 1). Orange line: linear fit. R = 0.48, p = 7.2e–20, n = 328 cells. **(I)** Normalized BTSP proximity to residual peak versus residual selectivity (centered on residual peak) on day 1 (novel env. B). For each cell, residual selectivity was computed as (mean in-field Δf/f – mean out-of-field Δf/f) / (mean in-field Δf/f + mean out-of-field Δf/f). BTSP proximity was normalized so that 1 indicates same location and 0 indicates maximum separation by 90 cm. Orange line: linear fit; R = 0.38, p = 2.3e–12, n = 328 cells, 6 mice. **(J).** Cumulative distribution of ΔPF stability (Post – Pre) for stable (red, 157 cells) and remapped (black, 171 cells) PCs on day 1 (novel env. B). ΔPF stability quantifies the change in PF similarity (c.c. versus day 0 PF) from Pre and Post described above. Positive values indicate that BTSP moved spatial tuning closer to the day 0 template. Dashed lines: median for each group. p = 7.6e– 6, two-sample Kolmogorov-Smirnov test. **(K)** Cross-environment PF stability (*c*. *c*. between day 0 and day 1 PFs) for high (top 20%) versus low (bottom 20%) training stability (see Methods). High: 0.45±0.055; low: 0.30±0.038; p = 1.3e– 3, paired *t*-test, n = 5 mice. Circles: mice; crosses: mean±s.e.m. **(L)** Cumulative distribution of residual selectivity (centered on BTSP event peak) for high-(red, top 20%, n = 65 cells) versus low-training-stability (black, bottom 20%, n = 65 cells) PCs on day 1 (novel env. B). Residual selectivity was computed as in (I) but with in-field and out-of-field defined relative to the BTSP event peak position. Out of all day 0 PCs, ∼60% had BTSP and available residual laps on day 1 in a novel environment. Training stability: mean pairwise PF correlation across 3 prior training days (top/bottom 20% defined per mouse; see Methods). p = 8.1e–4, two-sample Kolmogorov-Smirnov test.

To test if BTSP is involved in aligning activity in the novel environment with familiar environment activity subspaces, we split PCs from the novel environment into subgroups that did or did not show early BTSP signatures (see Methods). In cross-environment mice, cells with early BTSP had stronger subspace alignment than those without during the first 20 laps (Fig. 3D), and a similar difference was observed in same-environment controls (fig. S6H). Notably, both subgroups had comparable fractions of stable cells (fig. S6I and S6J). Subspace alignment during late laps (81–100) showed no significant differences between subgroups (fig. S6K and S6L). Together, our results suggest that early BTSP is preferentially associated with the rapid subspace alignment observed during generalization.

### Memory traces guide BTSP to task-relevant selectivity

How is BTSP able to produce a recognizable representation in a novel environment? BTSP induction appears to be driven by a dendritic computation featuring a population level target signal from the entorhinal cortex, a feedback inhibitory input reflecting the actual CA1 population activity and a weak memory trace that could provide cell specificity(*41*, *43*, *48*). While these traces typically do not drive strong PFs, they can provide spatially-informative depolarization that potentially lowers the plateau initiation threshold, biasing in which cells and where BTSP events occur. We therefore next examined recordings for hallmarks of these memory traces which present as low amplitude, inconsistent activity that precedes the above described BTSP induction events (we refer to these as residuals or residual activity)(*7*, *48–50*, *55*). In CA1 recordings, approximately 80% of novel environment PCs exhibiting BTSP showed detectable activity within five laps preceding induction (stable: 157/202; remapped: 171/210; Fig. 3E, 3F; fig. S7A, S7B, S7E, S7F; see Methods). We also observed PCs with no detectable calcium events during pre-BTSP laps (“silent residuals”, fig. S7C; S7G) and PCs whose first BTSP event occurred during the first lap (fig. S7D; S7H). In addition, residuals occurred at positions similar to previous session PF locations (Fig. 3G; S7A, S7B), capturing generalizable spatial information. Indeed, residual peaks were located near subsequent BTSP events (fig. S6M) and residual stability to the previous-session PFs significantly predicted subsequent BTSP stability (Fig. 3H). Further, cells with more selective residual activity exhibited BTSP events closer to the residual peak (Fig. 3I), suggesting that both the location and selectivity of residuals were critical for memory-trace-guided plasticity events. Consistent with this, stable cells’ residuals were more stable relative to previous sessions than remapped cells (fig. S6N), though both predicted their eventual current session PFs equally well (fig. S6O). As a result, BTSP shifted PF tuning toward the familiar-environment template significantly more for stable cells than for remapped cells (Fig. 3J). The main results were robust when spatial activity maps were constructed using a transient-based pipeline (fig. S8; see Methods). These data suggest that memory traces, manifest as weak but spatially-tuned activity, influence which neurons will be PCs and at what PF locations, even during the exposure to a previously unexperienced environment.

We next reasoned that if the above residual activity reflects an actual memory trace formed during past learning then PCs with the most stable PFs during previous learning episodes should generalize best. To test this, we analyzed mice (n = 5) where we tracked cells across three consecutive training days in the familiar environment before exposure to the novel environment and quantified each cell’s “training stability” as mean pairwise PF correlations across training days. Indeed, cells in the top 20% of training stability showed substantially higher cross-environment stability compared to the bottom 20% (PF: Fig. 3K; S6P; residual stability: fig. S6Q; S6R; residual selectivity: Fig. 3L; S6S). Together, these results demonstrate that residual activity serves as a putative memory trace biasing BTSP in novel contexts, enabling prior experience to systematically guide new learning toward task-relevant features to support hippocampal generalization.

### Realistic simulation exploring elements mediating generalization

To explore a scheme where neuronal dendrites find common elements among different experiences and use them, via BTSP modification of synaptic weights, to rapidly produce useful new representations based off those common elements, we developed a detailed computational network model. The model featured a set of postsynaptic compartmentalized CA1 neurons that received segregated excitation that varied with environment from: 1) CA3 input streams to the proximal integrative compartment and 2) entorhinal cortex (EC3) inputs to the distal compartment as well as 3) distal dendritic inhibition that was a function of population CA1 activity. CA3 synapses were adjusted by the BTSP weight change rule and cascade-like synaptic consolidations to generate synaptic memory traces in the CA3 input (Fig. 4A to 4C; fig. S9A; see Methods). Simulated repeated experience produced recurring potentiation of CA3-CA1 synapses that, in turn, drove these synapses through progressive transitions into increasingly stable biochemical states, making them more resistant to further weight change(*48*, *56*, *57*) (Fig. 4C). Plateau initiation and thus BTSP induction was a function of the above three elements (EC3 excitation, feedback inhibition and excitation from the CA3 memory trace; Fig. 4A; see Methods). Following a transition to a simulated novel environment after repeated simulations in a familiar environment, the network model showed appropriate levels of generalization, quantified as substantial fractions of stable PCs in both same-environment and cross-environment conditions (Same versus Cross, 52±0.4% versus 44±0.5%, p = 4.3e–13, unpaired *t*-test). In addition, the spatial profile of the PFs was comparable to that observed in mice (Fig. 4D; fig. S9B to S9F).

**Fig. 4.**
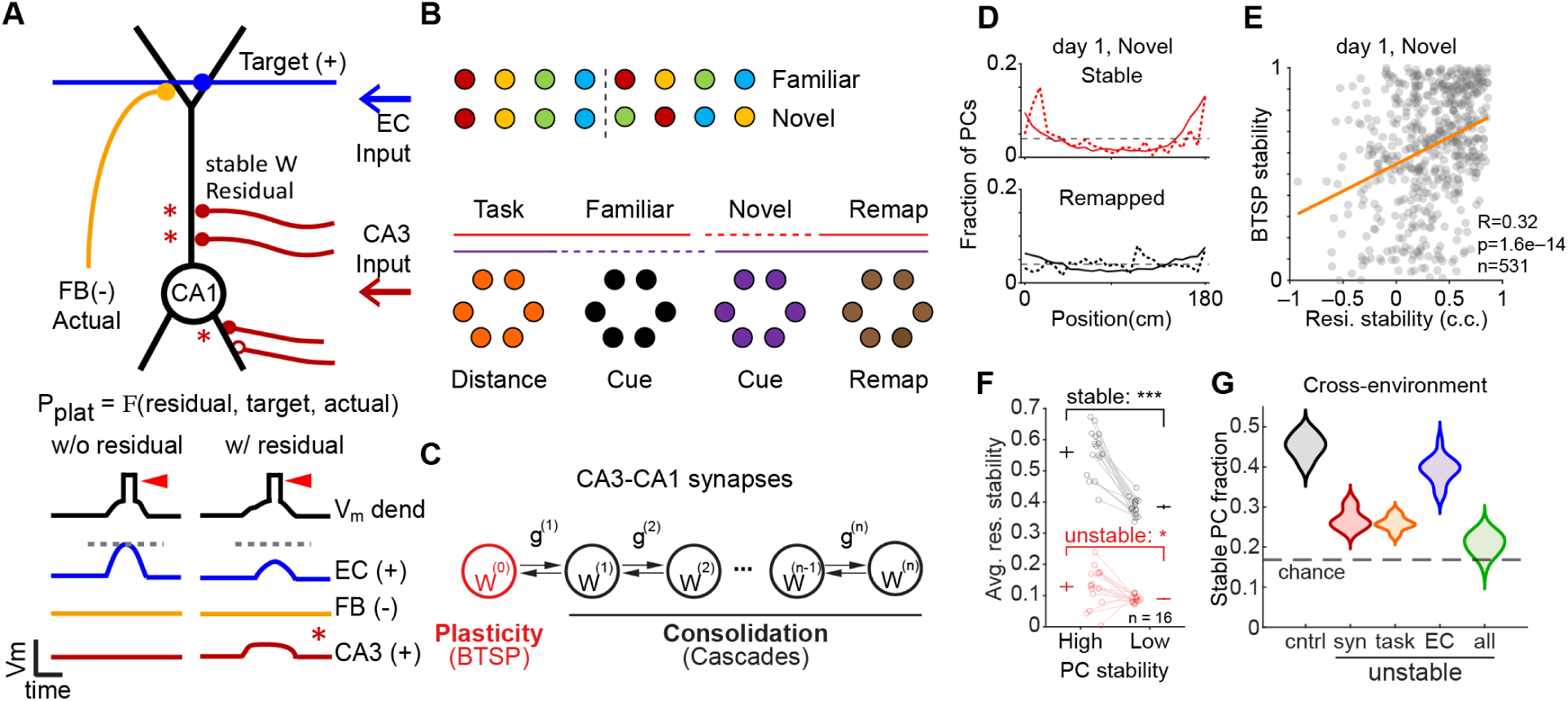
Realistic simulation exploring elements mediating generalization. **(A)** BTSP mechanism schematic. CA1 pyramidal neuron receives three inputs affecting plateau potential initiation probability (*P_plat_*): EC input providing target signal (blue, Target), CA3 input providing residual depolarization from stabilized synapses (red; asterisks indicate stable synapses, filled circles), and feedback inhibition (orange, FB, Actual). Below, voltage traces (V_m_) illustrate how the combination of EC input, feedback inhibition, and CA3 residual activity determines BTSP induction timing and probability. Two conditions: “w/o residual” shows plateau (red triangle) requiring strong EC input alone; “w/ residual” shows that CA3 residual depolarization reduces the plateau threshold (grey dashed line), requiring less EC input. This mechanism enables past experience stored in CA3-CA1 synaptic weights to bias future plasticity toward generalizable features. **(B)** Schematic showing EC (top) and CA3 (bottom) inputs. Top: color scheme indicates EC input changes across environments; dashed vertical line separates cross-environment stable (∼60%, left) and unstable (right) EC inputs. Bottom: CA3 input streams indicated by colors and labels (∼60% cross-environment stable, “Task”; see Methods). **(C)** Cascade model of CA3-CA1 synaptic consolidation. W(^0^) (red): synaptic weight modified by both plasticity (BTSP) and consolidation. W(^1^), … W^(*n*)^ (black): hidden consolidation variables. *g*(^1^), … *g*^(*n*)^: transition rates between states. **(D)** Simulated CA1 cell fraction versus PF position on day 1 (novel) for stable (top) and remapped (bottom) cells. Solid: mean; shading: ± SEM, n = 16 simulations; dashed: experimental data. **(E)** Residual activity stability (c.c. with day 0 PF) versus BTSP stability (plateau position difference with day 0 PF) in example simulation (day 1, novel). R = 0.32, p = 1.6e–14, n = 531 cells. Each dot: one cell. p < 0.05 for all 16 simulations. **(F)** Novel environment residual stability (c.c. with familiar environment PF) for high and low (see Methods) training stability cells under control (black, stable synapses (syn.)) and impaired CA3-CA1 synaptic stability (red, unstable syn.). Stable: high 0.56±0.018 versus low 0.38±0.0071, p = 2.0e–7 (“***”); Unstable: high 0.13±0.015 versus low 0.090±0.0029, p = 0.025 (“*”); paired two-sided *t*-test, n = 16 simulations. Circles: individual simulations; crosses: mean±SEM. **(G)** Cross-environment stable cell fraction for control (black, “cntrl”: 0.44±0.0072) and manipulations: impaired CA3-CA1 synapse stability (red, “syn”: 0.27±0.0061, p = 8.2e–18); removed task-relevant CA3 input (orange, “task”: 0.26±0.0042, p = 4.8e–20); removed stable EC input (blue, “EC”: 0.42±0.0083, p = 0.027); all three (green, “all”: 0.22±0.0063, p = 1.3e–20). Removal of cascade (see Methods): 0.33±0.0050, p = 1.3e–13. All versus control, unpaired two-sample one-tailed *t*-test, n = 16 simulations.

The model also revealed how prior learning can bias new plasticity to find generalizable features. Following the transition to a simulated novel environment, model CA1 neurons exhibited residual activity or subthreshold synaptic current before plateau initiation (fig. S9G, top; see also S9H, top (no residual); Fig. 3E and fig. S7 for real neurons). As found in the experimental data, BTSP stability was correlated with residual stability (Fig. 4E). Moreover, neurons with higher levels of synapse consolidation had residuals that were more co-localized with previous days’ PFs (Fig. 4F, black) and displayed stronger cross-environment PF generalization (fig. S9I). Consistent with this, removal of synapse stability significantly reduced residual stability and amplitude (Fig. 4F, red; fig. S9J) and PF generalization (Fig. 4G, red). These results indicate that residuals were supported, at least partially, by CA3 synaptic inputs that had been stabilized during past experience and that this memory component contributes to the generalization process through its influence on plateau initiation and thus BTSP induction (Fig. 4A).

To examine which specific CA3 inputs composed the memory trace present in the residual we systematically manipulated the size of the only consistent cross-environment CA3 input stream (task related inputs; Fig. 4B) and found that this proportionally modulated CA1 field stability (fig. S9K, black) and removing synapse stability in these inputs eliminated this relationship and the ability of the model to generalize (Fig. 4G, orange; fig. S9K, red). The model also showed that distal entorhinal input stability significantly correlated with BTSP stability (fig. S9G, bottom; S9H, bottom; S9L) and elimination of the entorhinal activity stability also significantly reduced generalization although the effect was small (Fig. 4G, blue). This observation is consistent with studies demonstrating that coordinated or matching CA3 and entorhinal input is the primary determinant of plateau potential generation in CA1(*41*, *44*, *45*) (Fig. 4G, green). Altogether the model indicates that a neuronal network implementing BTSP can generalize if there are common elements among the experiences to be found and used to generate a new but recognizable representation. The model suggests that BTSP can perform this function because it is driven by a dendritic computation that responds to matches between an entorhinal input and memory traces stored in the synaptic weights of particular CA3 inputs. This property of BTSP allows past learning to influence new learning.

### Prior experience and local CA1 consolidation history contribute to generalization

The above results and the network model together predicted that robust cross-environment generalization should require substantial prior training and that disrupting local CA1 consolidation history during training should impair later generalization. To test the first prediction, we exposed a separate cohort of mice to the novel environment (belt B) after minimal pretraining in the familiar environment (naïve group; 50±0 laps, over 1 day on belt A, n = 4 mice; Fig. 5A), and compared them with trained mice from the main dataset (expert group; 718±31 laps, over 5.7±0.3 days on belt A, n = 6 mice; Fig. 5A). Naïve mice had weaker anticipatory slowing on both belts (fig. S10A and S10C; running profiles in fig. S10B and S10D). Lick selectivity was lower in naïve than expert mice on the familiar belt (fig. S10E; lick profiles in fig. S10F) but comparable on the novel belt where naïve mice licked primarily after rather than before reward delivery (fig. S10G and S10H). At the cellular level, naïve mice had markedly reduced cross-environment PF similarity than expert mice (Fig. 5B; fig. S10I). The strong relationship between running similarity and PF similarity observed in expert mice was weak in naïve mice (Fig. 5C). This was unlikely to result simply from the narrower range of running similarity in naïve mice, as the relationship remained robust after restricting the analysis to the range of running similarity sampled by naïve mice (fig. S10J). Cross-environment decoding error was correspondingly higher in naïve mice (Fig. 5D). Together, these data indicate that robust cross-environment representational stability depends on substantial prior training.

**Fig. 5.**
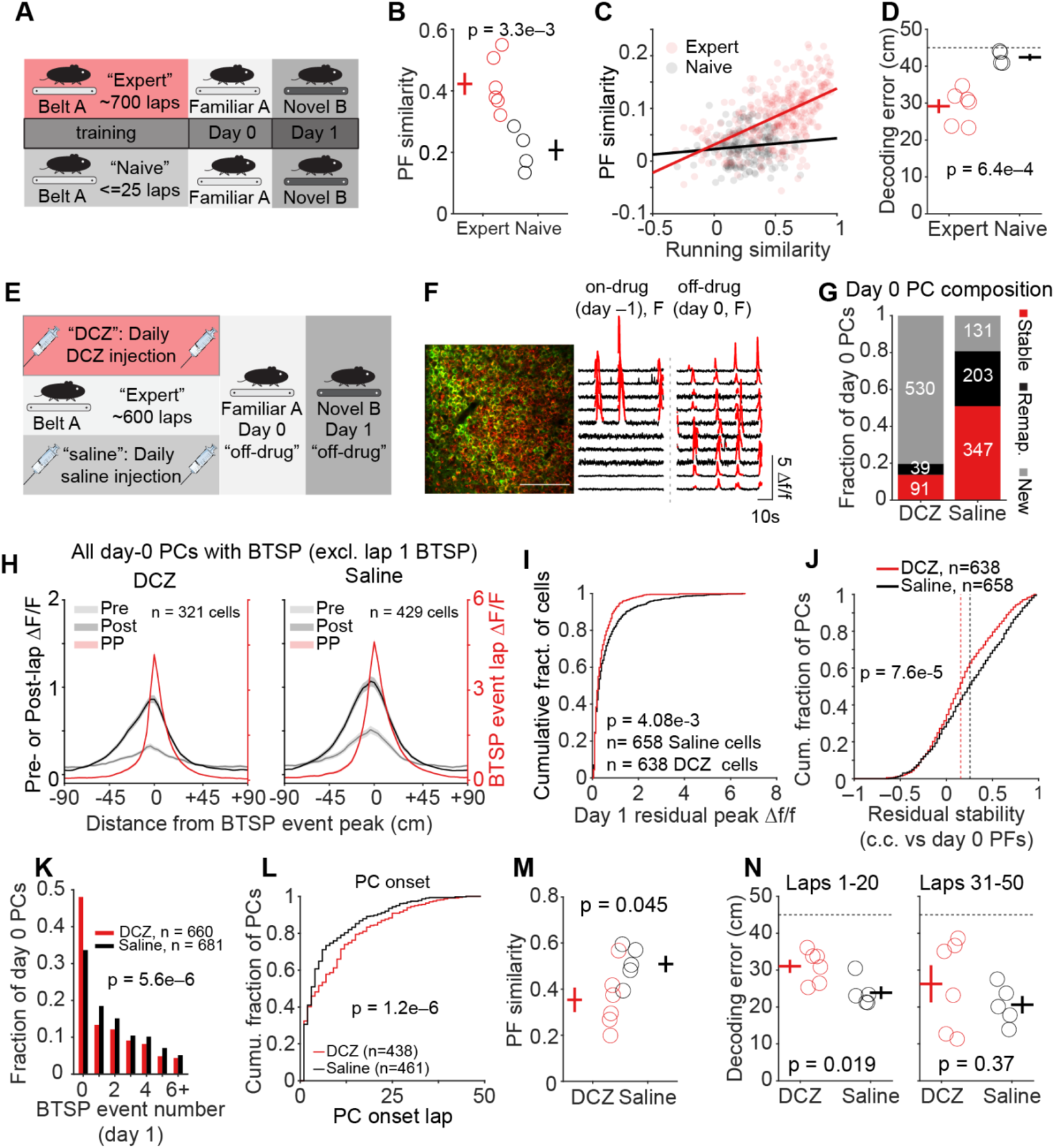
Prior experience and local CA1 consolidation history contribute to generalization. **(A)** Experimental design. Top: expert mice received extensive training on belt A (∼700 laps over ∼6 days) before exposure to familiar env. A (day 0) and novel env. B (day 1). Bottom: naïve mice received minimal training (≤25 laps, 1 day on belt A) before the same familiar-to-novel transition. **(B)** Cross-environment PF similarity (Pearson’s c.c. between day 0 and day 1 mean PFs) for expert (red, n = 6 mice) and naïve (black, n = 4 mice) groups using cells that have reliable PFs on both day 0 and day 1. Expert: 0.42±0.036; naïve: 0.21±0.034; p = 3.3e–3, unpaired *t*-test. Circles: individual mice; crosses: mean±s.e.m. **(C)** Trial-by-trial PF similarity versus running profile similarity (both in c.c. versus day 0 familiar env. template) for expert (red) and naïve (black) mice. Each dot: one lap from one mouse (first 50 laps). Lines: linear fit. Expert: R = 0.58, p = 4.5e–28; naïve: R = 0.12, p = 0.097. **(D)** Cross-environment decoding error for expert (red, 29.2±1.9 cm, n = 6 mice) and naïve (black, 42.4±0.88 cm, n = 4 mice) groups. p = 6.4e–4, unpaired *t*-test. Dashed line: chance level. Circles: individual mice; crosses: mean±s.e.m. **(E)** DREADDs experimental design. GP5.17;CaMKII-Cre mice co-expressing GCaMP6f and Cre-dependent Gi-DREADDs in CA1 pyramidal neurons received daily DCZ (n = 6 mice) or saline (n = 5 mice) injections throughout training (∼600 laps on belt A). Three imaging sessions: day –1 (belt A, on-drug), day 0 (belt A, off-drug), and day 1 (belt B, off-drug). Three additional DCZ mice were excluded due to insufficient silencing (fig. S11). **(F)** Representative two-photon field of view from a DCZ mouse. Left: transgenic expression of GCaMP6f (green) and viral expression of mCherry (red), showing dense Gi-DREADDs expression. Right: example Δf/f calcium traces on day –1 (on-drug) and day 0 (off-drug). Significant events are plotted in red. Note a small fraction of reactivated cells that escaped silencing even with DCZ. Scale bar: 100 μm; **(G)** Stacked bar showing composition of day 0 PCs by day –1 history for DCZ and saline groups. Cells that were reactivated (having PFs on both day -1 and day 0) were further split into stable (red): PCs on both day –1 and day 0 with stable PF (DCZ: 91; saline: 347) and remapped (black) cells: PCs on both days but PF changed more than 15 cm (DCZ: 39; saline: 203). New (grey): PC on day 0 but not day –1 (DCZ: 530; saline: 131). Numbers inside bars: cell counts. **(H)** Population-averaged day 1 spatial tuning aligned to each cell’s first BTSP event peak (position 0) for all day 0 PCs with detectable BTSP event on day 1 (excluding first-lap BTSP) for DCZ mice (Left, n = 321 cells in 6 mice) and saline controls (Right, n = 429 cells, 5 mice). For each cell, the mean Δf/f profile from the 5 laps preceding the first BTSP event (Pre, grey, left axis), the 5 laps following (Post, black, left axis), and the BTSP event lap (red, right axis) were circularly shifted so that the BTSP event peak location was centered. Lines and shading: mean±sem. **(I)** Cumulative fraction of cells versus day 1 residual peak Δf/f (maximum of the day 1 pre-BTSP residual within ±12 bins (≈43 cm) of each cell’s day 0 PF peak). All day 0 PCs, first-lap BTSP excluded. Saline (black, n = 658 cells, 5 mice); DCZ (red, n = 638 cells, 6 mice). p = 4.08e–3, two-sample Kolmogorov-Smirnov test. (H) included only day 0 PCs with a detected BTSP event on day 1, so Pre, Post, and BTSP event lap can be aligned to the BTSP event; (I) included all day 0 PCs, whether or not these cells had PFs on day 1, whether or not they had BTSP on day 1. For cells without detected BTSP event, a pre-BTSP window matched to the typical first-BTSP lap was used to produce the residual (see Methods). **(J)** Cumulative distribution of residual stability (c.c. between pre-BTSP spatial activity on day 1 and day 0 PF) for DCZ (red, n = 638 cells) and saline (black, n = 658 cells). p = 7.6e–5, two-sample Kolmogorov-Smirnov test. Dashed vertical lines: median for each group. **(K)** Fraction of day 0 PCs as a function of number of detected BTSP events on day 1 for DCZ (red, n = 660 cells, 6 mice) and saline (black, n = 681 cells, 5 mice) groups. p = 5.6e–6, two-sided Wilcoxon rank-sum test. **(L)** Cumulative distribution of day 1 PC onset lap for DCZ (red, n = 438 cells, 6 mice) and saline (black, n = 461 cells, 5 mice), using cells that are PCs on both day 0 and day 1 with a detectable day 1 PC onset. p = 1.2e–6, two-sample Kolmogorov-Smirnov test. **(M)** Cross-environment PF similarity (c.c. between day 0 and day 1 mean PFs, using all day 0 PCs) for DCZ (red, 0.35±0.053, n = 6 mice) and saline (black, 0.51±0.035, n = 5 mice) groups. p = 0.045, unpaired *t*-test. Circles: individual mice; crosses: mean±s.e.m. **(N)** Cross-environment decoding error (cm) for DCZ (red) and saline (black) during early (laps 1– 20, left) and late (laps 31–50, right) exposure to novel env. B. Laps 1–20: DCZ 31.0±1.8 versus saline 23.9±1.7cm, p = 0.019; laps 31–50: DCZ 26.3±5.0 versus saline 20.6±2.4, p = 0.37; unpaired *t*-test. Dashed line: chance level. Circles: individual mice; crosses: mean±s.e.m. n = 6 DCZ, 5 saline mice.

We next tested whether CA1 generalization requires local CA1 consolidation history rather than being inherited entirely from upstream inputs. We used inhibitory chemogenetic Gi-DREADDs (designer receptors exclusively activated by designer drug) to suppress local CA1 activity throughout training. We injected AAV-flex-Gi-mCherry into GP5.17;CaMKII-Cre mice to co-express GCaMP6f and Cre-recombinase-dependent Gi-DREADDs in a subset of excitatory CA1 neurons from which we longitudinally imaged during training (Fig. 5E and 5F). This manipulation was designed to disrupt the activity-dependent consolidation history locally in CA1 without directly suppressing upstream CA3 or EC inputs.

Mice received daily deschloroclozapine (DCZ; n = 6 mice) or saline (n = 5 mice) treatment throughout ∼5 days of training in the familiar environment. Three additional DCZ mice were excluded from the main analyses due to insufficient silencing (fig. S11; see Methods). Longitudinal calcium imaging was performed on the final drug day (day –1, belt A, familiar), the first off-drug day (day 0, belt A, familiar), and the novel-environment day (day 1, belt B, off-drug). Both groups showed robust anticipatory slowing and selective licking across all three testing days (fig. S12A to S12F), indicating that local CA1 silencing had minimal impact on task-relevant behavior. DCZ effectively reduced the PC fraction in CA1 by 77% on day –1 (saline: 0.57 ± 0.059; DCZ: 0.13±0.022; fig. S12I). On drug-off days 0 and 1, the PC fraction in DCZ mice recovered to levels comparable to saline controls (fig. S12J and S12K). Consequently, DCZ mice contained a greatly expanded population of day 0 PCs that had not been detected as PCs on day –1 (“new” PCs; DCZ: 530/660 versus saline: 131/681; Fig. 5G; fig. S12G, S12H). These new PCs had markedly later PF onset on day 0 than that of reactivated cells that escaped DREADD silencing on day –1, consistent with *de novo* PF formation on day 0 after removal of DREADD-mediated suppression (fig. S12L).

We next asked whether disrupting CA1 activity during training impaired cross-environment residual activity, BTSP induction and population activity level generalization. Indeed, DCZ mice exhibited significantly reduced residual activity amplitudes (Fig. 5H, 5I) and correlation stability relative to day-0 PFs (Fig. 5J), whereas residual peak displacement among cells with detectable residual activity was unchanged (fig. S13A). Consistent with the biasing role of residual activity, BTSP recruitment was significantly reduced in DCZ mice (Fig. 5K), as was the amplitude of BTSP-lap activity aligned to the day-0 PF (fig. S13B), although among cells exhibiting BTSP, neither the timing of the first BTSP event nor BTSP peak displacement relative to the day-0 PF differed significantly between groups (fig. S13C, S13D). In addition, PC onset in the novel environment was also delayed in DCZ mice (Fig. 5L). Among PCs present on both day 0 and day 1, cells without detectable day 1 BTSP still developed spatially localized activity after PC onset, but at substantially lower amplitude than BTSP-positive cells in both groups (fig. S13E to S13J; see fig. S13K for a summary). Finally, DCZ mice had lower cross-environment PF similarity on day 1 than saline mice (Fig. 5M), and cross-environment decoding error was elevated during the first 20 laps in the novel environment (Fig. 5N) but was comparable between groups during later laps (laps 31-50; Fig. 5N). We also observed that neither the fraction of stable PCs nor cross-environment PV similarity differed significantly between groups (fig. S13L, S13M) and suspect that these measures are relatively insensitive to the population of neurons missing BTSP that nevertheless exhibit small amplitude spatially tuned activity reminiscent of residual activity.

Because the day 0 DCZ population contained both reactivated PCs (20%) that escaped silencing and newly emerged PCs (80%), we next asked whether these populations were differentially affected. Reactivated DCZ PCs showed cross-environment PF similarity comparable to saline controls (fig. S13N), whereas new PCs had lower PF similarity than reactivated PCs in both treatment groups (fig. S13O, S13P). Residual stability was likewise preserved in reactivated DCZ PCs but reduced in new DCZ PCs (fig. S13Q, S13R), indicating that the impairment was primarily in the new cells. Consistent with these single-cell analyses, novel-environment population activity showed weaker alignment with the familiar-environment subspace when the analysis was restricted to new DCZ PCs (fig. S14A to S14C), whereas subspace alignment across all PCs did not differ significantly between groups (fig. S14D). New DCZ PCs nevertheless retained measurable cross-environment generalization (fig. S13O, S13P). Together, these results demonstrate that local CA1 consolidation history supports the strength of residual activity, BTSP recruitment, and cross-environment generalization.

## Discussion

Above we report, 1) that during the exploration of novel environments a preserved CA1 population activity, residing in a sub-space shared with population activity recorded in familiar environments, was actively generated. These data, which are consistent with previous studies(*6*, *25*, *26*), confirm CA1 possesses mechanisms that allow it to produce generalized population activity. 2) As we have previously reported, PF activity recorded in novel environments exists almost exclusively in a goal reference frame(*7*), and this is true here even for PCs that maintained the same PF location as in the familiar environment. This could be because PCs in our novel environments are encoding the only information that is common to both experiences (task structure or run 180 cm for reward). This further suggests that this shared structure may have been involved in producing the population level activity that exists in the shared sub-space. 3) We found evidence that BTSP is heavily involved in producing PF activity in the novel environment. Further we observed the presence of weak “residual activity” preceding BTSP induction of new robust PFs. In the PCs generating the preserved activity these residuals were found near the location of the PFs recorded during both the previous and current experience, even when the current experience was of a novel environment. We and others(*48*) repeatedly find evidence that CA1 PC representations are rebuilt at the start of each experience through an active, BTSP mediated, process. Residual activity is a prominent feature of this rebuilding process and is thought to reflect components of PFs that are more stable (hypothesized to be consolidated synaptic weights(*43*, *48*)) thus allowing them to be present in future experiences that contain some element common with past experiences. 4) We constructed a computational model that featured: multiple synaptic inputs using different reference frames, BTSP mechanisms to adjust synaptic weights and a synaptic consolidation mechanism that was also driven by plateau potentials. This model reproduced our experimental results and predicted that residuals generated from stabilized synaptic weights affiliated with CA3 inputs encoding task structure biased which CA1 cells would express place fields and at what location to build the recognizable PC activity that supports generalization. This represents a core mechanism whereby CA1 can extract, maintain and use the shared structure present in both the familiar and novel environments. 5) We tested the model predictions with two manipulations. First, a simple behavioral manipulation that reduced the amount of experience available to the network to consolidate any relevant aspects weakened cross-environment neural generalization and its coupling to behavior. Second, we used chemogenetic methods to suppress local CA1 network activity during learning to reduce the amount of activity-dependent consolidation present within the recorded network while leaving behavior intact. Following activity suppression, we observed residual activity, BTSP induction, PF formation and network level measures of generalization to all be impaired. These results add weight to the concept that an experience- and activity-dependent consolidation process occurs during learning episodes that can later influence new learning if there are enough shared aspects of the experiences to activate the appropriate input. In the end the above results suggest a cellular/synaptic mechanism for generalization where a given network extracts from its own activity the structure common to multiple experiences and uses such structure to rapidly generate population activity residing within a shared sub-space to speed learning of novel environments.

There are two caveats concerning the above scheme that should be noted. First the synaptic current present within residual activity is but one element involved in determining the probability of plateau potential initiation in pyramidal neurons which could explain the observation that residual activity is more of a biasing function as opposed to a pure instructive signal(*43*). This does leave a potential role for the other elements controlling plateau initiation (e.g. EC3 excitatory input(*41*, *44*, *45*) and feedback inhibition(*58*, *59*)) to be explored in subsequent experiments on CA1 generalization. Second, we have used a computational model to explore the capabilities of plateau potential dependent synapse consolidation or weight stabilization as a potential mechanism for the extraction of shared structure among various experiences. Although our experimental data are consistent with the predictions produced by a model implementing such a mechanism we have no direct evidence, yet, that synapses are involved. Indeed, such evidence awaits the discovery of the molecular mechanisms of synapse consolidation (aka memory).

It is worth distinguishing the reference frame of a place field from the generalization described here. Referencing concerns what a field is anchored to and where the origin of that anchoring lies; it is a property of the representation that can be read within a single session and that can be altered rapidly, within a few trials, by a shift in the origin alone(*7*). Generalization concerns whether a recognizable representation is rebuilt when the sensory world changes; it is assessed across sessions and against behavior, and it depends on structure accumulated over previous days. The two are nonetheless linked, in that the reference frames a cell receives determine which stored synaptic structure is available to bias plateau initiation in a new environment. Because the sensory cues were changed completely here, the goal frame was the only frame carrying structure shared between the two experiences, and generalization was accordingly carried by goal-referenced inputs. Which frame carries generalization is thus a consequence of what recurs across experiences rather than a privileged property of the goal frame itself. Viewed this way, the available reference frames constitute a small set of candidate organizing variables, a compressed description of the ways a current experience might be structured, among which consolidation selects by retaining whatever recurs.

Individual CA1 PCs receive an experience-dependent mixture of differently referenced synaptic inputs(*7*), any of which they can use to form new representations. The potential advantage this provides to a network is that it alleviates the need to determine, a priori, which aspects of experience will be important for guiding learning in the future. Instead, the network simply stores any structure that recurs consistently across experience. It may turn out that this simple mechanism could allow any individual network to extract shared structure from its own activity online during behavior. Indeed, the possibility that every network implementing BTSP and experience-dependent information consolidation mechanisms is capable of generalization, raises the question as to how much and what type of generalization may be present in CA3 and EC3. However, our CA1 activity manipulation suggests much of the generalization we observe in CA1 is built there locally, but future experiments are needed to explore similar phenomena in other brain regions, including CA3, EC, prefrontal cortex and other neocortical areas.

Where do our current experiments fit into long discussed ideas about orthogonalization, generalization and PF remapping in the hippocampus(*60–62*)? As we see it, global remapping or orthogonalization(*17*, *53*, *63*, *64*) occurs when there is insufficient common structure among multiple experiences or if, somehow, the consolidation of existing structure is impeded. Partial remapping, where rate remapping is the limit, is thought to occur when there are substantial similarities between different environments that are also, perhaps, experienced over short timeframes (within the same session or day). It is noteworthy that many groups find there to be a significant amount of CA1 PC remapping or representational change that occurs even when familiar environments are repeatedly experienced over sequential days(*48*, *65–67*), and that the amount of representational similarity increases with experience(*48*). As discussed above, that we observed a similarly moderate amount of remapping here for experiences of both familiar and unfamiliar sensory environments suggests that there was enough common structure between these experiences to allow a recognizable representation to be formed (i.e. activity in the shared subspace). Within this framework then, the generalizations we found are likely to be closest to a partial remapping that was more substantial than simple rate remapping because the appropriate representations were actively rebuilt, with enough common structure residing in the goal referenced PFs to aid in this rebuilding process.

Generalization is a critical function observed across multiple brain regions(*21*, *25*, *35*, *68*, *69*) and this process requires extracting common elements from prior experience for use in novel conditions. Here we present evidence that a network implementing BTSP and an information consolidation mechanism (perhaps synaptic weight stabilization) can extract shared structure and use it to direct new learning that rapidly reforms recognizable population activity, thus mediating generalization. Although demonstrated here in a relatively simple spatial task, this memory-trace-guided rapid learning may represent a core principle for how brains use prior knowledge to adapt to complex, novel situations.

## Acknowledgments

We thank R. Chitwood for technical support. We thank C. Kemere for comments on the manuscript and helpful discussions. We thank J. Hennig and X. Chen and Magee lab members for useful discussions.

## Author contributions

F.K.Q and J.C.M were responsible for conceptualization. F.K.Q performed in vivo recordings and analyzed the data. G.L designed and implemented the computational model and analyzed the simulated data with inputs from F.K.Q, J.C.M. D.L provided guidance for the subspace analysis, and S.R provided guidance for the computational modeling. F.K.Q and J.C.M wrote the paper with feedback from all authors.

## Data and materials availability

The CA1 data supporting this study’s findings will be freely available upon publication. The CA3 axon data are available from the corresponding author upon request. Scripts that support the experimental data results of this study will be freely available via GitHub upon publication.

Simulation scripts are available from the corresponding author upon request.

## Materials and Methods

All experimental procedures performed were approved by the Baylor College of Medicine Institutional Animal Care and Use Committee (protocol AN-7734).

### Datasets

The longitudinal CA1 imaging dataset from Qian et al. (2025)(*1*) was reanalyzed to examine cross-environment generalization, rather than the within-environment reference frame dynamics for which it was originally collected (Figs. 1–3 and related supplementary figures, also the expert group in Figs 5A to 5D and related supplementary figures). This reanalysis included analyses of behavioral transfer and place-field stability across environments, Bayesian decoding, principal component analysis (PCA), residual activity preceding BTSP, and reference-frame switching. We also analyzed previously collected behavioral data from naïve and trained mice that had not been analyzed for this purpose (Figure 1B to 1F). Data collected for the present study included CA3 axon recordings (fig. S5), naïve-control experiments, and Gi-DREADD perturbation experiments testing how prior experience and local consolidation history contribute to generalization (Fig. 5; fig. S10 to S14).

### Surgery

Surgical procedures followed previously published methods(*1*). Experiments were performed in adult GP5.17 (*2*) (n = 15 mice) or Grik4-cre mice(*3*) (n = 10 mice) or GP5.17;CaMKII-Cre crosses (n = 14 mice for the DREADD experiments; n = 4 mice for naïve controls (Fig.5)) ≥8 weeks old at surgery, of either sex, targeting the right hemisphere. Animals were housed under an inverse dark-light cycle (dark: 9:00-21:00). All recordings occurred during the dark phase. Mice were anesthetized with ∼2% isoflurane for surgeries as previously described(*1*, *4*). Following local anesthetics application, a section of flap was removed, and the skull cleaned and leveled. A 3.0-mm-diameter craniotomy centered at 2.3 mm posterior to bregma and 2.15 mm lateral to midline was marked for both CA1 soma or CA3 axon imaging. The dura was removed using forceps (Fine Science Tools), and overlying cortical tissue was gently aspirated using a blunt needle (McMaster-Carr) under repeated saline irrigation. Aspiration exposed the external capsule, which was lightly peeled without direct needle contact. Irrigation continued until the bleeding ceased. A cannula (3-mm diameter, 1.75–2.2-mm long) with a coverslip (Potomac, 2.90-mm diameter, #0) attached using UV-curable optical adhesive (Norland Products) was inserted and secured with dental cement (Ortho-Jet, Lang Dental). A custom titanium head bar was cemented to the skull parallel to the coverslip surface for optimal imaging quality.

### Virus injection

Virus injection for CA3 axon imaging was adapted from previous studies(*5*). A mixture of AAV1-hSyn-Flex-jGCamp8m-WPRE (∼2e12 GC/mL) and AAV1-hSyn-tdTomato-WPRE (∼5e11 GC/mL) injected at 2.2 mm posterior to bregma and 2.25 mm lateral to the midline. Three injections (∼50nL per depth) were made at 1.9, 1.75, and 1.6mm (dorsoventral) below dura, followed by a 10-min wait with the pipette positioned above the final injection depth. The injection system comprised a pulled glass pipette (broken and beveled to ∼30 μm (outer diameter); Drummond, Wiretrol II Capillary Microdispenser) backfilled with mineral oil (Sigma). A fitted plunger advanced by a manipulator (Drummond Nanoject II) displaced the pipette contents, while plunger retraction loaded virus. The injection pipette was positioned using a Sutter MP-285 manipulator.

For DREADD experiments, AAV1-hSyn-Flex-hM4D(Gi)-mCherry (Addgene: 44362-AAV1) was injected into dorsal CA1 using the same injection system. Two cohorts differed in viral titer and injection volume. In both cohorts, injection targeted two sites in the right hemisphere (2.4 mm posterior and 2.3 mm lateral; 2.0 mm posterior and 1.8 mm lateral), with three injections per site at 1.2, 1.05, and 0.9 mm below the dura. Cohort 1 received virus at a final titer of 2.4e12 GC/mL (∼25 nL per depth) and included 5 DCZ-treated mice (two weakly and three strongly silenced) and three controls. One control was Cre-negative, received DCZ and was pooled with the other controls. Cohort 2 received virus at a final titer of 8e12 GC/mL (∼50nL/depth) and included 4 DCZ-treated mice (one weakly and three strongly silenced) and two controls. Imaging started after 3-4 weeks of viral expression and this was comparable between groups (DCZ, 25.5±1.5 days; Control, 26±1.1days; measured on day 0 on belt A). Weak silencing occurred in both cohorts, but in a smaller fraction of cohort 2 (1/4) than cohort 1(2/5).

### Behavior task

Behavioral procedures were described previously(*1*, *6*). Briefly, the linear treadmill system featured a ∼180-cm velvet fabric belt (McMaster-Carr). Water-restricted mice were head-fixed to custom stainless-steel posts and self-propelled the belt. Movement speed was measured by a rotary encoder using Arduino-based microcontrollers. The digitized speed signal interfaced with a behavior control system (Bpod r0.9–1.0, Sanworks) via custom MATLAB code (2019b, MathWorks) on a Windows PC. Reward delivery (10% sucrose in water) was controlled by a solenoid valve (The Lee Co.) through a custom lick port, with licking detected by an optical sensor (FX-300 series, Panasonic). The Bpod system interfaced with the rotary encoder, solenoid valve and lick sensor. Behavior data were digitized through a PCIe-6343 X-series DAQ system (National Instruments) running WaveSurfer software (v.0.982, Janelia Research Campus).

After ≥6 days of recovery from surgery, mice were placed on water restriction (1.5 ml/day) and handled daily (10–30 min) for ≥4 days. Mice were then introduced to the treadmill system, running on a tactile-cue-featured belt, receiving water at epoch-dependent increasing distances (8– 15 laps per epoch). Reward distances were fixed within epochs but increased between epochs. Manual rewards were occasionally provided to encourage running. Once the reward distance reached 126 cm, a single reward was delivered per full belt lap at a fixed location (50 cm).

Fifteen mice underwent behavioral training (fig. S1A). Mice were classified as “naive” if they completed ≥50 laps on training belt with <25 total previous full laps on the belt (n = 10 mice qualified; 5 did not but continued in the study). All 15 mice then received ∼6 days of training on belt A to establish familiarity with the task structure and tactile environment. Following training, mice were assigned to one of two conditions: “cross-environment” (cross-env., n = 6 mice) introduced belt B, which featured distinct tactile cues from belt A, with the reward-switch task, testing generalization across sensory contexts with identical task structure; “same-environment” (same-env., n = 9 mice) introduced the reward-switch task on the familiar belt A.

Day 0 was defined as the final training session on the familiar belt A, with rewards at 50 cm. The reward location was switched on Day 1: rewards were delivered at 50 cm for the first 100 laps, then 140 cm for the next 100 laps. Sessions typically lasted 30–60 min, with one session per day. Belt A featured three uniform tactile cues (Velcro, glue stick and white fabric; 60 cm per cue zone) and belt B featured six distinct cues (30 cm per cue zone). For CA3 axon imaging, animals were trained similarly as CA1 dataset, except that we reduced the threshold to 100 laps per session for final training days. CA3 analyses were limited to 50 laps before (pre) and after (post) the reward shift. Initial reward-switch behavior was quantified over the first 20 post-shift laps.

Naïve mice. To test whether cross-environment generalization required prior experience, five mice received minimal training before the belt switch and were treated as naïve controls. For this cohort, day 0 was the first session in which a mouse completed ≥50 laps. Lap and day counts began with the first session in which reward was fixed on the belt. To control both cumulative lap number and training days, mice were included as naïve controls only if they had completed ≤25 training laps and <2 training days before day 0 on belt A. Four mice met these criteria. Each completed 50 laps during its first exposure to belt A with reward fixed at 50 cm (day 0) and therefore had no prior laps. One mouse completed <50 laps/session for three consecutive days and was excluded. On the following day (day 1), the four included mice were introduced to a novel belt B and completed 50 laps with reward at the same location (50 cm). The session ended after 50 laps, and no reward switch was performed.

DREADD experiments. The chemogenetic experiments used the same behavioral paradigm. Deschloroclozapine dihydrochloride (DCZ; MedChemExpress) was used to activate the inhibitory DREADD receptor hM4D(Gi). DCZ was selected instead of Clozapine N-oxide (CNO) because DCZ activates DREADD receptors more effectively at far lower doses(*7*, *8*) and avoids the slower kinetics and off-target effects associated with CNO(*9*). The application of DCZ followed previously published methods(*10*). DCZ was dissolved in DMSO at 10 mM (∼3mg/mL) and stored at –20℃ in 10-uL aliquots. On each experimental day, one aliquot was thawed at room temperature and diluted in saline to 0.04 mg/mL. Mice received an intraperitoneal injection of 0.05 to 0.1 mL of this solution (0.1 mg/kg) 15 min before imaging. Control mice received an equal volume of saline on the same schedule. DCZ was administered on every training day on belt A, from the first exposure through the final training day (day –1), when mice met the well-trained criterion. On the following day (day 0), mice performed the same task on belt A with no injection, allowing new PFs to form in the absence of silencing. Day 1 on the novel belt B was also injection-free. Only the first 50 laps on belt B were analyzed. DCZ and saline groups were matched for cumulative training laps (DCZ, 532±41 laps; Saline, 569±33laps; p = 0.51, unpaired t-test) and training days on belt A (DCZ, 5.8±0.6 days; Saline, 4.8±0.8 days; p = 0.32, unpaired t-test).

### Two-photon imaging

All two-photon Ca2+ imaging recordings were performed in the darkness using a custom-made microscope (Janelia MIMMS 2.0). GCaMP expressed in hippocampal pyramidal neurons was excited at 920 nm by a Ti:Sapphire laser (Chameleon Ultra II, Coherent; typically 30–50 mW measured after the objective) and fluorescence was collected through a Nikon ×16, 0.8 numerical aperture objective. Emitted light passed through a 565 DCXR dichroic filter (Chroma) and either a 531/46 nm (green channel, Semrock) or a 612/69 nm (red channel, Semrock) filter before detection by a GaAsP photomultiplier tube (11706P-40SEL, Hamamatsu). Images (512 × 512 pixels) were acquired at ∼30 Hz using ScanImage (Vidrio Technologies). For CA1 soma imaging, fields of view (FOVs) (∼300 × 300 µm) were selected based on visible somatic Ca^2+^ transients. A reference FOV was registered daily for longitudinal tracking. Experiments were terminated if substantial FOV changes occurred between sessions. For CA3 axon imaging, FOVs (∼250um by 250um) were selected based on fiber morphology in the red channel and occasional Ca^2+^ activity. No attempt was made to track the same FOV across days. For DREADD CA1 soma imaging, each FOV was first illuminated at 1000 nm for ∼15s while the belt was held stationary to assess mCherry expression. Imaging was then switched to 920 nm for functional GCaMP6f recording. FOVs in chemogenetic experiments (∼350um by 350um) were selected to include only neurons with clear mCherry expression. The same criterion was applied to DCZ and control groups.

### Data analysis

#### Pre-processing and signal processing

CA1 data were analyzed as described previously (*1*) with modifications. Two-photon images were motion-corrected with Suite2p(*11*) (Python version; http://github.com/MouseLand/suite2p). For longitudinal imaging, data acquired each day were concatenated before motion correction. Regions of interest (ROIs) were automatically detected and time series fluorescence traces were extracted using Suite2p. Manual curation discarded unwanted ROIs and reselected ROIs rejected by Suite2p based on anatomical features in the Suite2p interface. ROI contamination was examined manually as previously described(*1*). No neuropil subtraction was applied. Further analyses used custom MATLAB code. Raw fluorescence signals were baseline-corrected and converted to Δf/f ((F – F_0_)/F_0_), where F_0_ was the histogram mode of F. Signals were mean-smoothed using 0.2-s sliding window (6 frames). Significant calcium transients were defined as events exceeding 3 standard deviations above baseline noise, estimated from deviations below the mode of Δf/f values. Spatial maps of Δf/f were generated using frames where running speed exceeded 5 cm/s. The 180-cm belt was uniformly divided into 50 bins (3.6 cm/bin), and mean Δf/f was calculated per spatial bin (excluding frames with speed <5cm/s). We note that main conclusions are still valid if we excluded frames without significant calcium transients before calculating mean Δf/f per spatial bin (fig. S8). Gaussian smoothing (3 bins) was applied for display only. Only the first 100 laps before and after reward switch were analyzed (all available laps were used if <100). Reward locations were at bin 14 (50 cm) pre-switch and bin 39 (140 cm) post-switch. Day 0 sessions typically had ≥100 laps (50 laps for naïve mice). Reward locations are indicated in all figures.

#### Place cell identification

Place cell (PC) identification was based on each cell’s spatial Δf/f map. PC onset was defined as the first lap for which the current lap and ≥2 of the next five laps each contained ≥3 consecutive spatial bins with at least one frame of significant calcium (activity >3×SD above baseline), detected within ±45 cm (90 cm total) of the place field (PF) peak bin. PFs were defined as the trial-averaged Δf/f activity vectors across 50 spatial bins (50 by 1 vectors). Only the first onset lap was identified, separately for each condition (pre- and post-reward switch). Laps following the onset lap were used for PC classification.

A cell was spatially modulated if its trial-averaged PF spatial information (SI) exceeded the 95^th^ percentile of the shuffled SI distributions. SI was calculated as previously described(*12*):

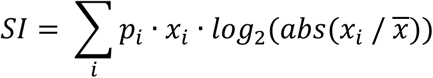

Here, *p_i_* is the occupancy probability of bin *i*, *x_i_* is the smoothed mean activity in bin *i*, and *x* is the overall mean activity. For each shuffle, the Δf/f time-series traces was circularly shifted by ≥500 frames, then divided into six equal segments, and randomly permuted. The shuffled spatial Δf/f map and SI were then re-calculated (200 shuffles). A cell was considered reliable if >20% of post-onset laps had significant calcium events within ±45 cm of the PF peak. PCs were cells whose mean PFs were both spatially modulated and trial-by-trial reliable. Onset was assigned NaN if not detected, and PC identification began from lap 1. No additional processing was performed for cells with multiple PFs.

#### Determination of place-cell types

PC reference frames were classified as described previously (*1*). In reward-switch experiments, reward moved from 50 cm (pre) to 140 cm (post; ΔSwitch = 90 cm). Spatial-cue-referenced (“space”) PCs maintained stable PF peaks relative to belt cues with pre-to-post displacement of 0– 15 cm in the belt coordinates. Reward-goal-referenced (“goal”) PCs shifted their PF peaks by 75– 90 cm in belt coordinates and therefore remained aligned to reward in reward-centered coordinates. For cross-day analyses, the reward location was unchanged from belt A (day 0) to belt B (day 1). PCs with reward-relative peak displacement ≤15 cm between days were classified as stable; all others were classified as remapped. For each animal, the stable cell fraction was the number of stable cells divided by the number of PCs with reliable PFs in both conditions (e.g., environments A and B).

#### Behavioral spatial maps

Behavioral signals were sampled at 10 kHz. Running velocity and lick count maps were generated by binning the 180-cm belt into 50 bins (3.6 cm/bin). Mean velocity (cm/s) and total lick count were then calculated per bin. Lick probability was calculated as the fraction of laps with ≥1 lick per bin. Anticipatory slowing was calculated as the mean speed 30 cm before reward minus mean speed in a 30-cm zone starting 40cm after reward (approximately 30 cm after run start). Lick selectivity was the ratio of licks within ±20cm around reward to total track licks.

#### Post-reward stop and run onset identification (CA3 axon only)

For each lap, a search window was defined ending when speed-integrated distance exceeded 0.5 m or 50 s elapsed, whichever occurred first. Within this window, the stop point was identified hierarchically: (1) the first point where a stationary index #1 (speed < 5 cm/s = 1, else 0; 0.5-s moving average) exceeded 0.9, or 0.5 if 0.9 was absent; (2) if absent, the earliest occurrence of speed < 1 cm/s; (3) if still absent, we allowed < 5 cm/s; (4) if still absent, the minimum speed point within 1 s after reward release. Distance was realigned to zero by subtracting the mean distance over the subsequent 0.1 s around the stop point. To detect run onset, speed was binarized (>1 cm/s = 0, else 1) and a 1.0-s moving average applied to generate stationary index #2. Brief run bouts were removed by detecting stand-run transitions (0.5 cm/s threshold) and zeroing speed within bouts traveling < 8 cm, producing a bout-removed stationary index #2. Run onset was defined as the last time point before 18 cm traveled where the stationary index ≥ 0.5 and its centered trend (future minus past over ±0.5-s window) was < −0.1. If no extended stationary period was detected, speed was smoothed (0.25-s window), the minimum within the first 1 s post-reward identified, and run onset defined as the first time speed exceeded 1.25× this minimum. Ambiguous cases (typically < 10% of laps) were manually examined using custom plots. Spatial maps after reward switch were re-aligned by shifting the map by 90 cm (25 spatial bins).

#### DREADD silencing verification (activity inspection)

Gi-DREADD activation did not silence all labeled CA1 neurons. We therefore evaluated silencing in each DREADD mouse and excluded weakly silenced animals as described below (fig. S11). Because the same FOVs were tracked longitudinally, CA1 activity on the final drug-on training day (day –1) was compared with activity in the same mouse on the drug-free day 0. The same comparison was performed in control mice for reference. Analyses were restricted to running frames (speed≥5 cm/s). For each cell, day –1 activity was the fraction of running frames containing a significant transient. Analyses focused on cells classified as PCs on drug-free day 0, and a successfully silenced PC was expected to have near-zero activity on day –1, whereas an unsuccessfully silenced PC was expected to retain activity near the control level. Because cells were not tracked over earlier training days, each day-0 PC’s day –1 activity served as a proxy for suppression by Gi-DREADD activation. The silencing threshold was determined as the control-group mean minus 2 SD for the day –1 active-frame fraction. DREADD mice below this threshold were classified as successfully silenced; Mice at or above this threshold were classified as weakly, or unsuccessfully, silenced and excluded from subsequent analyses. Six of nine DREADD mice met the successful-silencing criterion. To visually assess whether silencing of individual neurons covaried with mCherry expression, we overlaid each cell’s activity level on the mCherry image. Within AAV-infected regions, neurons that retained high activity and escaped DREADD silencing displayed a salt-and-pepper distribution, without clear clustering around cells with high mCherry expression (fig. S11C). The basis of the incomplete silencing is unknown.

#### CA3 axon analysis

CA3 axon data were analyzed using procedures adapted from previously published methods(*5*, *13*). Unless otherwise specified below, analyses were identical to those used for CA1 somata.

### Signal processing

Raw fluorescence was smoothed with a 0.5-s moving-average window (15 frames) before baseline correction and calculation of Δf/f as described for somata. The signal threshold matched the somatic definition (mean + 3 SD; see *Pre-processing and signal processing*). In addition, a significant transient was required to remain above threshold for ≥7 consecutive frames (∼0.23 s). This seven-frame minimum corresponded to a trough in the pooled event duration distribution and separated genuine transients from brief threshold-crossing noise. The resulting frame-wise event mask defined significant activity in all subsequent analyses.

### Spatial binning

Because mice stopped after reward and sometimes remained stationary for extended periods between laps, CA3 analyses included an additional procedure to identify cells whose activity peaked during standing before run onset or after the post-reward stop. Spatial lap boundaries were anchored to detected run onset and post-reward stop rather than belt position (see *Post-reward stop and run onset identification*). Each lap extended from one run onset to the stop after the next reward and was divided into 50 equal-distance bins, with mean Δf/f computed per bin. While trial-to-trial variation in stopping location produced small differences in lap length, mice typically stopped 10-20 cm after reward, binning directly by belt position did not alter the conclusions. Cells with activity peaked during post-reward standing were excluded from place-cell referencing analyses. Including these cells did not alter the conclusions.

ROI clustering. Because one CA3 neuron could in principle produce many Suite2p axon ROIs, ROIs originating from the same neuron were identified and merged before place-field analysis. For each ROI pair, we computed the Pearson correlation between their Δf/f traces after excluding frames in which both ROIs were silent. Pairs with no co-active frames were assigned a correlation of –1. Excluding jointly silenced frames made the correlation reflect covariation in transient amplitude rather than coincident activity or silence. Correlations were calculated from the full time series rather than spatial PF maps so that shared co-fluctuation at any moment could be detected in cells that remapped after the reward switch (fields move) or were active only during stationary periods (no spatial fields). The resulting correlation matrix was thresholded separately for each animal at 0.3-0.4 based on inspecting of the correlation histogram; fig. S5B to S5D). ROIs with at least one above-threshold partner entered the clustering process. The thresholded matrix was binarized (1, above threshold; 0 otherwise) and then converted to a distance matrix (distance = 1 − binary value), and hierarchically clustered using weighted linkage. Cluster number was selected by maximizing a goodness score defined, for each cluster, as mean within-cluster correlation minus mean between-cluster correlation. This score favored tightly correlated clusters that were well separated from other ROIs and penalized both overmerging and oversplitting. To reduce merging of distinct axons, we selected the largest number of clusters whose goodness remained within 97% of the maximum and visually confirmed each solution from ROI traces and footprints. Traces from ROIs within a cluster were normalized to their respective baselines and averaged to form one unit, which alone was used in subsequent analysis. ROIs with very small footprints (≤ 25 pixels) were excluded.

### Place cell classification

PCs were identified as described for somata (SI above the 95th percentile of a 200-shuffle distribution), with two axon-specific modifications. First, the reliability threshold was 0.3 rather than 0.2. Second, a coherence criterion excluded fragmented or very narrow fields. Coherence was calculated from the trial-averaged PFs (50 spatial bins by 1) as the Pearson correlation between activity in each bin and the mean activity in its neighboring bins (±4 bins). Only cells with coherence >0.9 were retained; this excluded ∼5-10% candidate PCs. Coherence was independent of PC’s reference frame, and relaxing this criterion did not alter the conclusions.

#### Population vector correlation analysis

To quantify how the PC population encodes location, we performed population vector (PV) analysis(*14*). At each of the 50 belt spatial bin positions, the PV was an N-cell vector whose elements were the trial-averaged Δf/f values of individual neurons in the corresponding 3.6-cm bin. Pearson correlations between PVs at all pairs of positions yielded the similarity matrix M. Each element M_i,j_ represents the correlation coefficient between PVs at locations i and j. M has dimensions of 50 by 50 (50 spatial bins per track), measuring the position representation similarity between conditions. PV similarity was quantified as the mean correlation along the diagonal of M between day 0 and day 1 PFs (Fig. 1J). Only cells with reliable PFs on both days were included; including all cells did not alter conclusions.

#### BTSP event identification

Behavioral timescale synaptic plasticity (BTSP) events were detected from the spatial Δf/f maps adapted from previously published methods(*1*, *5*, *15*). A candidate event was a contiguous set of spatial bins with significant calcium events (defined under *Pre-processing and signal processing*). Candidate bins separated by gaps of ≤2 silent bins were merged. For each event, peak amplitude and position (bin and lap) were measured. Pre- and post-event activity was evaluated within a ±12-bin (∼±45cm) window centered on the event peak. Putative BTSP events met four criteria. (1) Top-amplitude event: peak Δf/f was within the top 10% of that cell’s Ca²⁺ events and exceeded an absolute threshold of 2 Δf/f. In reward-switching sessions, the top 10% threshold was computed separately before and after the switch. (2) Stable location: the center of mass (COM) of the five-lap post-event mean field was within 12 bins (∼43 cm) of the induction-lap COM, excluding spurious large jumps. (3) Reliable new field: significant transients occurred within the ±12-bin window on ≥60% (≥3/5) of the five post-event laps, and the trial-averaged activity from the five post-event laps formed a valid place field as defined below. (4) Backward shift: the post-event field COM was behind the induction location (negative COM shift). Events during the final five laps (laps 96–100) were excluded because five post-event laps were unavailable. Including these events did not alter the conclusions. All putative BTSP events were detected, so a cell could have more than one BTSP events. All BTSP-related analyses except event count (Fig. 3A; 5K; fig. S8H) were restricted to each cell’s first BTSP event in each condition, because only the first induction event is preceded by a clean pre-event window in which residual activity from previous experience can be isolated. Activity before later events could reflect both residual activity from prior sessions and plasticity already induced in the current session.

Single-lap event fields were distinguished from trial-averaged place fields. On the single induction lap (no averaging), the event field comprised consecutive bins with Δf/f >35% of that lap’s peak, and the event field width was the number of bins in this interval. A place field (PF) was a single 50-bin Δf/f trace obtained by averaging Δf/f across a five-lap pre- or post-BTSP window or across an entire session. A trial-averaged PF was valid only if it met four conditions: (i) it contained consecutive bins with Δf/f values >35% of the field peak, which defined the local fields; (ii) field width was <90% of the track; (iii) peak in-field Δf/f exceeded 0.2; and (iv) mean in-field Δf/f exceeded twice the mean Δf/f outside the field defined in criterion i. BTSP criterion 3 applied this valid-field definition to the five-lap post-event mean. PF COM was computed only in bins within the field defined in criterion i:

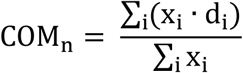

Where x_i_ is the Δf/f within spatial bin i, and d_i_ is the spatial position of that bin. The COM was assigned NaN if no valid field was present for the 5-lap mean PF. Induction velocity was defined as the mean running speed within the induction-lap event field.

#### Residual activity analysis

For each cell with a BTSP event, residual activity was defined as the trial-averaged Δf/f map (50 bins; mean Δf/f per bin) from the five laps immediately preceding that cell’s first BTSP event in each condition, or from all available preceding laps if fewer than five were available. A pre-event window was classified as containing active residual if ≥3 consecutive active bins across the five-lap window (5 laps by 50 spatial bins) at any location. Cells with no detectable calcium activity in this window were classified as having silent residual. Note that here ‘silent’ reports the absence of detectable somatic Ca2+ transients under our criteria, not the absence of any electrical activity (e.g., subthreshold membrane potential changes). Cells without a detected BTSP event lacked a first-BTSP lap, so residual activity was measured in the five laps before a reference lap matched to the typical timing of the first BTSP event. In training-stability mice, the reference lap was the median first-BTSP lap of cells with BTSP in each group (laps 15–19). In DREADD mice, DCZ and control cells were compared directly, so both groups shared one reference lap, the median first-BTSP lap pooled across groups (lap 17; laps 12–16). These cells were included only in analyses that explicitly quantified residual amplitude or stability in cells without detected BTSP and were excluded from analyses relating residual location to BTSP-event location or BTSP-event stability. This distinction is specified in each figure legend.

Residual stability was the Pearson correlation coefficient (c.c.) across 50 spatial bins between the residual PF and the indicated reference PF: day 0, day 1, or the induction-lap (BTSP-lap) PF. BTSP stability was calculated identically, with the induction-lap PF substituted for the residual PF. All correlations were computed between binned spatial Δf/f PFs (50 by 1 vectors), not binarized fields or raw Δf/f traces. As an amplitude- and shape-insensitive alternative, peak-position stability was calculated as 1 minus the circular distance between the two PF peak bins normalized by the track length which yielded the consistent conclusions.

Three additional measures described residual spatial structure relative to the subsequent BTSP event. Residual selectivity was defined as (mean in-field Δf/f − mean out-of-field Δf/f) / (mean in-field Δf/f + mean out-of-field Δf/f). In-field bins comprised ±12 bin (∼43cm) around either the residual peak (Fig. 3I) or the subsequent BTSP-event peak (Fig. 3L; fig. S6), as indicated. Values near 1 indicated spatially selective residual activity, whereas values near 0 indicated nonselective activity. BTSP-to-residual proximity was derived from the circular distance between the BTSP-event and residual peak bins and normalized such that 1 denoted the same bin and 0 denoted the maximal separation (half the track, ∼90 cm; Fig. 3I). To test whether BTSP shifted tuning toward the consolidated day-0 representation, ΔPF stability was defined as post-event PF stability minus pre-event PF stability. Each PF stability value was the Pearson c.c. between the corresponding five-lap mean PF and the day-0 PF template (50 by 1 spatial Δf/f vectors; Fig. 3J). Positive ΔPF-stability values indicated a shift toward the day-0 representation. The same definitions were used for the DREADD and saline cohorts (Fig. 5; fig. S13).

For population-average tuning curves aligned to plasticity events (Fig. 3F; fig. S7), each cell’s five-lap pre-event mean PF, five-lap post-event mean PF, and induction-lap activity were circularly shifted to center the BTSP-event peak at bin 25 and then averaged across cells. To summarize the timing and position of plasticity separately (fig. S6), BTSP events were binned by induction lap in 20-lap blocks and independently by spatial position. The same BTSP detection and alignment procedures were applied to DREADD and saline controls.

#### Training stability analysis

Training stability was analyzed in five mice imaged on three consecutive training days on belt A (days –2, –1, and 0; no reward switch) and on belt B on day 1. For each cell, training stability was the mean of the three pairwise PF Pearson correlations among three training days (day –2 versus –1, –1 versus 0, and –2 versus 0) with each correlation calculated across the 50 bins of the two days’ trial-averaged Δf/f maps. Cross-context generalization was the corresponding correlation between day 0 (belt A) and day 1 (belt B). Cells were included if they were PCs on day 0, regardless of their classification on earlier days, because silent or unreliable PF on an earlier day also reflected weak consolidation. Restricting the analysis to cells with reliable PFs on both days 0 and 1 did not alter the conclusions. Within each mouse, day-0 PCs were ranked by training stability, and highest and lowest 20% were designated the most- and least-stable groups. Values were averaged within mouse to yield one value per group before cross-mouse comparison. Percentile thresholds were therefore determined separately for each mouse.

#### Bayesian decoding

Bayesian decoding followed established methods(*16–19*) and used custom MATLAB scripts adapted for calcium imaging. The decoder assumed Poisson statistics, conditional independence among neurons, and a uniform spatial prior. The decoder estimated the animal’s position from the moment-to-moment population activity, given a template that described each cell’s activity as a function of position.

Signal preprocessing. Each cell’s Δf/f trace was smoothed with a six-frame moving average and converted to a decoding signal (“baye-Δf/f”) containing only the rising phase of significant transients. Rising phases were continuous periods of ≥7 frames (∼0.23 s) during which Δf/f exceeded the cell-specific signal threshold and had a positive first derivative. All other samples were set to zero. This procedure isolated active transients from baseline fluctuations and reduced the positional smear introduced by the slow decay of the calcium indicator. The baye-Δf/f traces were binned into 50-bin spatial maps and averaged across laps using the same procedure as for standard Δf/f PF maps, yielding a decoding template PF for each cell. Stacking these templates across cells produced a 50-spatial-bin by N-cell activity rate matrix, where N was the number of PCs.

Decoding procedure. Spatial position was decoded in consecutive non-overlapping windows of ∼150 ms (5 frames). The posterior probability over position p was:

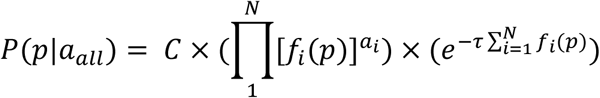

Here, *a_all_* is the activity vector across all cells, *C* is the normalization constant, *τ* is the temporal bin duration (0.15 s), *N* indexes cells, *f_i_*(*p*) is the spatial activity template for cell *i*, and *a_i_* is that cell’s activity in the current time bin. A small constant equal to 5% of the mean template activity was added to *f* so that silent template bins did not force the posterior to zero. The posterior was evaluated in log space for numerical stability. Decoded position was the spatial bin with the maximum posterior probability.

### Cross-day generalization analysis

To test how well the day 0 spatial code predicted position on day 1, we used day 0 PF templates to establish the decoder and applied it to decode position on day 1 using day 1 activity. Decoding error was the absolute distance (cm) between decoded and actual position in each 0.15s window, averaged within each lap. Windows with running speed < 5 cm/s were excluded. Analyses included all cells classified as PCs on *both* days 0 *and* 1. PC counts were comparable between conditions (same-versus cross-environment: 97±15 versus 82±17 PCs, p = 0.54, unpaired *t*-test). The same procedure was applied to DREADD and saline mice, using drug-free day-0 templates to decode day 1 position based on day 1 population activity.

#### Principal component analysis (PCA) and subspace alignment

To examine dimensionality and alignment of neural subspaces across days, we performed PCA on temporal activity patterns using baye-Δf/f data. Neural activity X was organized as time-by-cells matrices (rows: time bin *T*, *τ* = 0.15s; columns: neurons *N*), excluding stationary periods (speed < 0.05 m/s).

#### Data standardization and PCA

For each recording day, neural activity was z-scored independently per day and per cell across time, yielding X*^z^*. PCA was applied to X*^z^* (*T* by *N*) to identify principal components ordered by descending variance explained. Each principal component (*N* × 1 vectors) described a direction in neural space; specific combinations of principal components defined subspaces. PCA yielded: (1) loadings *coeff* ( *N* × *components*), where each column is an orthonormal vector (PC-1,2,3,…), and (2) *scores* (*T* × *components*) where each column contains the projection of the X*^z^* onto corresponding components.

### Variance explained by cross-day subspaces

To quantify how much day 1 neural variance was explained by the day 0 neural subspace, both days’ data were z-scored (yielding 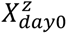 and 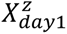), and PCA was applied separately to yield *coeff_day0_* and *coeff_day1_*.

We quantified explained variance in two ways:

(1) Individual component variance: For each of the target day’s (day 0 or day 1) top *k* principal components (typically *k* = 10), we projected 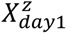 onto that single component *coeff_day_*__*tarttet*_ [:, *i*] yielding:

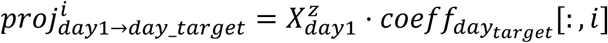

We then calculated the variance of the resulting 1D projection as:

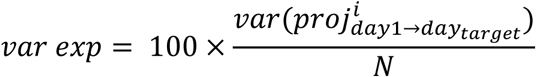

Where 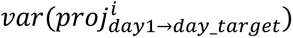 is the variance of the projection onto the *i*-th components of *day*_*target*, and *N* is the total variance of z-scored 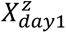 Here N denotes the total variance of the z-scored day-1 matrix (the sum of all cell variances).

(2) Cumulative variance by top *k* components: We projected 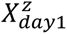 onto top 2 components from *day*_*target* simultaneously (e.g., Fig. 2B) yielding 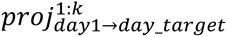 and calculated the variance captured by this 2D projection:

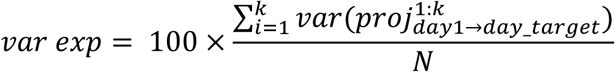

Controls included: (1) variance explained by day 1’s own top *k* components, and (2) shuffles randomly selecting *k* columns from *coeff_day_*_0_ (*k* = 2, 1000 iterations).

Identification of critical dimensions. We focused on the top two principal components because: (1) the top three components of activity from familiar environment explained a comparable amount of variance in the novel environment as same-environment controls (p>0.5; fig. S2A) and exceeded chance levels (p<0.05; fig S2A). (2) decoding error exhibited characteristic U-shaped profiles (fig. S2B) decreasing with 2-3 components then increasing with additional dimensions.

#### Neural trajectory visualization

Visualizations used spatially binned maps; quantitative analyses used time-binned activity. Three conditions: (1) Spatial maps from day 0 and day 1 were preprocessed before obtaining subspaces using PCA, and projecting z-scored activity onto corresponding subspaces. Trajectories formed rings with coherent color when coded by reward distance. (2) Same as (1), but using concatenated data per group, with neurons split into stable and remapped cells before PCA. (3) Similar to (2), but using concatenated data from all animals with reward switching on belt A (day 1), and B (day 2) where we tracked the same neurons across both days (5 mice). PCA was performed once on concatenated (A-pre, A-post, B-pre, B-post, 50 laps each) spatial maps. Activity was projected onto top components to visualize task-relevant (expanded rings) and context-relevant (belt A/B separation) subspaces. All visualizations used 10-lap mean data before PCA.

#### Subspace alignment analysis

Z-scored day 1 activity 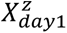 was divided into 20 sections (5 laps each): 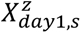 (*s* is the section number). PCA obtained section-specific *coeff_day_*_1,*s*_ (cells by components). We quantified subspace alignment adapted from(*20–22*), using:

(1) Subspace alignment index A:

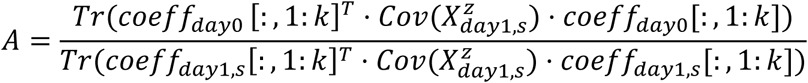

to measure the amount of section *s* variance shared by day 0 and day 1 top *k* subspaces, where *coeff_day_*_0_[:, 1: *k*] and *coeff_day_*_1,*s*_[:, 1: *k*] defined the top k principal components from day 0 (entire session) and day 1 (per section), respectively. *Cov* 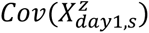 is the covariance of the matrix 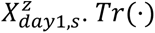. *Tr*(⋅) is the matrix trace. The numerator measures day 1 total variance captured by the top *k* principal components from day 0. The denominator normalizes the alignment index by maximum variance capturable by any *k*-dimensional subspace. Thus, *A* ∈ [0 1]. We used *k* = 2 as described above (fig. S2A; S2B). We compared three analyses: First, for overall population dynamics (Fig. 2C), we calculated subspace alignment index per section for all PCs with reliable PFs on both days. Second, to compare stable versus remapped cells, we separated stable and remapped PCs, performed PCA, calculated alignment index per section per cell type, then averaged across sections per mouse (One animal per group excluded for <20 cells for each cell type). Third, to assess BTSP’s role in subspace alignment by shaping synaptic weights, we ranked PCs that had reliable PFs on both days 0 and 1 by the lap of their first day 1 BTSP event (cells without BTSP ranked last) and, within each mouse, assigned the earlier half to the “with early BTSP” group and the later half to the “without early BTSP” group so that both groups had the same number of cells, calculated alignment index per section for each cell type, then averaged across sections per mouse (One animal per group excluded for <20 cells for each cell type). To allow better comparison across populations of different sizes, each alignment index was z-scored relative to a section-specific distribution from 1000 random two-dimensional subspaces drawn from the day-0 components, computed separately for each subpopulation. A z-score of 0 indicated chance-level alignment, and positive values indicated above-chance alignment. The shuffle-normalized score, rather than the raw ratio, was used for all between-group comparisons. The same procedure was applied to DREADD and saline groups. (2) Principal angles: From *coeff_day_*_1, *s*_[:, 1: 2] and *coeff_day_*_0_[:, 1: 2] we calculated principal angles *θ*_1_ using ‘subspacea’(*23*) (https://www.mathworks.com/matlabcentral/fileexchange/55-subspacea-angles-between-subspaces) in MATLAB which computes all principal angles between subspaces through singular value decomposition. The function identifies *θ*_1_ as the smallest angle between any pairs of orthonormal vectors, one from each subspace, while *θ*_2_ is the angle between a second pair of unit vectors that are orthogonal to the first pair within their respective subspaces. We took the mean of *θ*_1_ and *θ*_2_ to represent subspace alignment angle. Using *θ*_1_ alone did not alter conclusions.

#### Subspace-based decoding

Subspace decoding adapted the Bayesian decoder with Gaussian probability function. Day 1 z-scored data 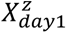 was projected onto day 0’s top *k* PCs (varying from 1 to total cell count, denoted ‘X’ as animals differ in recorded cell numbers), yielding *propp_day_*_1→*day*0_ (*T_day_*_1_ × *k*). Spatial templates *μ_c_*(*pos*) (50 spatial bins × *k*) were constructed by averaging day 0’s projected activities *propp_day_*_0→*day*0_within each spatial bin. Standard deviations *σ_c_*(*pos*) were calculated as 0.1 times each component’s *SD* across day 0. For each time bin *t*, the likelihood of position pos given the *k*-dimensional projected activity *propp_day_*_1→*day*0_[*t*, :]:

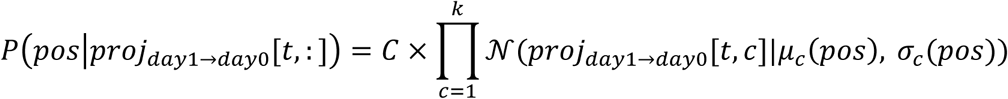

Where *P*(*pos*|*propp_day_*_1→*day*0_[*t*, :]) is the probability of position *pos* given the projected neural activity *propp_day_*_1→*day*0_[*t*, :]. *C* is the normalization constant. N is the Gaussian probability density function; *propp_day_*_1→*day*0_[*t*, :] is the scalar projected activity on the *c* -th principal component at time *t*. Decoded position was the position with maximum likelihood. Decoding error was absolute difference between decoded and actual positions, averaged across time bins. Subspace-based decoding used all PCs that had reliable PFs on both days.

#### Co-activity

For PCs with reliable PFs on both day 0 (belt A) and day 1 (belt B), pairwise co-activity was the total time for which both neurons were active divided by the total time either neuron was active. Cell pairs were pooled across animals, and day-1 co-activity was regressed against day 0 co-activity with a single linear model. Day 1 co-activity was computed over the first 20 laps, representing early exposure to the novel belt B, and day 0 co-activity was computed over the final 20 laps (laps 81-100) on the well-trained familiar belt.

### Characterization of the day-0 task subspace

To characterize the day-0 task subspace (fig. S3), we identified cells with the strongest contributions to the leading principal components and examined their encoded properties. For each mouse, the day-0 PCA loading matrix (coeff, defined above) was used to assign each cell a loading weight equal to its mean absolute loading on PC-1 and PC-2, (|PC-1| + |PC-2|)/2. Loading weights were calculated exclusively from day-0 data. Cells were ranked within each mouse and assigned to strong- and weak-loading groups using the upper and lower thirds of that mouse’s loading-weight distribution. Group assignment was therefore animal-specific, and the pooled 33rd- and 67th-percentile lines in fig. S3A are shown only for visualization. Included cells were reliable PCs on both days 0 and day 1 (n = 493 cells from 6 mice, 163 cells per third).

Strong- and weak-loading cells were compared across four single-cell properties computed from the 50-bin maps. Field reliability (the lap-1 reliability defined under “Place-cell identification”), PF onset lap, and single-cell SI were compared between groups with two-sample Kolmogorov-Smirnov tests on pooled cell distributions (fig. S3B to S3D). Spatial tuning was displayed as peak-normalized Δf/f maps, sorted by day-0 peak position and shown for both days 0 and 1 in each group (fig. S3E). Cross-day tuning stability was the Pearson correlation across 50 spatial bins between each cell’s day-0 and day-1 trial-averaged PFs and was compared between groups with a two-sample Kolmogorov-Smirnov test (fig. S3F).

### Validation of spatial metrics against running-speed-dependent dwell-time bias

Because dwell time differed among spatial bins as a function of running speed, standard binned maps, which averaged Δf/f across all imaging frames in each bin, could have speed-dependent bias. We therefore recalculated spatial maps using only frames that contained a significant calcium transient (the transient-based pipeline) and compared them with maps generated from the standard Δf/f pipeline (fig. S8).

We first tested whether differences between pipelines depended on running speed. For each cell, the standard-minus-transient map difference was computed for every bin and lap. We focused on only bins that contained significant transients (fig. S8B; example running-speed map in fig. S8A) because bins without significant transients are not meaningfully affected by dwell-time-dependent averaging and therefore do not contribute to this potential bias. Samples were pooled across six mice, and Spearman correlation was calculated between the map difference and running speed in the corresponding bin and lap (fig. S8C). The corresponding per-bin difference in day-0-versus-day-1 PV correlation was calculated and averaged across mice (fig. S8D) and displayed with the matched group-mean running-speed map (fig. S8E).

We then tested whether the main day 1 results were reproduced by the transient-based pipeline. Lap-wise PF correlation with the day 0 template, averaged across stable PCs (fig. S8F), cross-day PV similarity across reliable PCs (fig. S8G), and mean BTSP event count per PC (fig. S8H) were compared between pipelines with paired t-tests across mice. At the event level, both pipelines were used to calculate peak-Δf/f traces aligned to the first BTSP-induction lap (fig. S8I), distributions of the distance between residual-activity and day-0 PF peaks (fig. S8J, two-sample Kolmogorov-Smirnov test), and the relationship between residual stability and BTSP-event stability (fig. S8K, Pearson correlation). Both pipelines supported the same conclusions, indicating that the spatial and plasticity results were not explained by running-speed-dependent dwell-time bias.

#### Computational model of CA1 generalization

We developed a network-level model of two-compartment CA1 pyramidal neurons that integrates the latest experimental findings and theoretical insights. This model includes realistic presynaptic inputs from CA3 and the entorhinal cortex (EC), as well as feedforward and feedback inhibition, effectively capturing the complexity of the hippocampal circuit. We have implemented BTSP at CA3–CA1 synapses along with a biophysically informed mechanism for plateau potential generation. Additionally, we have incorporated a cascade-based model of synaptic consolidation to study long-term stabilization processes. Moreover, we introduced phenomenological rules to systematically vary presynaptic input patterns from entorhinal cortex layer III (EC3) and CA3 pathways across different contexts. The model simulates animals running on a track over multiple days, during which synaptic weights evolve through plasticity and consolidation.

#### Presynaptic inputs and CA1 activity

Each CA3 pyramidal neuron was modeled as a place cell with a Gaussian spatial tuning curve. The firing rate of CA3 neuron *pp* at position *x* (normalized between 0 and 1) is

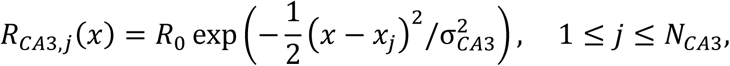

where *R*_0_ = 40*Hz* is the maximal firing rate, *x_jj_* is the preferred spatial location of neuron j and σ*_CA_*_3_ = 0.0833 controls the width of the place field. In contrast, each EC3 neuron is modeled as a two-state Markov process following Grienberger and Magee(*5*), with neuron-specific preferred locations shaping the CA1 representation.

The peri-somatic input *I^s^*(*t*) to CA1 neuron *i* was calculated as a weighted sum of CA3 place-cell activitities,

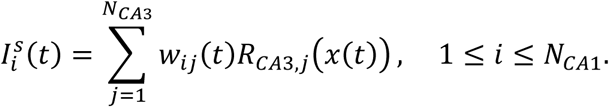

where *w_ijj_*(*t*) denotes the synaptic weight from CA3 neuron *pp* to CA1 neuron *i* and *x*(*t*) is the animal’s spatial position. The firing rate of CA1 neuron *i* was described by a rectified linear gain function of somatic inputs:

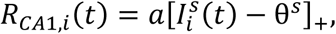

where [⋅]+= max(⋅,0). We set gain *a* = 2 and threshold θ*^s^* = 8 to match empirical firing rates.

#### Plateau potential probability

Plateau potentials were modeled as stochastic, all-or-none events driven by both somatic and distal dendritic inputs. The probability that CA1 neuron *i* generates a plateau potential at time *t* depends on its somatic input 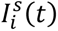, distal dendrite input 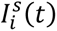 and the feedback inhibition *I_CA_*_1,*in*ℎ_(*t*):

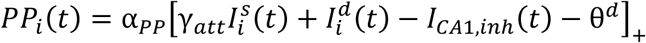

where α*_PP_* = 0.25*s*^−1^modulates the overall plateau probability, γ*_att_* = 0.096 denotes the attenuation of peri-somatic input to the distal dendrite and θ*^d^* = 0.5 is the dendritic threshold. The plateau potential event for each neuron is independently drawn as a non-homogeneous Poisson process based on the probability *PP_i_*(*t*).

The total distal dendritic input 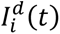 was modeled as the net input of the weighted sum of EC3 activity minus feedforward inhibition:

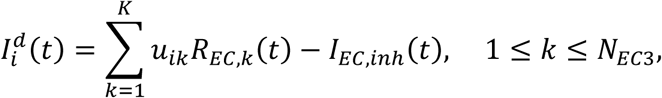

where *u_ik_* is the synaptic strength from EC3 neuron *k* to CA1 neuron *i*. Each CA1 neuron received inputs from a randomly selected subset comprising 5% of the total presynaptic EC3 neurons, with synaptic strengths randomly drawn from the uniform distribution *u_ik_* ∼ *U*(0,12.5) and remain fixed during the simulation. Feedforward inhibition of EC3 input is modeled as

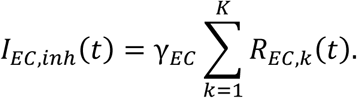

with γ*_EC_* = 0.25 controlling the strength of inhibitory interneurons in the EC3 pathway. The feedback inhibition *I_CA_*_1,*in*ℎ_(*t*) represents the contribution of CA1 interneurons on plateau initiation and was modeled as

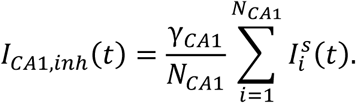

with γ*_CA_*_1_ = 0.286 controlling the strength of inhibition.

#### Synaptic plasticity and consolidation

To capture rapid experience-dependent synaptic changes, we implemented BTSP following Milstein et al. (2021)(*24*). Each CA3–CA1 synapse maintained an eligibility trace *ET_jj_*(*t*) that accumulates recent presynaptic activity and a plateau trace *PT_i_*(*t*) that reports recent plateau events:

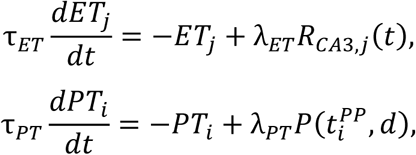

where *t_PPi_* denotes the timing of plateau potential for CA1 neuron *i* and *d* is the duration of plateau potential (fixed to be 300 ms). Synaptic weights evolved according to:

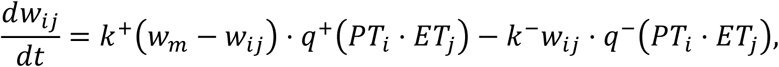

where *w_m_* is the maximal synaptic strength, *k*_+_ and *k*_−_ are potentiation and depression rates, and the functions *q*^±^ are sigmoidal functions of the product *PT_i_* ⋅ *ET_jj_*. These sigmoidal functions are defined as:

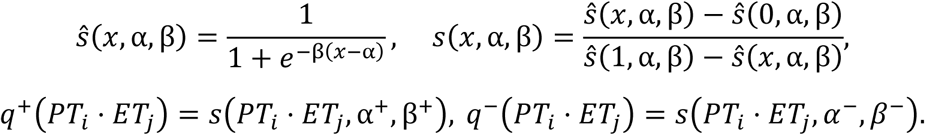

with parameters α^±^ and β^±^controlling the thresholds and slopes of potentiation and depression. Parameter values were taken from the original model in Milstein et al.(*24*).

To model long-term consolidation across days, we used the cascade model of synapse proposed by ref.(*25*). For each CA3-CA1 synapse *w_ijj_*, we introduce a chain of hidden states 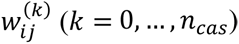 (*k* = 0, …, *n _cas_*). The synaptic weight is related to the first hidden state via *w_ijj_* = 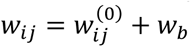 + *w_b_*, where *w_b_* = 0.75 is the baseline synaptic weight. The dynamics of the cascades are governed by

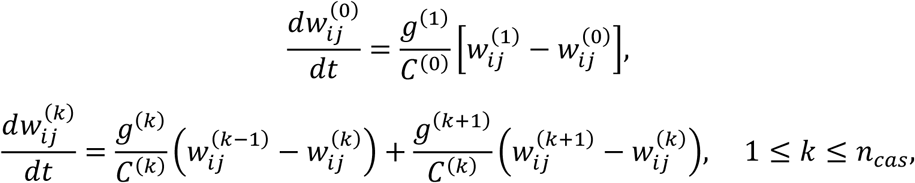

with boundary conditions 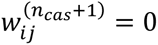 = 0 and parameters *C*^(*k*)^ = 2*^k^T*, *g*^(*k*)^ = 2^−*k*^, where *T* =

24 hours and *n_cas_* = 10 set the characteristic consolidation timescale. For CA3-CA1 synaptic stability impaired case (Fig. 4F; fig. S9J and S9K), we set *T* = 6 hours and *n_cas_* = 0. We also performed the manipulation of removing cascade mechanism alone (*n_cas_* = 0 but retaining *T* = 24 hours). This manipulation partially reduced synapse stability and residual activity as strong synaptic weights persisted through exponential decay given the long time constant. However, qualitative conclusions (fig. S9J and S9K) remained consistent.

#### Input patterns and simulation protocol

Across distinct contexts, we systematically varied the overlap of presynaptic representations. In two contexts, CA3 neurons were divided into four distinct groups: (1) neurons with the same preferred spatial locations in both environments; (2) neurons active in both environments but with uncorrelated preferred locations; (3) neurons active only in environment A; and (4) neurons active only in environment B. These groups were assigned in proportions 6:2:1:1. Similarly, EC3 neurons were split into two groups: neurons with the same preferred location across contexts and neurons active in both environments but with uncorrelated preferred locations, in proportions 6:4. We set *N_CA_*_1_ = 2500, *N_CA_*_3_ = 200, *N_EC_*_3_ = 2000, respectively.

Within individual sessions, we further incorporated the dynamic reconstitution of CA3 place fields by progressively refining preferred positions across laps, consistent with experimental observations. Each CA3 neuron’s preferred location at lap *k* was determined by allowing its original preferred location (*x*_0_) to drift according to the following equation:

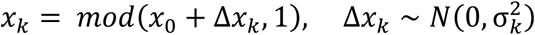

where σ*_k_* = σ_0_ exp(−*k*/τ*_cst_*) describes the diminishing spatial variability across laps. The parameters controlling initial variability (σ_0_) and refinement rate (τ*_cst_*) were set as σ_0_ = 0.25, τ*_cst_* = 5 laps for familiar environments, and τ*_cst_* = 15 laps for novel environments.

To model learning across days, we simulated animals running 50 laps per day, with each lap lasting 10 seconds, over seven consecutive days in a familiar environment (context A). On day 8, the animals remained in familiar environment A or switched to a novel environment (context B) for another 50 laps. Throughout the simulated period, synaptic weights evolved under the BTSP rule within sessions and underwent cascade consolidation across days.

#### Quantification and Statistics

In our simulations, we defined several metrics to quantify place cell stability and plasticity across different timescales. A place cell was considered “stable” from day 0 to day 1 if its place field on day 1 remained within 15 cm of its original location on day 0, whereas cells that did not maintain their place field on day 1 were classified as “remapped.” As a continuous measure of the stability of the place field in the day-to-day, we calculate the stability of the place field as one minus the normalized distance between the place field of a cell on day 0 and day 1. Similarly, the stability of the BTSP plateau potential location was quantified as one minus the normalized distance between the plateau location on day 1 and place field on day 0.

In addition, we also quantified the stability of neural activity patterns beyond the main place field: the stability of the residual activity of each cell was measured as the correlation between its day 1 residual activity pattern (5-lap average, before BTSP induction) and its firing activity on day 0, and the stability of distal dendritic activity was defined likewise as the correlation between the distal dendritic activity on day 1 (5-lap average, before BTSP induction) and its place cell firing pattern on day 0.

Over the full 7-day pre-training period in environment A, a cell was defined as “high training stability” if it maintained a place field at the same location on at least 5 of the 7 days (including day 0, the final training day), whereas cells defined as “low training stability” never had a field at the same location as on day 0 (Fig. 4F; fig. S9I), giving out a proportion of about 15% and 26% of all active cells, respectively. In CA3-CA1 synaptic stability impaired case where overall field stability was low, this multi-day criterion of “high training stability” was relaxed to 4 out of 7 days (Fig. 4F), resulting in a proportion of about 9% of all cells.

### Statistical Methods

The exact sample size (n) for each experimental group is reported in the figure legends or main text. While no statistical tests were used to predetermine sample sizes, our sample sizes align with previous publications(somata(*15*, *26–28*) and axon(*13*, *27*, *29*)) that used similar behavior tasks and were guided by the number of neurons that could be imaged or patched in awake, behaving mice. In some cases, where data distribution was assumed but not formally tested, parametric *t*-tests were applied to analyze the data. Experiments were randomized by randomly assigning littermate mice to the experimental groups. Data analyses were performed automatically without consideration of trial types or experimental groups, but they were not blinded to the experimenter. Unless otherwise indicated in the figures, data were presented as mean±SEM.

---

**Fig. S1.**
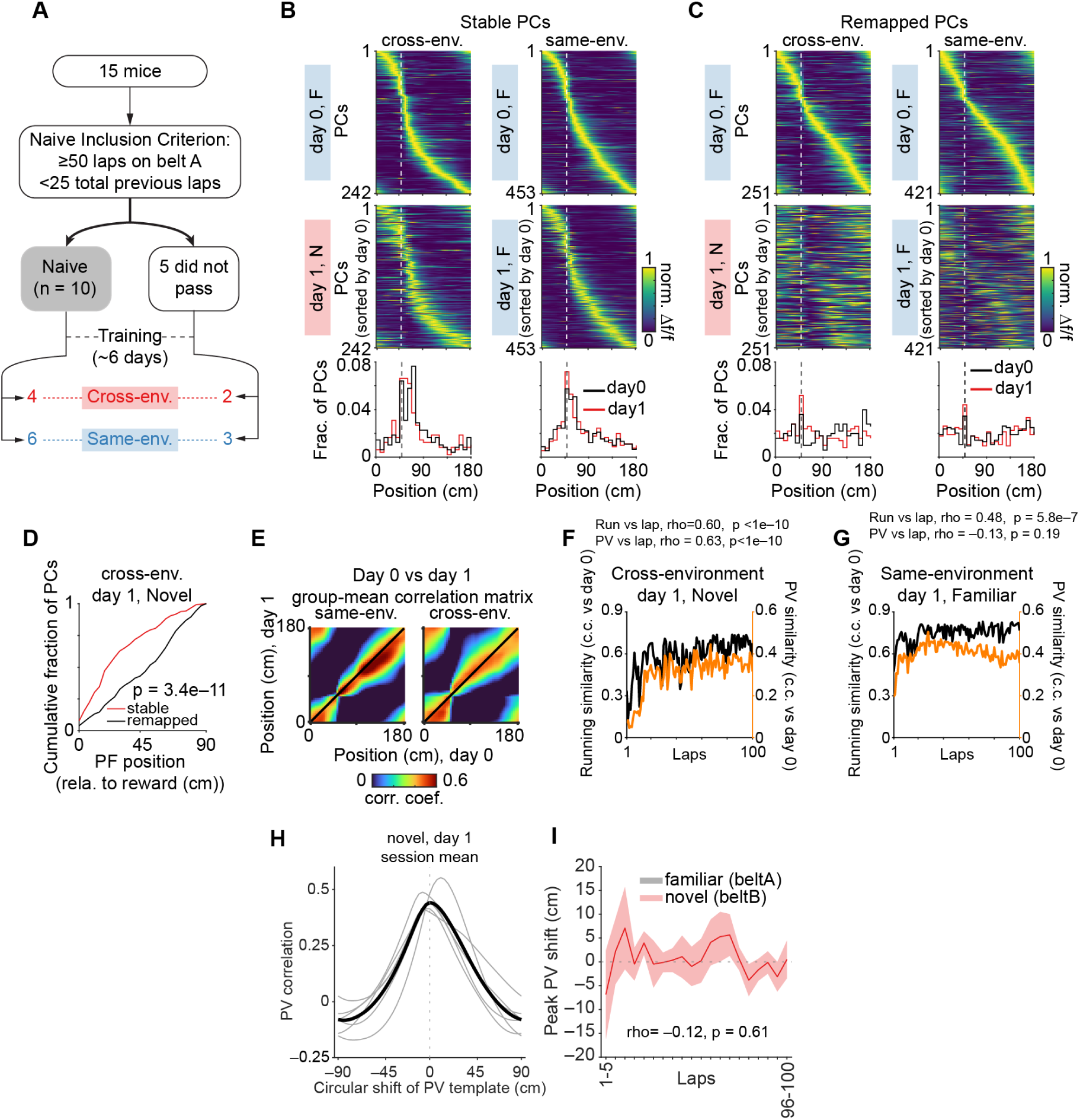
Behavioral and CA1 population dynamics across environments. **(A)** Schematic showing animal selection and grouping. Of 15 mice trained on belt A with a fixed reward, 10 met the naïve criterion (first day with ≥50 laps per session and <25 prior training laps) while five did not; All naïve mice had no prior experience with fixed reward training (0±0 day). After ∼6 days of training, naïve and non-naïve mice were assigned to cross-environment (familiar to novel environments but same reward rule; 4 + 2) or same-environment (familiar to familiar; 6 + 3) groups. **(B)** Left: peak-normalized mean spatial Δf/f for all cross-environment stable cells, day 0 (familiar env. A, top) and 1 (novel env. B, middle), sorted by day 0 PF peak (n = 242 cells, 6 mice). Bottom: PC fraction versus peak position for both days. Dashed line: reward location. Right: as left but for same-environment conditions (n = 453 cells, 9 mice). F/N: familiar/novel environments. **(C)** As in (B), but for remapped cells. Cross: 251 cells, 6 mice; Same: n = 421 cells, 9 mice. F/N: familiar/novel environments. **(D)** Cumulative PC fraction versus PF peak position relative to reward for stable and remapped cells in novel env. B (day 1). p = 3.4e–11, two-sample Kolmogorov-Smirnov test. Rewarded zone: ±30 cm around reward; non-rewarded zone: ±30 cm around 90cm from reward. Stable: rewarded versus non-rewarded, 26.8 versus 8.1 cells per 10.8 cm; remapped: rewarded versus non-rewarded, 15.4 versus 15.1 cells per 10.8 cm. **(E)** Group-mean PV correlation matrix comparing day 0 and 1 activity for same- (left, n = 9 mice) and cross-environment (right, n = 6 mice) conditions. Matrix shows c.c. between PVs at all locations. Black line: diagonal. **(F)** Trial-by-trial running profile similarity (black) and PV(orange) similarity for cross-environment mice. Similarity: day 1 novel env. B profile per lap versus day 0 familiar template (mean over laps 81-100) from env. A (mean over 6 mice). Each group-mean trace was correlated with lap number (Spearman, 100 laps): running similarity, rho = 0.60; PV similarity, rho = 0.63; both p < 1e–10. **(G)** As in (F), but for same-environment mice (mean over 9 mice). Running similarity, rho = 0.48, p = 5.8e–7; PV similarity, rho = –0.13, p = 0.19. **(H)** PV correlation as a function of circular shift of the day-1 novel-environment (env. B) population vector map relative to the day-0 familiar-environment (env. A) map, for cross-environment mice. Grey: individual mice; black: group mean. Peak near shift = 0 indicates no systematic spatial offset between environments (mean peak shift = −1.2 cm, p = 0.88, Wilcoxon signed-rank test vs. 0; n = 6 mice). **(I)** Signed peak PV shift across laps. The circular shift maximizing PV correlation between each single-lap novel-environment map and the day-0 familiar template was computed lap by lap per mouse, then averaged within non-overlapping 5-lap sections. shading: s.e.m. Spearman rho = −0.12, p = 0.61. n= 6 mice.

**Fig. S2.**
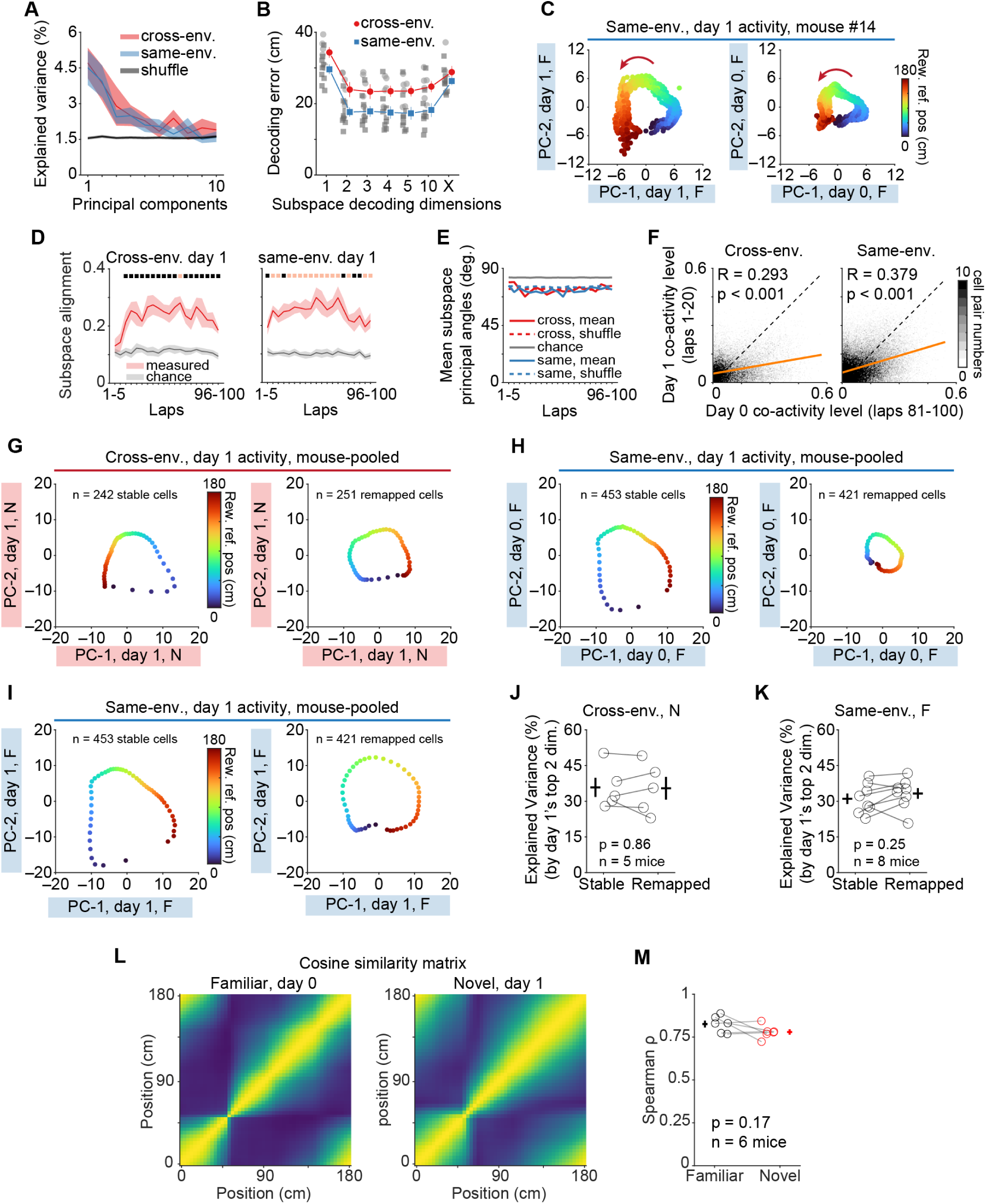
Additional low-dimensional subspace analysis. (A-B). Dimensionality of CA1 data in a linear spatial learning task. **(A)** Explained variance (%) in day 1 activity by top 10 principal components from day 0 for cross- (red, n = 6 mice), same-environment (blue, n = 9), and shuffled controls (black, 1000 shuffles of cross). Cross versus Same: p>0.05 for all components, unpaired *t*-test. Cross versus shuffle, p<0.05 for the top 3 components, paired *t*-test. Shading: S.E.M. **(B)** Mean decoding error versus day 0 subspace dimensions used for projecting day 1 activity for Cross- (red) and same-environment (blue) conditions. Grey dots: individual animals; blue/red dots: mean±sem. ‘X’: all available dimensions per animal. (C) Example mouse: 10-lap mean day 1 (familiar env. A) activity projected onto day 1 (left) and day 0 (familiar, right) principal components (PC-1, PC-2), colored by reward distance (cm). Red arrow: running direction. F: familiar environment. **(D)** Mean subspace alignment using ratio of variance explained by day 0’s top 2 principal components to day 1’s own top 2 principal components per section (5 laps/section) for cross- (left, red; n = 6 mice) and same-environment (right, blue; n = 9 mice) sessions. Grey: chance level using mean of shuffle control (subspace alignment using randomly selected 2 day0 principal components). Shading: sem. Black squares: p<0.05; red squares: p<0.001, paired t-test of measured versus shuffle mean across mice, per section. **(E)** Mean subspace principal angles between day 0 and day 1’s top 2 principal components per section (5 laps/section) for cross- (red; solid: mean, n = 6; dashed: shuffle, 95%) and same-environment (blue; solid: mean, n = 9; dashed: shuffle, 95%) animals. Grey line: chance. **(F)** Two-dimensional histogram of pairwise co-activity from day 1 (laps 1-20) versus day 0 (laps 81-100) for cross- (left, n = 6) and same-environment (right, n = 9) mice. Each bin counts cell pairs with corresponding co-activity values pooled across mice. Co-activity = (time both active) / (time either active). Orange line: linear fit; black dashed: unity. Cross-env.: R = 0.29; Same-env.: R = 0.38; both p<0.001, Pearson correlation. **(G)** Cross-lap mean within-day projection of day 1 (novel, “N”) activity onto day 1 principal components (PC-1, PC-2). Stable: left, n = 242 cells; remapped: right, n = 251 cells. Pooled data from 6 cross-env. mice. **(H)** Cross-lap mean cross-day projection of day 1 (familiar, “F”) activity onto day 0 principal components (PC-1, PC-2). Stable: left, n = 453 cells; remapped: right, n = 421 cells. Pooled data from 9 same-env. mice. **(I)** Cross-lap mean within-day projection of day 1 (familiar, “F”) activity onto day 1 principal components (PC-1, PC-2). Stable: left, n = 453 cells; remapped: right, n = 421 cells. Pooled data from 9 same-env. mice. **(J)** Explained variance (%) of day 1 (novel, “N”) activity by day 1’s top 2 principal components (PC-1, PC-2) for stable and remapped cells in cross-environment mice. Stable: 35.8±4; remapped: 35.4±4.8; p = 0.86, paired *t*-test, n = 5 mice. Circles: individual mice. Crosses: mean±s.e.m. One mouse excluded (<20 cells per group). **(K)** As in (J), but for same-environment. Stable: 31±2.2; remapped: 33.2±2.2; p = 0.25, paired *t*-test, n = 8 mice (one excluded). F: familiar environment. **(L)** Representational similarity matrices (RSMs) of bin-by-bin population vector (PV) cosine similarity, computed within each session using reliable PCs identified on both days (cross-environment mice). Left: day 0 (familiar env. A); right: day 1 (novel env. B); group-mean across n = 6 mice. Color: cosine similarity. Per mouse, the Spearman correlation between the upper triangles of the day-0 and day-1 RSMs was ρ = 0.88±0.021 (n = 6 mice), indicating that pairwise representational geometry is largely preserved across environments. F/N: familiar/novel environments. **(M)** Per-mouse Spearman’s correlation (ρ) between pairwise neural distance (1 – cosine similarity) and pairwise circular spatial distance across all bin-PV pairs, for cross-environment mice. Familiar (day 0, env. A): 0.83±0.02; novel (day 1, env. B): 0.78±0.016; p = 0.17, paired t-test, n = 6 mice. Circles: individual mice; grey lines connect the same mouse across environments; crosses: mean±s.e.m.

**Fig. S3.**
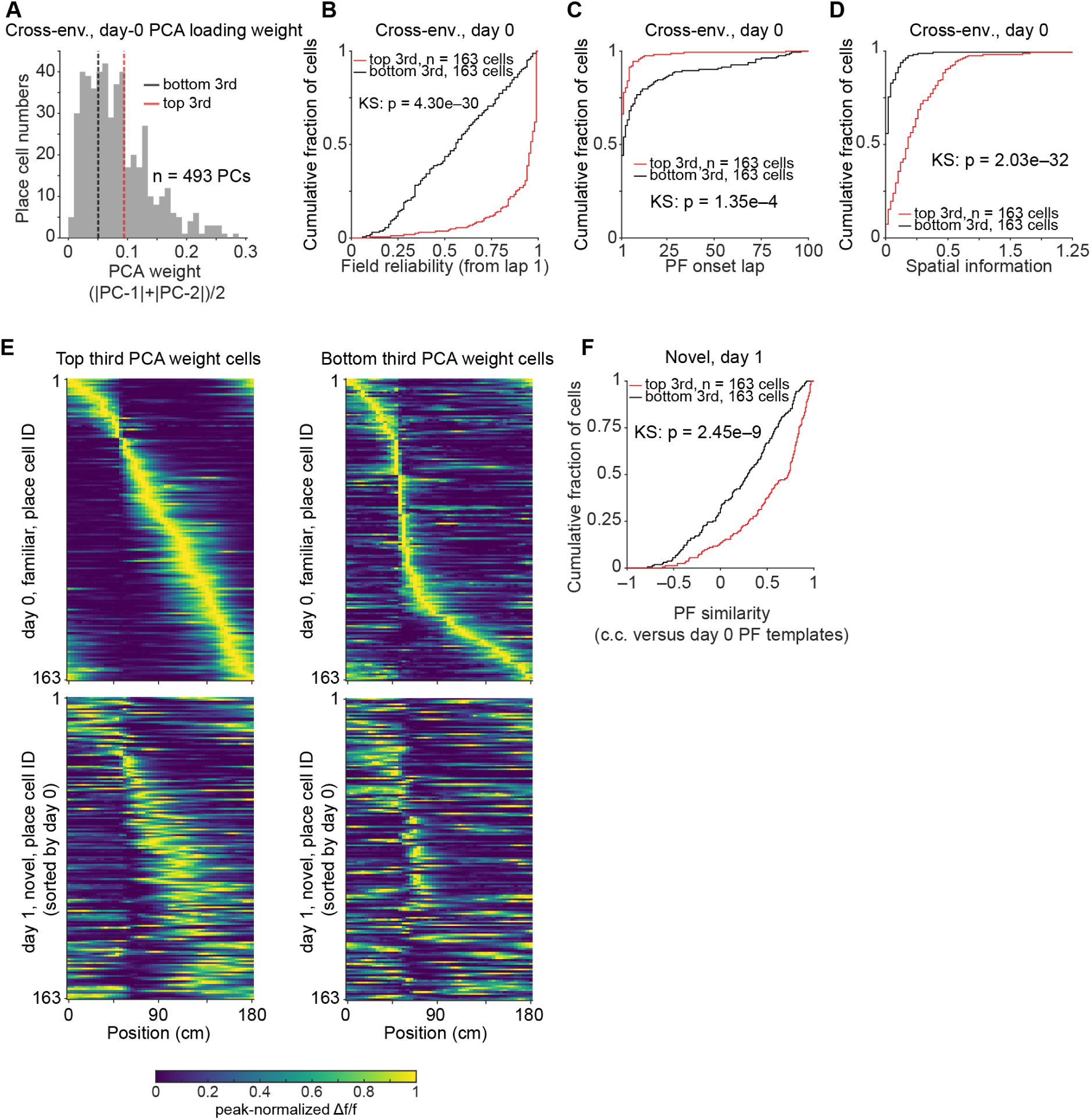
Characterization of task subspace PCA loadings. **(A)** Distribution of day0 PCA loading weights (mean of absolute value of principal component 1 and 2’s loadings) across all place cells in cross-environment mice (n = 493 cells, 6 mice). Dashed lines: pooled 33^rd^ (black, bottom-third threshold) and 67th (red, top-third threshold) percentiles shown for visualization. Top-third/bottom-third classification was performed independently within each mouse. PCA loading weight was computed exclusively from day 0 data. **(B)** Cumulative distribution of place field reliability for top-third (red, n = 163 cells) and bottom-third (black, n = 163 cells) loading weight cells on day 0 (familiar environment). Two-sample Kolmogorov-Smirnov test, p = 4.3e–30. **(C)** Cumulative distribution of PF onset lap on day 0. Two-sample Kolmogorov-Smirnov test, p = 1.35e–4. **(D)** Cumulative distribution of spatial information on day 0. Two-sample Kolmogorov-Smirnov test, p = 2.03e–32. **(E)** Peak-normalized spatial maps for top-third (left) and bottom-third (right) loading weight cells, sorted by day 0 peak position. Top row: day 0, familiar environment; bottom row: day 1, novel environment. Color bar: peak-normalized Δf/f. **(F)** Cumulative distribution of cross-day place field similarity (Pearson correlation of day 1 place fields versus day 0 place field template) for top-third (red) and bottom-third (black) loading weight cells. Two-sample Kolmogorov-Smirnov test, p = 2.5e–9.

**Fig. S4.**
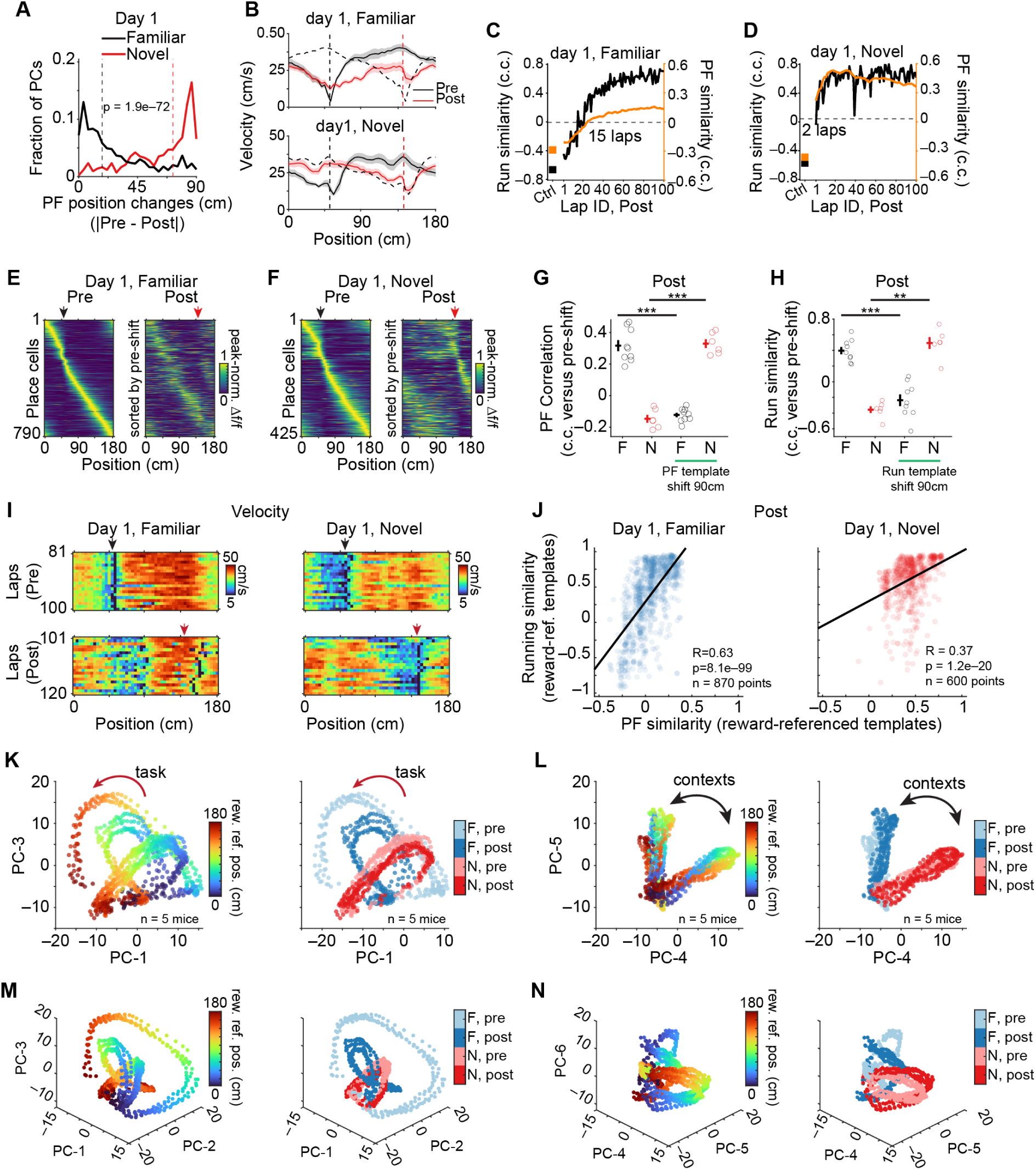
Additional analyses related to reference frame switching and subspace coding across environments. **(A)** Cell fraction versus PF position changes 20 laps post-reward switch on day 1. Familiar (env. A, n = 790 cells, 9 mice): median 18 cm (black); novel (env. B, n = 425 cells, 6 mice): median 72 cm (red). p = 1.9e–72, two-sample Kolmogorov-Smirnov test. Vertical dashed lines: median shifts for familiar (black) and novel (red) conditions. **(B)** Velocity pre- (black) and post- (red) reward switch in familiar (env. A, top) and novel (env. B, bottom) environments. Black dashed: Pre shifted 90 cm. Vertical dashed lines: reward locations for pre- (black) and post-switch (red) conditions. **(C)** Running (black) and PF (orange) similarity across 100 laps post-switch in familiar env. A (day 1). Similarity: *c*. *c*. versus reward-aligned template (mean, last 20 Pre laps). ‘Ctrl’: baseline (laps 81-100 (reward-aligned) versus laps 71-80 (original); both in Pre). Dashed: half-maximum running similarity (15 laps). **(D)** As in (C), for novel env. B. Half-maximum: 2 laps. **(E)** Peak-normalized mean spatial Δf/f for all PCs pre- (left) and post-reward switch (right) in familiar env. A (day 1). Arrows: reward at 50 (black) and 140 cm (red). n = 790 PCs, 9 same-environment mice. **(F)** Same as (E), but for novel environment B (day 1). n = 425 PCs, 6 cross-environment mice. **(G)** Mean PF correlation calculated relative to pre-switch original templates (left, F & N) versus pre-switch reward-aligned (90cm-shift) templates (right, F & N) during 20 laps post-switch. F versus F-shift, 0.32±0.035 versus –0.12±0.014, p = 3.9e–5, paired *t*-test, n = 9 mice; N versus N-shift, –0.15±0.025 versus 0.33±0.028, p = 1.4e–4, paired *t*-test, n = 6 mice; Circles: mice; crosses: mean±s.e.m. F/N: familiar/novel environments, day 1. “***”: p<0.001. “**”: p<0.01. **(H)** Same as (G), but for velocity profile correlation. F versus F-shift, 0.4 ± 0.047 versus – 0.24 ± 0.075, p = 6.2e–4, paired *t*-test, n = 9 mice; N versus N-shift, –0.36 ± 0.043 versus 0.49±0.076, p = 1.3e–3, paired *t*-test, n = 6 mice; Circles: individual mice; crosses: mean±s.e.m. F/N: familiar/novel environments, day 1. “***”: p<0.001. “**”: p<0.01. **(I)** Example velocity heatmaps across space and laps in Pre (laps 81-100) and Post (laps 101-120) in a representative same- (familiar env. A, left) and cross-environment (novel env. B, right) mice. Arrows: reward in Pre (black, 50cm) and Post (red, 140 cm). **(J)** Trial-by-trial PF versus running profile similarity (both in c.c. versus pre-switch, reward-aligned template) for familiar (day 1, left) and novel (day 1, right) environments. Each dot: one lap from one mouse. Black line: linear fit. Familiar env.: R = 0.63, p = 8.1e–99, n = 870 points, 9 mice (one mouse had 70 laps after reward switch). Novel env.: R = 0.37, p = 1.2e–20, n = 600 points, 6 mice. **(K)** Population activity from four conditions (F, day 1, pre/post, N, day 2, pre/post) projected onto principal components 1 and 3, revealing overlapping ring structures that allowed generalization. Pooled data from 5 mice. Color: distance from reward (left); contexts (right). F: familiar env., day 1; N: novel env., day 2. Both days had reward switching performed. **(L)** Same as (K), but for principal components 4 and 5 showing environment-specific (F versus N), but relatively reward-position-invariant (F-pre versus F-post; N-pre versus N-post) projections. **(M)** Similar to (K), but showing 3D projection onto principal components 1-3. **(N)** As in (M), but for principal components 4-6.

**Fig. S5.**
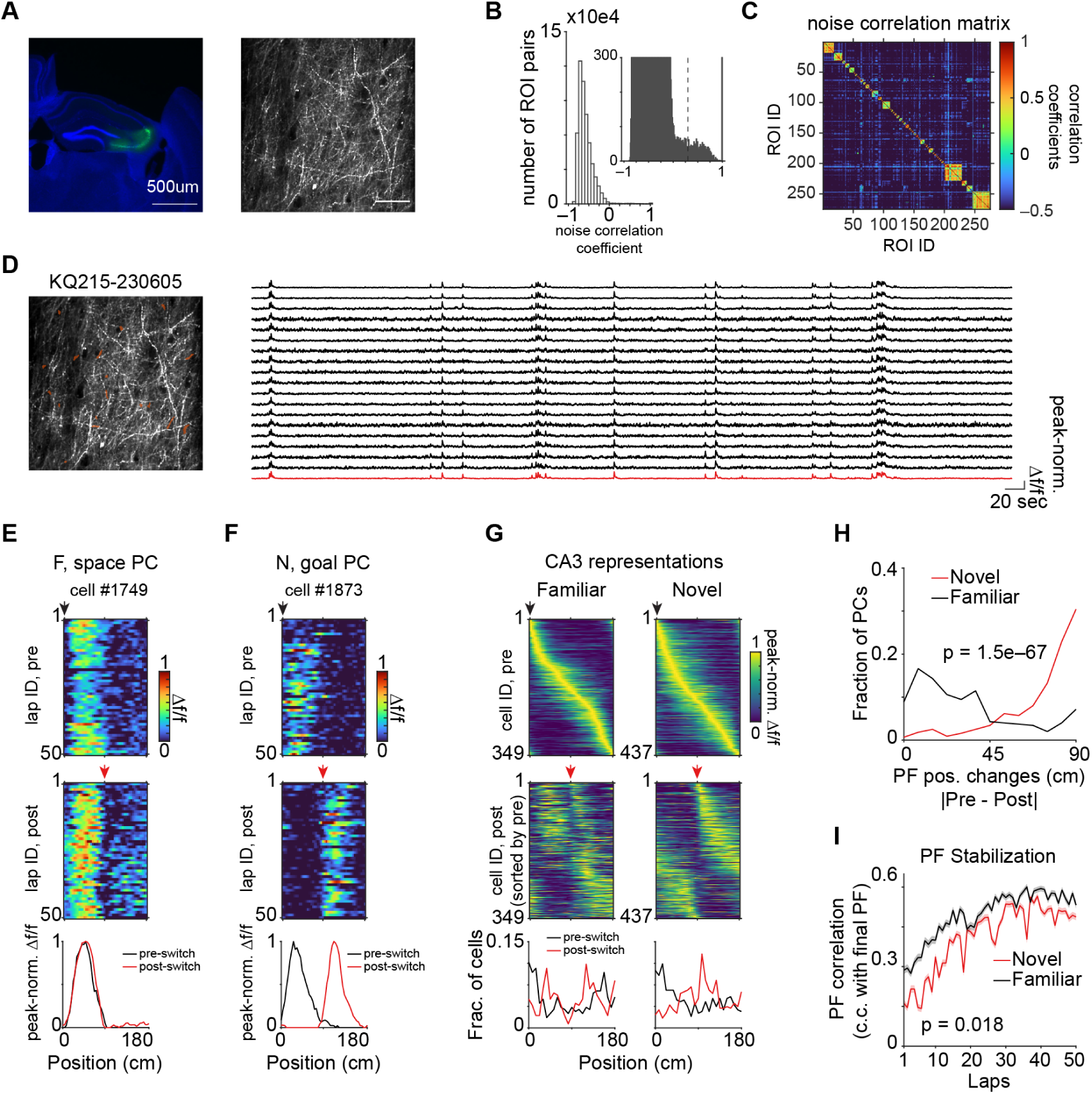
Context-dependent place cell referencing in CA3. **(A)** Left: representative coronal slice (100-*μm* thick, scale bar: 500 *μm*). Blue: DAPI; green: jGCaMP8m. Right: representative two-photon image showing GCaMP8m expression in CA3 axons (scale bar: 50 *μm*). **(B** to **D)** Analysis pipeline to identify regions of interest (ROIs) belonging to the same functional unit. (B) Histogram of noise correlation coefficient distribution for active ROI pairs (see Methods). Inset: magnified view of a distinctly binned histogram with dashed line showing high correlation threshold. (C) Noise correlation matrix for all axonal ROIs with active events identified in this animal. (D) Example of 18 ROIs putatively belonging to same axon. Left: red regions depicting 18 ROIs overlapped with the FOV. Right. Peak-normalized Δf/f traces for these ROIs (black) and resulting single axon (average, red). **(E)** Example space-referenced CA3 PC (familiar) with heatmaps showing spatial Δf/f across laps pre- (top) and post-reward (middle) switch. Bottom: Peak-normalized (norm.) mean Δf/f in Pre (black) and post (red). Black/red arrows: reward at 0 and 90 cm for (E to G). **(F)** As in (E), but for an example goal-referenced CA3 PC in a novel environment. **(G)** Peak-normalized mean Δf/f for all CA3 PCs in Pre (top) and Post (middle, sorted by pre-shift peak) in familiar (left, n = 349 cells, 10 mice) and novel (right, n = 437 axons, 10 mice) environments. Bottom: fraction versus PF peak position during pre (black) and post-switch (red). **(H)** Fraction versus PF peak position change (pre versus post) after reward switch for familiar (env. A, black, n = 349 axons, 10 mice) and novel (env. B, red, n = 437 axons, 10 mice) environments. p = 1.5e–67, two-sample Kolmogorov-Smirnov test. **(I)** Time course of PF correlation from laps 1 to 50 before reward switch (familiar env. A, black; novel env. B, red) relative to the final PFs in Pre. Cross-lap mean, then cross-cell mean yielding 1 mean PF correlation value per animal. Novel versus familiar, 0.39±0.015 versus 0.49±0.013, p = 0.018, paired *t*-test, n = 10 mice.

**Fig. S6.**
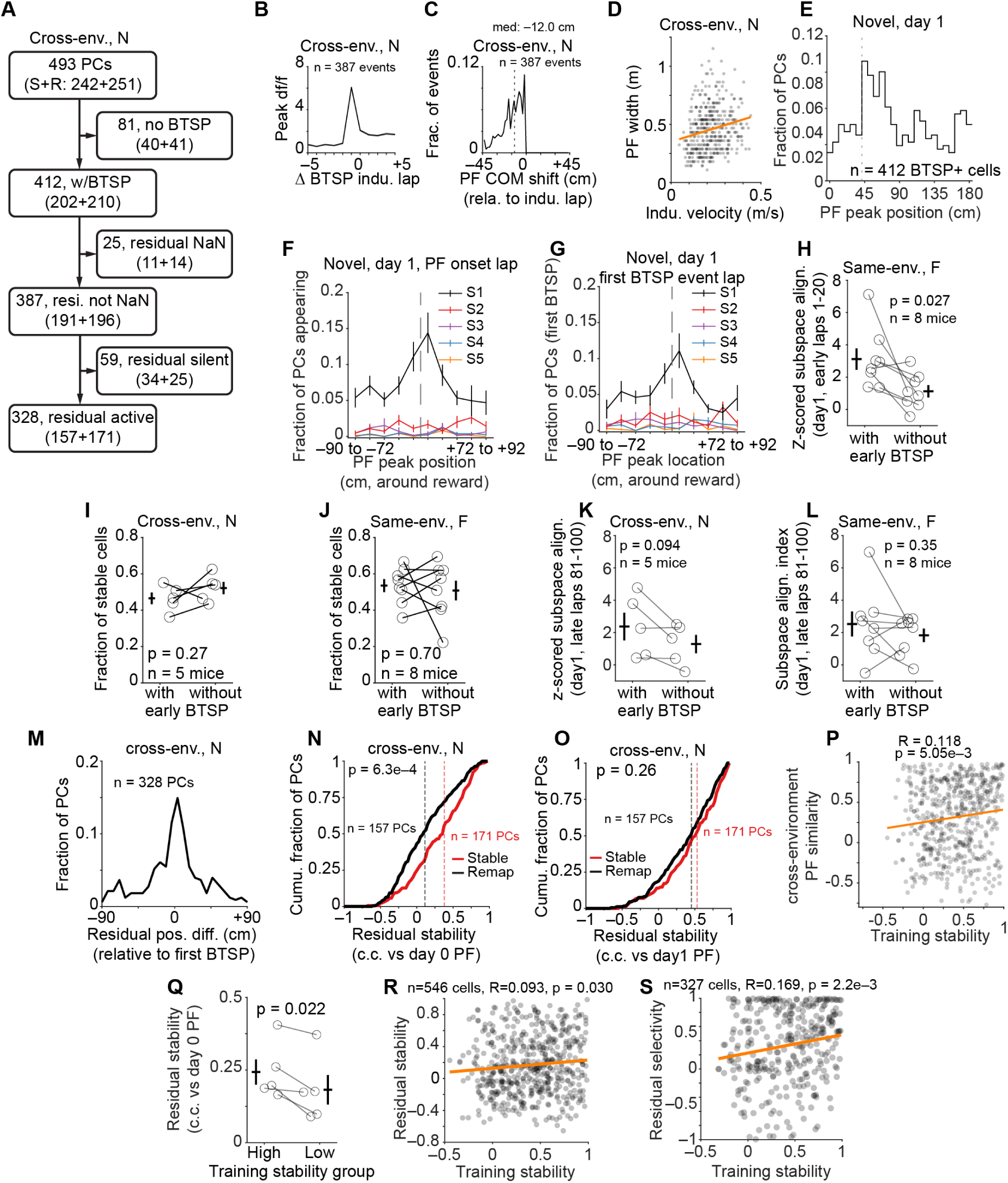
Additional analysis related to BTSP. **(A)** Flowchart illustrating the sequential filtering of CA1 PCs for BTSP related analyses from 6 cross-env. mice. Of 493 PCs (had reliable PFs on days 0 and 1), cells lacking BTSP events, with non-quantifiable residual signals (NaN, e.g., no available preceding laps before BTSP), and residual-silent cells were removed, yielding 328 residual-active PCs. S: Stable; R: Remapped. **(B** to **D)**, BTSP event characteristics for cross-environment mice on day 1 (novel env.). (B) Mean of the peak Δf/f within fields aligned to BTSP induction lap. Before (5-lap mean): 0.74±0.03Δf/f; BTSP Lap: 6.08±0.16; after (5-lap mean): 1.77±0.04; Before versus after, p = 6.2e–40, paired *t*- test, n = 387 events. (C) Event fraction versus PF center of mass shift relative to induction location. Dashed: median shift (–12.0 cm). The sharp boundary at zero reflects the backward-shift requirement imposed by the BTSP detection criteria. (D) PF width following BTSP versus mean velocity during induction. Orange line: linear fit (y = 0.48x+0.35), R = 0.20, p = 5.8e–5. n = 387 events from 6 mice for (B to D). (E) PF peak position distribution for BTSP positive cells on day 1 (novel env. B). Dashed line: reward location at 50 cm. n = 412 BTSP+ cells in 6 mice. (F) Fraction of PCs whose PF onset falls in each 20-lap section and whose PF peak falls in each reward-centered spatial division on day 1 (novel env. B). The first 100 laps were divided into 5 equal sections (S1-S5, 20 laps each), and the 50 spatial bins were grouped into 10 reward-centered divisions (5 bins each, ∼18 cm). Dashed line: reward location. Lines and error bars: mean±sem. N = 6 mice. (G) As in (F), but PCs are binned by the lap of their first BTSP event rather than PF onset lap. Only BTSP positive PCs contribute to the numerator; the denominator remains all traced PCs per mouse. **(H)** Z-scored subspace alignment between day 1 (familiar env. A, early laps 1-20) and day 0 (familiar env. A) population activity, for cells with versus without early BTSP (as in Fig. 3D) in same-environment mice. Z-scores were computed relative to 1000 shuffles of randomly paired subspaces from day 0 (computed separately for each subpopulation), enabling comparison across groups with different cell number or population structure. With: 3.1±0.66; without: 1.1±0.38; p = 0.027, paired *t*-test, n = 8 mice. One excluded (fewer than 20 cells per either group). **(I)** Stable cell fraction (day 0 familiar env. to day 1 novel env.) for PCs with versus without early BTSP (as in Fig. 3D) in cross-environment mice. With: 0.47±0.032; without: 0.52±0.033; p = 0.27, paired *t*-test, n = 5 mice. **(J)** As in (I), but for same-environment. With: 0.54±0.035; without: 0.51±0.054; p = 0.70, paired *t*-test, n = 8 mice. **(K)** z-scored subspace alignment index (late laps 81-100 on day 1 novel versus day 0 familiar) for cells with vs. without early BTSP in cross-environment mice. With: 2.4±0.86; without: 1.3±0.57; p=0.094, paired t-test, n = 5 mice. **(L)** As in (K), but for same-environment mice. With:2.5±0.77; without: 1.8±0.42; p = 0.35, paired *t*-test, n = 8 mice. For (H to L), circles: mice; crosses: mean±s.e.m; one mouse excluded per panel. **(M)** PC fraction versus position difference between residual and subsequent first BTSP event peak cross-environment mice (env. B, day 1). n = 328 cells, 6 mice. **(N)** Cumulative fraction versus residual stability (c.c. versus day 0 PF) for stable (n = 157 cells, 6 mice) and remapped (n = 171 cells, 6 mice) cells on day 1 (novel env.). Black/red dashed: median for remapped (0.11) / stable (0.38) cells; p = 6.3e–4, two-sample Kolmogorov-Smirnov test. **(O)** As in (N), but for day 1 residual stability versus day 1 PF. Black/red dashed lines: median for remapped (0.45) / stable (0.53) cells; p = 0.26, two-sample Kolmogorov-Smirnov test. **(P)** Cross-environment PF similarity (c.c. between day 0 and day 1 mean PFs) versus training stability (mean pairwise PF correlation across 3 training days, see Methods) for day 0 PCs in cross-environment mice tracked across 3 training days (5 mice). Each dot: one cell. Orange line: linear fit. R = 0.12, p = 5.05e–3; **(Q)** Residual stability (c.c. relative to day 0 PFs) for high- (top 20%) versus low- (bottom 20%) training-stability cells on day 1 (novel env. B). Training stability was defined as described above; Top and bottom 20% were defined per mouse (see Methods). For cells with BTSP on day 1, residual was the mean spatial activity of the 5 laps preceding the first BTSP event. For cells without BTSP on day 1, residual activity was computed from a matched 5-lap window (lap 20; laps 15-19; see Methods). High: 0.24±0.044; low: 0.18±0.051; p = 0.022, paired t-test, n = 5 mice. Circles: mice; crosses: mean±sem. **(R)** As in (Q), but showing all individual data points as a scatter plot. Residual stability (c.c. between day 1 Pre-BTSP spatial tuning and day 0 PF) versus training stability for all day 0 PCs. Each dot: one cell. Orange line: linear fit; R = 0.093, p = 0.03, n = 546 cells. **(S)** As in Fig. 3L, but showing all individual data points as a scatter plot. Cells without BTSP or with first-lap BTSP were excluded. Each dot: one cell. Orange line: linear fit; R = 0.17, p = 2.2e– 3, n = 327 cells. F/N: familiar/novel environments for (A to S).

**Fig. S7.**
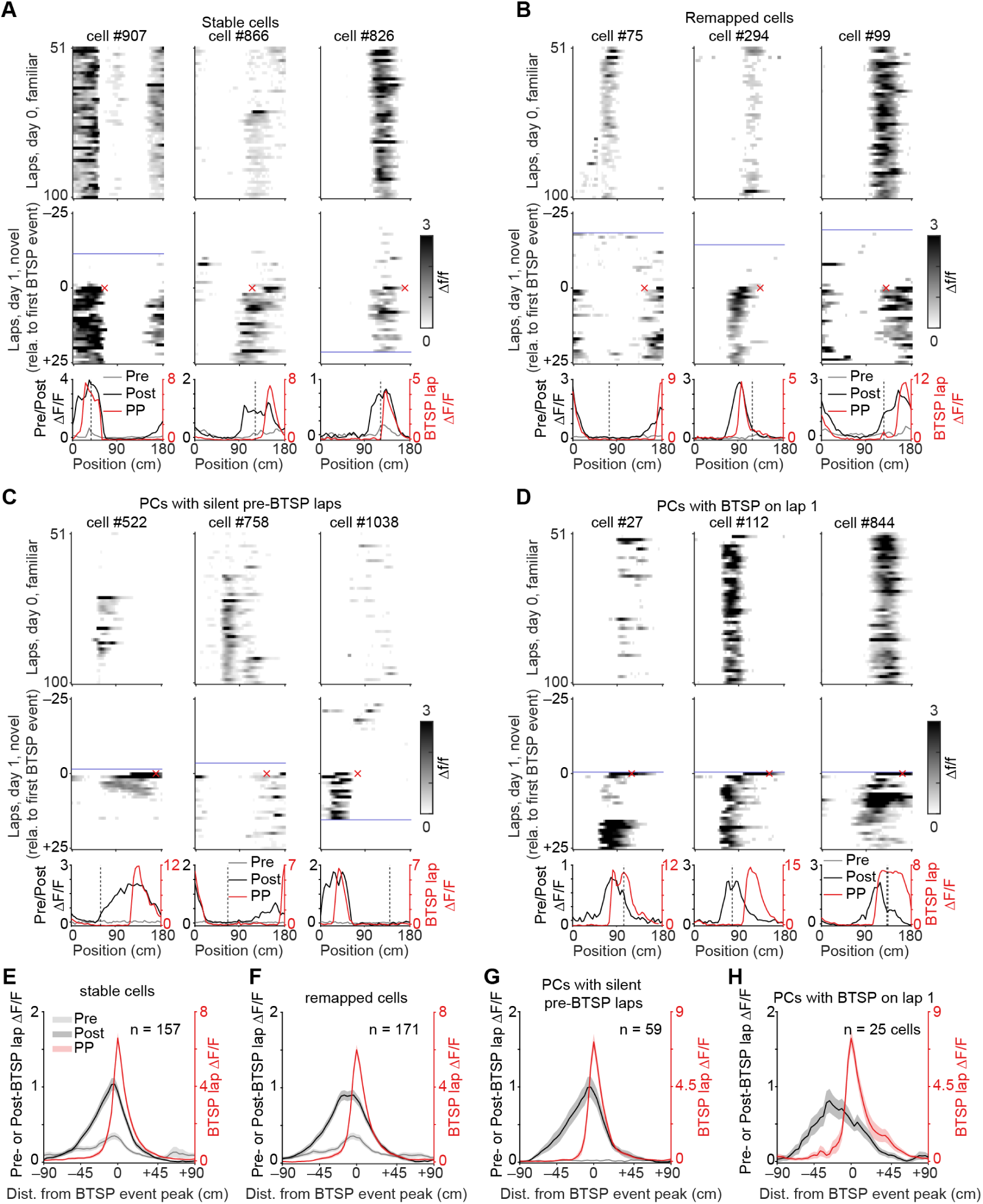
Additional residual examples and population summary aligned to BTSP events. (**A to D**) Three representative cells from each of four categories on day 1 (novel env. B). For each cell: top Δf/f heatmap on day 0 (familiar env. A, laps 51-100); middle, df/f heatmap on day 1, ±25 laps centered on the first BTSP event (red cross, first BTSP event; blue lines, session boundaries); bottom, mean spatial tuning for the 5 pre-BTSP laps (Pre, grey, left y-axis), 5 post-BTSP laps (Post, black, left y-axis), and BTSP event lap (red; right y-axis), with the day 0 PF peak indicated (dashed line). **(A)** stable cells with detectable residual activity aligned to the subsequent BTSP event location. **(B)** as in (A) but for remapped cells. **(C)** place cells with no detectable somatic calcium activity in the 5 laps preceding the first BTSP (silent pre-BTSP laps). **(D)** place cells whose first BTSP event occurred on lap 1, thus no pre-event activity was available. (**E to H**) Population-averaged spatial tuning aligned to the BTSP event peak (position = 0 cm), plotted as in (A-D) bottom panels but averaged across cells within each category. Lines and shading: mean±sem. (E) Stable cells with residual activity (n = 157 cells). (F) Remapped cells with residual activity (n = 171 cells). (G) Place cells with silent pre-BTSP laps ( n = 59 cells). (H) Place cells with BTSP on lap 1 (n = 25 cells). All cells were place cells active on both day 0 and day1 with at least one detected BTSP event on day 1 (6 mice).

**Fig. S8.**
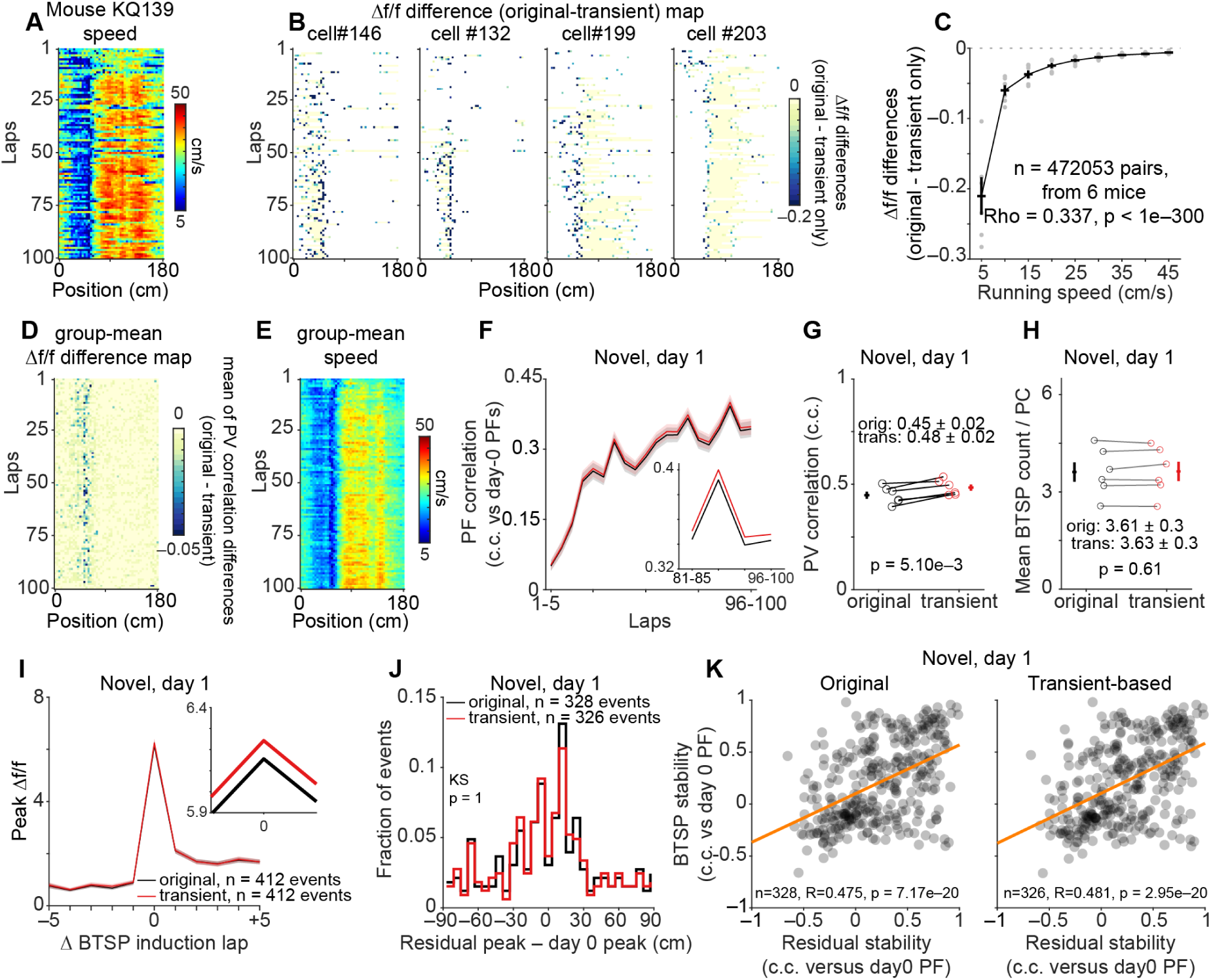
Validation of spatial metrics against running-speed-dependent dwell-time bias. **(A)** Running speed heatmap for a representative cross-environment mouse on day 1 (novel env. B). Each pixel shows the mean speed (cm/s) within one spatial bin (x-axis, 50 bins spanning the 180-cm belt) on one lap (y-axis, 100 laps). **(B)** Per-cell Δf/f difference heatmaps (original minus transient-based pipeline) for four example place cells from the same session as (A). The original pipeline computes the mean Δf/f across all imaging frames within each spatial bin on each lap; the transient-based pipeline restricts this computation to frames containing significant calcium transients (see Methods). Each pixel represents one spatial bin × one lap entry. Color is displayed only at entries where a significant calcium transient was detected; all other entries are masked (white). Negative values (deeper blue) indicate entries where the transient-based estimate exceeded the binned mean, as expected when dwell time dilutes the bin-level average. **(C)** Mean Δf/f difference (original minus transient) as a function of running speed, sampled only at spatial bin × lap entries where a significant transient was detected. Nine speed bins (5 to 45 cm/s). Grey dots: per-mouse bin means; black line and error bars: group mean ± s.e.m. Dashed line: zero. Spearman ρ computed on all raw (speed, Δf/f difference) pairs. ρ = 0.337, p < 1e–300, n = 472,053 pairs from 6 mice. The positive correlation indicates that the divergence between pipelines is largest at low running speeds, where dwell time per bin is longest. **(D)** Group-mean difference in population vector (PV) correlation (original minus transient pipeline) across spatial bins (x-axis) and laps (y-axis). For each spatial bin on each lap, PV correlation was computed as the Pearson’s c.c. between the day 0 (familiar env. A) and day 1 (novel env. B) population vectors at that bin, using all PCs. The difference between pipelines was then averaged across mice. n = 6 mice. **(E)** Group-mean running speed (cm/s) across spatial bins and laps for the same mice as (D). **(F)** Lap-by-lap PF correlation (Pearson’s c.c.) between each cell’s day 1 (novel env. B) per-lap spatial profile and its day 0 (familiar env. A) mean spatial profile, averaged across stable PCs (place cells active with stable PF locations on both days). Black: original pipeline; red: transient-based pipeline. Lines and shading: mean ± s.e.m. Inset: zoomed view of the last four 5-lap sections (laps 81–85 through 96–100). n = 6 mice. **(G)** Per-mouse PV similarity (mean diagonal Pearson’s c.c. between day 0 and day 1 mean spatial profiles across reliable PCs) for original versus transient-based pipelines. Each pair of connected dots: one mouse. Crosses: mean ± s.e.m. Original: 0.45 ± 0.02; transient: 0.48 ± 0.02; p = 5.10e– 3, paired t-test, n = 6 mice. **(H)** Per-mouse mean BTSP event count per place cell for original versus transient-based pipelines. Format as (G). Original: 3.61 ± 0.3; transient: 3.63 ± 0.3; p = 0.61, paired t-test, n = 6 mice. **(I)** Mean peak Δf/f within each cell’s place field, centered on the first BTSP induction lap (lap 0), spanning 5 laps before (–5) to 5 laps after (+5). Black: original pipeline; red: transient-based pipeline. Inset: zoomed at BTSP event lap. n = 412 events from 6 mice for both pipelines. Only PCs with at least one BTSP event on day 1 were included. **(J)** Fraction of events versus position difference between the residual activity peak (mean spatial profile from 5 pre-BTSP laps on day 1) and the day 0 (familiar env. A) PF peak (in cm), for original (black, n = 328 events) and transient-based (red, n = 326 events) pipelines overlaid. Only residual-active PCs were included (see fig. S7A; S7B). p = 1, two-sample Kolmogorov-Smirnov test. **(K)** BTSP stability (Pearson’s c.c. between each cell’s BTSP event lap spatial tuning and day 0 PF) versus residual stability (Pearson’s c.c. between pre-BTSP spatial tuning and day 0 PF) for original (left, n = 328 events, R = 0.48, p = 7.2e–20) and transient-based (right, n = 326 events, R = 0.48, p = 3e–20) pipelines. Orange line: linear fit. Each dot: one cell. Only PCs passing the residual-active filter (place cell on both days, ≥1 BTSP event, detectable pre-BTSP activity) were included. Only first BTSP event was included per cell. F/N: familiar/novel environments for all panels.

**Fig. S9.**
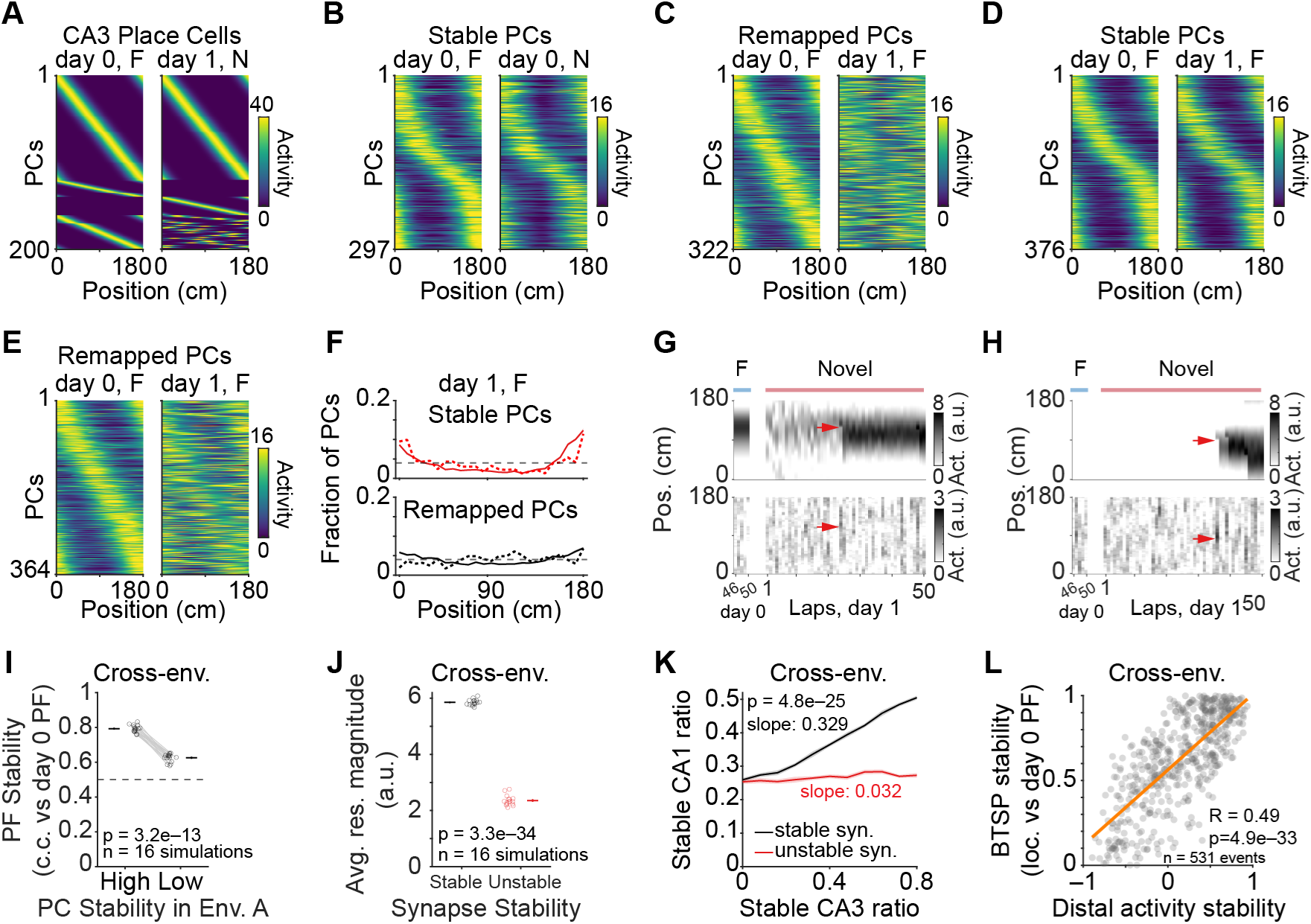
Additional analysis related to the computational model. **(A)** CA3 input profiles in the model showing firing activity in space in familiar (F: env. A) and novel (N: env. B) environments (sorted by activity peak in F, except that novel environment cue specific neurons were sorted by peak in N). Cells sorted separately for task-input (top), cue-input (middle), and remapping inputs (bottom). **(B)** Mean firing activity of stable cells in familiar env. A (left, day 0, sorted by PF peak position) and novel env. B (right, day 1, sorted by F). **(C)** Same as (B), but for remapped cells. **(D** to **E)**, Same as (B to C), but for stable (D) and remapped (E) cells in same-environment conditions. **(F)** Same as Fig. 4D, but for same-environment conditions. **(G)** Simulated peri-somatic (top, “CA3”) and distal dendritic (bottom, “EC”) activity of one example cell across space and laps from day 0 (familiar) to day 1 (novel). Note spatially-informative residual from CA3, combined with EC input, biases plateau initiation timing that induces BTSP (red arrow). **(H)** Same as (G), but for a representative cell without reliable residual activity. Red arrow: first plateau driven by strong EC input alone, producing strong PF in one shot. **(I)** Simulated PF stability (c.c. versus day 0 familiar env. A) in the novel environment B for cells with high versus low stability during training in env. A. High: 0.79±0.0051; low: 0.63±0.0058; p = 3.2e−13, paired two-sample two-sided *t*-test, n = 16 simulations. Circles: simulations; crosses: mean±s.e.m). **(J)** Simulated residual activity magnitudes across all cells in novel environment under control (black, stable synapses) and impaired CA3-CA1 synaptic stability (red, unstable synapses). Stable: 5.9 ± 0.026 a.u.; unstable: 2.4 ± 0.045 a.u., p = 3.3e–34, unpaired two-sided t-test, n = 16 simulations. Circles: simulations; crosses: mean±s.e.m. Removal of cascade (see Methods): 4.8±0.0030, p = 1.0e–21 (versus stable), unpaired two-sided *t*-test, n = 16 simulations. **(K)** Stable CA1 PC ratio versus stable CA3 PC ratio across environments for both control (black) and impaired CA3-CA1 synaptic stability (red). Solid: mean; shading: ± SEM; stable: 0.37±0.0025; unstable: 0.27±0.0020; p = 4.8e–25, unpaired two-sided *t*-test on AUC, n = 16 simulations. Removal of cascade (see Methods): 0.31±0.0018, slope=0.142, p = 3.2e–18 (versus stable), unpaired two-sided *t*-test on AUC, n = 16 simulations. **(L)** Relationship between cross-environment distal dendritic activity stability (driven by EC, in c.c.) and BTSP (plateau) location stability (in cm; both versus day 0 PFs) in an example simulation. R = 0.49, p = 4.9e–33, n = 531 cells (same simulation as Fig. 4E). p < 0.05 across all 16 simulations.

**Fig. S10.**
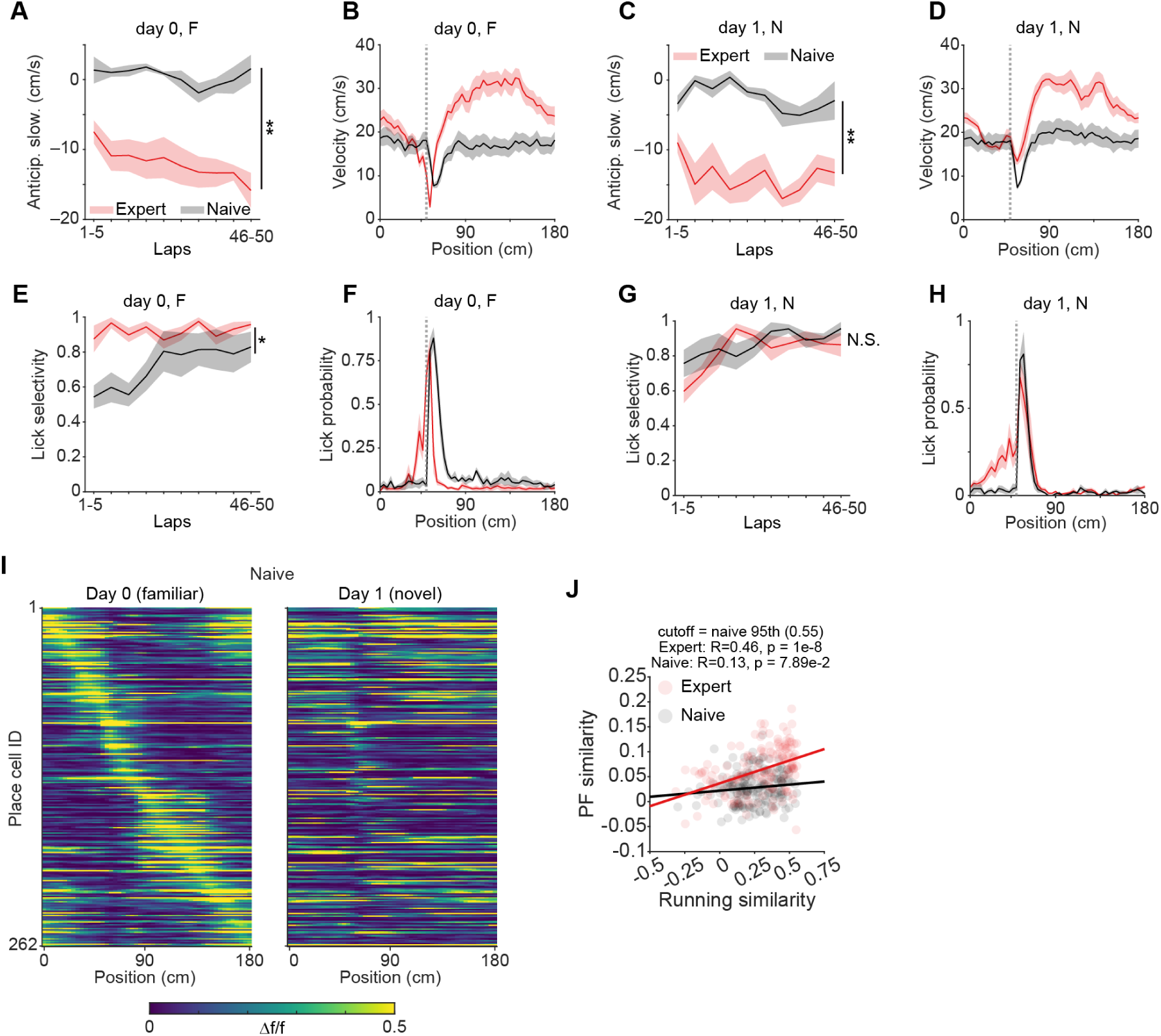
Behavioral comparison between expert and naïve mice. **(A)** Anticipatory slowing (see Methods) across 5-lap sections on day 0 in the familiar environment (belt A). Expert (red, n = 6 mice) versus naïve (grey, n = 4 mice). Lines and shading: mean±s.e.m. Two-way repeated ANOVA: group p = 2.7e–3; section p = 5.1e–2; interaction p = 0.20. Asterisk denotes significant group difference (**p<0.01). F: familiar environment. **(B)** Mean velocity versus position on day 0 (familiar env.) for expert (red) and naïve (grey). Dashed line: reward location. Lines and shading: mean±s.e.m. F: familiar environment. **(C)** As in (A), but for day 1 in the novel environment (belt B). Two-way repeated ANOVA: group p = 3.05e–3; section p = 0.16; interaction p = 0.13. N: novel environment. **(D)** As in (B), but for day 1 in the novel environment. N: novel environment. **(E)** Lick selectivity (fraction of licks within the reward zone relative to total licks) across 5-lap sections on day 0 (familiar env.). Expert (red) versus naïve (grey). Two-way repeated ANOVA: group p = 0.039; section p = 0.04; interaction p = 0.07. Asterisk denotes significant group difference (*p<0.05). F: familiar environment. **(F)** Mean lick probability versus position on day 0 (familiar env.) for expert (red) and naïve (grey). Dashed line: reward location. F: familiar environment. **(G)** As in (E), but for day 1 (novel env. B). Two-way repeated ANOVA: group p = 0.57; section p = 0.012; interaction p = 0.22. N.S.: not significant. N: novel environment. **(H)** As in (F), but for day 1 (novel env.). Naïve mice licked primarily after reward delivery rather than anticipatorily. N: novel environment. **(I)** Spatial Δf/f heatmaps for all day 0 PCs in naïve mice (n = 262 cells, 4 mice), sorted by day 0 PF peak position. Left: day 0 (familiar env.); right: day 1 (novel env.). Same sort order applied across both days. **(J)** Trial-by-trial PF similarity versus running profile similarity (both in c.c. versus day 0 familiar template) for expert (red) and naïve (black) mice, after restricting both groups to the running-similarity range sampled by naïve mice (cutoff = naïve 95th percentile, 0.55). Expert: R = 0.46, p = 1.0e–8; naïve: R = 0.13, p = 0.08. Each dot: one lap from one mouse (first 50 laps).

**Fig. S11.**
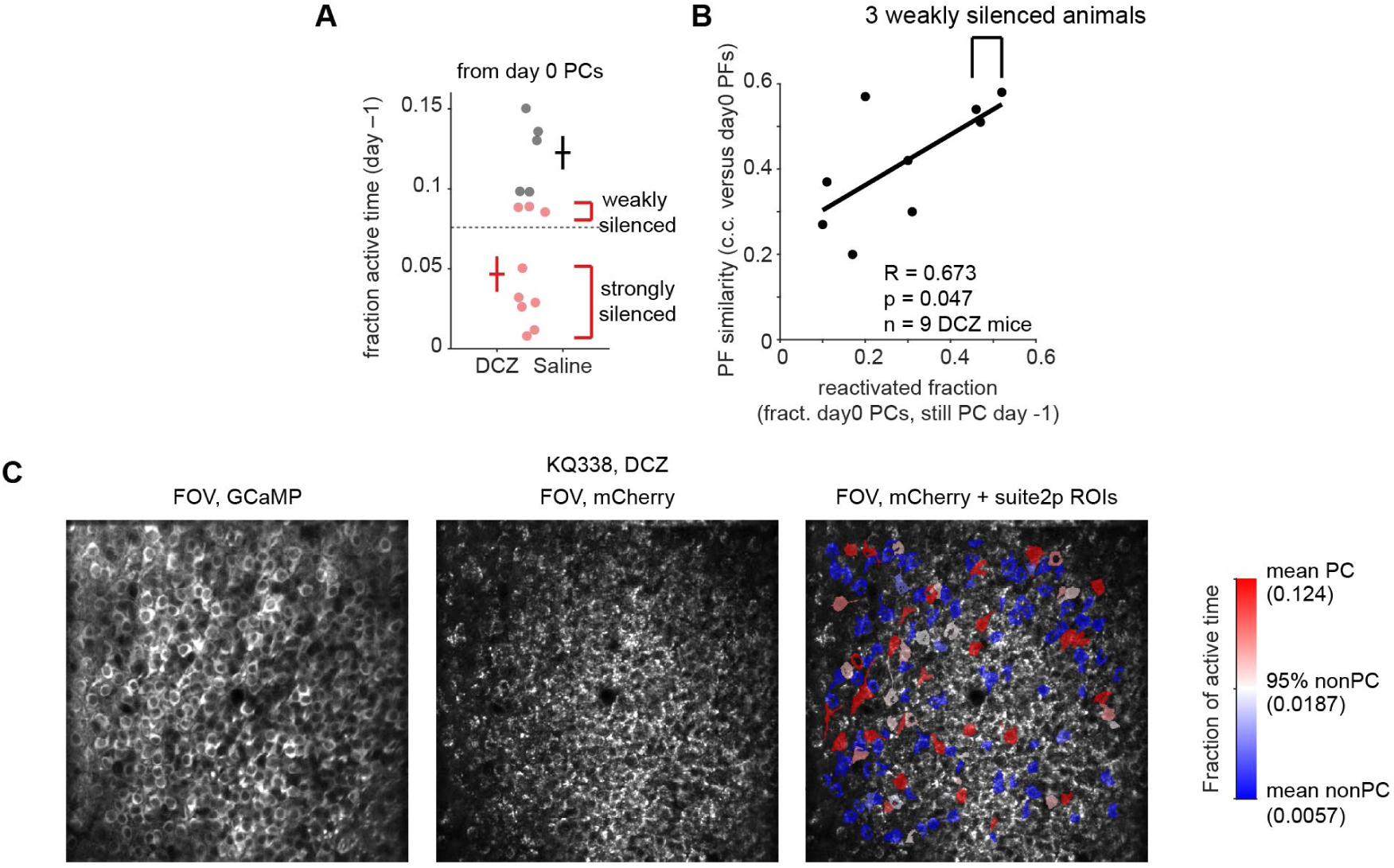
DREADDs silencing verification and exclusion criteria. **(A)** Fraction of active time on day –1 (on-drug) for day 0 PCs in DCZ (red, n = 9 mice) and saline (black, n = 5 mice) groups. Activity was quantified as the fraction of running frames with significant calcium transients, averaged over cells classified as PCs on day 0 (drug-off session). Dashed line: threshold for exclusion (saline mean – 2 SD = 0.076). DCZ mice were classified as “strongly silenced” (below threshold, n = 6) or “weakly silenced” (above threshold, n = 3). Three weakly silenced mice were excluded from the main analyses. Saline: 0.12±0.01. Circles: individual mice; crosses: mean±s.e.m. **(B)** PF similarity (c.c. between day 0 and day 1 mean PFs) versus reactivation fraction (fraction of day 0 PCs that were also PCs on day –1) across all 9 DCZ mice. R = 0.673, p = 0.047, Pearson’s correlation. Bracket: the 3 weakly silenced animals clustered at high reactivation fraction and high PF similarity, indicating insufficient disruption of consolidation history. Black line: linear fit. **(C)** Representative field of view from a DCZ mouse (KQ338). Left: GCaMP6f mean image. Middle: mCherry fluorescence showing Gi-DREADDs expression. Right: mCherry FOV overlaid with Suite2p ROIs colored by fraction of active time on day 0 (off-drug). Color scale aligned to day 0 non-PC distribution: blue = mean non-PC activity (0.0057); white = 95th percentile non-PC (0.0187); red = mean PC activity (0.124). Scale bar: 100 μm.

**Fig. S12.**
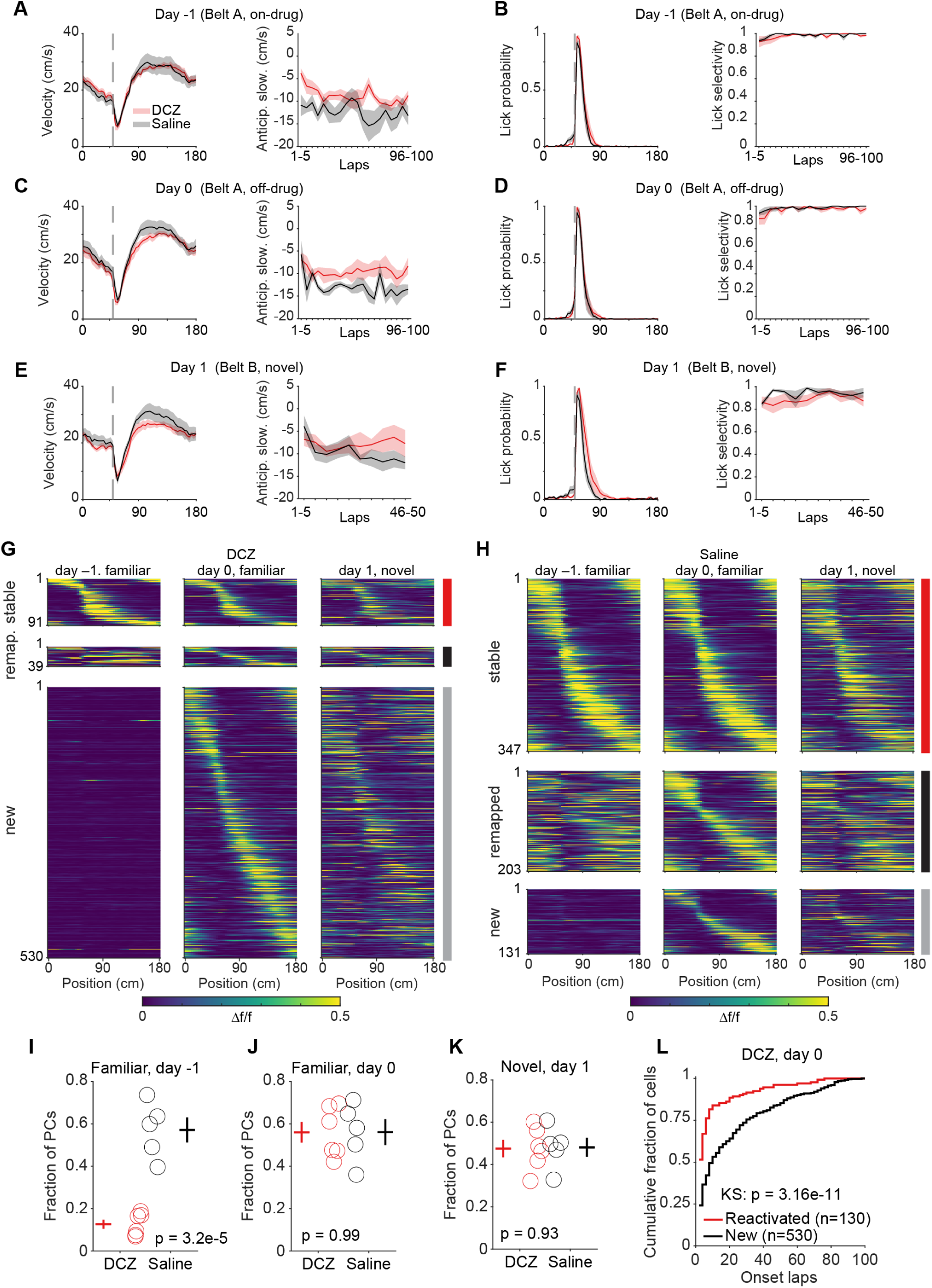
DREADDs behavioral and cellular characterization across three testing days. **(A)** Velocity profile (left) and anticipatory slowing across 5-lap sections (right) for DCZ (red, n = 6 mice) and saline (black, n = 5 mice) on day –1 (belt A, on-drug). Two-way repeated ANOVA: group p = 0.12; section p = 0.17; interaction p = 0.15. Dashed line in velocity panel: reward location. **(B)** Lick probability profile (left) and lick selectivity across 5-lap sections (right) on day –1. Two-way repeated ANOVA: group p = 0.59; section p = 0.13; interaction p = 0.67. **(C)** As in (A), but for day 0 (belt A, off-drug). Two-way repeated ANOVA on anticipatory slowing: group p = 0.081; section p = 0.078; interaction p = 0.39. **(D)** As in (B), but for day 0. Two-way repeated ANOVA on lick selectivity: group p = 0.43; section p = 0.12; interaction p = 0.64. **(E)** As in (A), but for day 1 (belt B, novel). Two-way repeated ANOVA on anticipatory slowing: group p = 0.36; section p = 0.37; interaction p = 0.45. **(F)** As in (B), but for day 1 (novel). Two-way repeated ANOVA on lick selectivity: group p = 0.42; section p = 0.19; interaction p = 0.33. **(G)** Spatial Δf/f heatmaps for DCZ group day 0 PCs classified by day –1 history. Rows: stable (PC on day –1 and day 0, with stable PF; n = 91 cells), remapped (PC on both days, PF changed (>15cm); n = 39 cells), and new (PC on day 0, not PC on day –1; n = 530 cells). Columns: day – 1 (familiar, on-drug), day 0 (familiar, off-drug), day 1 (novel, off-drug). Cells sorted by day 0 PF peak position; same sort order applied across all three days. Note that the combination of stable and remapped is all reactivated cells. Colored bars at right: stable (red), remapped (black), new (grey).\ **(H)** As in (G), but for saline group. Stable: n = 347 cells; remapped: n = 203 cells; new: n = 131 cells. **(I)** PC fraction on day –1 (on-drug) for DCZ (0.13±0.022) and saline (0.57±0.059). p = 3.2e–5, unpaired *t*-test. n = 6 DCZ, 5 saline mice. Circles: individual mice; crosses: mean±s.e.m. **(J)** PC fraction on day 0 (off-drug). DCZ: 0.56±0.048; saline: 0.56±0.06; p = 0.99, unpaired *t*-test. Format as (I). **(K)** PC fraction on day 1 (novel). DCZ: 0.48±0.041; saline: 0.48±0.045; p = 0.93, unpaired *t*-test. Format as (I). **(L)** Cumulative distribution of PF onset lap on day 0 for reactivated (red, n = 130 cells) versus new (black, n = 530 cells) PCs in DCZ mice. p = 3.2e–11, two-sample Kolmogorov-Smirnov test. Lines and shading in (A–F): mean±s.e.m. For all panels, n = 6 DCZ mice, 5 saline mice.

**Fig. S13.**
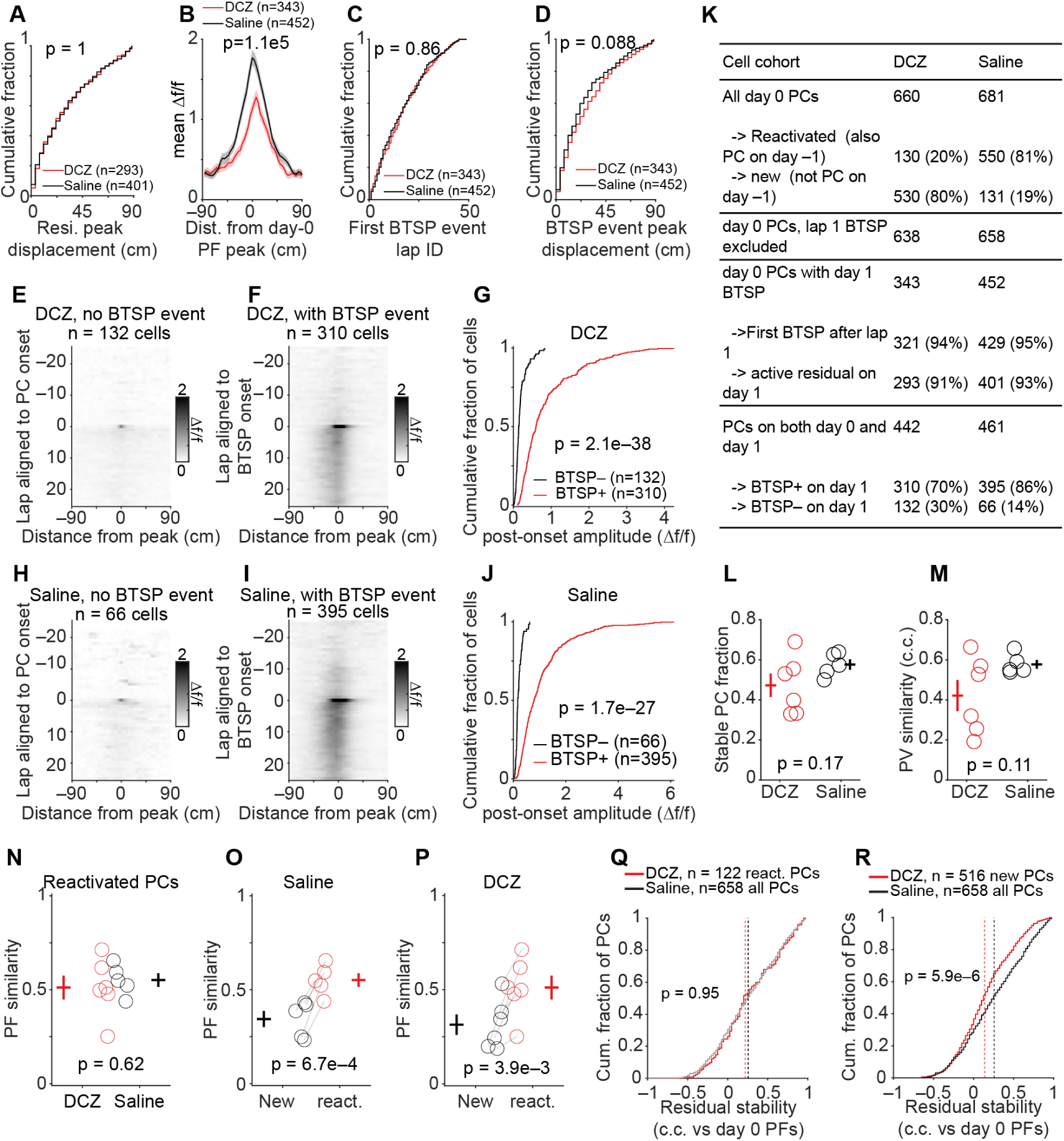
Additional residual and BTSP analyses in DCZ and saline mice. **(A)** Cumulative distribution of the distance between each cell’s day 1 residual peak and its day 0 PF peak, for day 0 PCs with an active residual. DCZ (red, n = 293 cells, 6 mice) versus Saline (black, n = 401 cells, 5 mice), medians 25.2 versus 25.2 cm, p = 1, two-sample Kolmogorov-Smirnov test. **(B)** Population-averaged day 1 BTSP event lap Δf/f, circularly aligned to each cell’s day 0 PF peak rather than to the event peak, for day 0 PCs with a day 1 BTSP. DCZ in red, n = 343 cells, 6 mice; Saline in black, n = 452 cells, 5 mice. Lines and shading: mean±s.e.m. Mean Δf/f at the day 0 peak: DCZ 1.12 versus saline 1.77, p = 1.1e–5, two-sample Kolmogorov-Smirnov test. **(C)** Cumulative distribution of the lap of the first day 1 BTSP event, same cells as (B). DCZ (red, n = 343 cells, 6 mice), Saline (black, n = 452 cells, 5 mice). Medians 15.0 versus 14.5 laps, p = 0.86, two-sample Kolmogorov-Smirnov test. **(D)** Cumulative distribution of the distance between the first BTSP event peak and the day 0 PF peak, same cells as (B). Medians 21.6 versus 18.0 cm for DCZ versus saline mice, p = 0.088, two-sample Kolmogorov-Smirnov test. **(E, F)** Population-averaged day 1 Δf/f across laps and position for cells in DCZ mice with a PF on both days 0 and 1, split by whether a BTSP event was detected on day 1. (E) without a BTSP event, n = 132 cells, aligned by lap to PC onset and by position to the peak of that onset lap. (F) with a BTSP event, n = 310 cells, aligned to the first BTSP event lap and its peak **(G)** Cumulative distribution of post-onset amplitude for the cells with and without BTSP events on day 1 in DCZ mice. For each cell, the mean PF was computed over every lap from its own alignment lap to the end of the session, and the peak of that mean profile within ±4 bins (14.4cm) of the aligned center was taken. Median 0.14 (BTSP–, n = 132 cells) versus 0.60 (BTSP+, n = 310 cells). p = 2.1e–38, two-sample Kolmogorov-Smirnov test. **(H, I, J)** As in (E,F,G) for saline mice. n = 66 cells (H), and 395 cells (I). Median 0.17 (BTSP–) versus 0.79 (BTSP+), p = 1.7e–27, two-sample Kolmogorov-Smirnov test. **(K)** Cell counts for each cohort used in Fig. 5 and Fig. S13 for DCZ and saline mice. Percentages are relative to the nearest preceding total. **(L)** Stable PC fraction, defined as the fraction of cells that had stable PF locations out of all cells that are PCs on both days 0 and 1. DCZ (red, 0.47±0.059, n = 6 mice) and saline (black, 0.58±0.026, n = 5 mice). p = 0.17, unpaired t-test. Circles, individual mice. Crosses, mean±s.e.m. **(M)** Population vector similarity between days 0 and 1, using cells with a PF on both days. DCZ (red, 0.42±0.078, n = 6 mice) and saline (black, 0.58±0.022, n = 5 mice). p = 0.11, unpaired t-test. Circles, individual mice. Crosses, mean±s.e.m. **(N)** Cross-environment PF similarity (c.c. between day 0 and day 1 mean PFs) for reactivated PCs (cells that were PCs on both day –1 and day 0) in DCZ (0.51±0.063) and saline (0.55±0.036) groups. p = 0.62, unpaired *t*-test. n = 6 DCZ and 5 saline mice. Circles: individual mice; crosses: mean±s.e.m. **(O)** PF similarity for new versus reactivated PCs in saline mice. New: 0.34±0.043; reactivated: 0.55±0.036; p = 6.7e–4, paired *t*-test, n = 5 mice. Circles: individual mice; crosses: mean±s.e.m. **(P)** As in (O), but for DCZ mice. New: 0.32±0.054; reactivated: 0.51±0.063; p = 3.9e–3, paired *t*-test, n = 6 mice. **(Q)** Cumulative distribution of residual stability, the correlation between day 1 activity preceding the first BTSP event and the day 0 PF for DCZ reactivated day 0 PCs (red, n = 122 cells) versus saline all day 0 PCs (black, n = 658 cells). Cells with no day 1 BTSP event used five-lap window before a reference lap shared by both groups (lap 17; Methods). Cells whose first event fell on lap 1 are excluded, having no preceding laps. p = 0.95, two-sample Kolmogorov-Smirnov test. Dashed vertical lines: median for each group. **(R)** Cumulative distribution of residual stability for DCZ new PCs (red, n = 516 cells) versus saline all PCs (black, n = 658 cells). p = 5.9e–6, two-sample Kolmogorov-Smirnov test. Dashed vertical lines: median for each group.

**Fig. S14.**
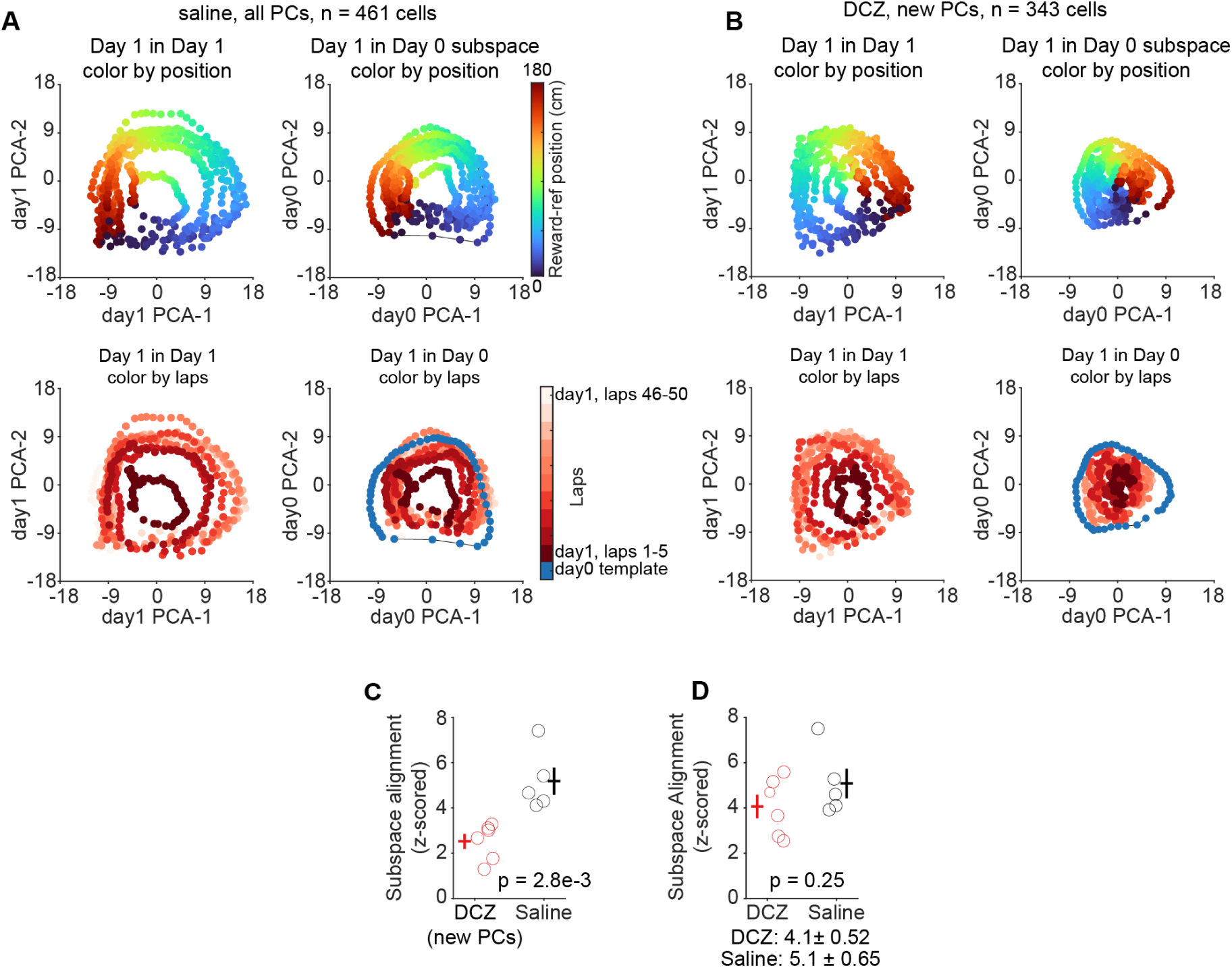
Low-dimensional subspace alignment between day 0 and day 1 in DCZ and saline mice. **(A)** Population activity visualization for saline group. Mean spatially binned Δf/f maps (50 laps, 50 spatial bins per cell) from all PCs that have reliable PFs on both day 0 and day 1 were concatenated across mice to form a single pseudo-population (n = 461 cells, 5 mice), and PCA was performed on this concatenated spatial representation. Lap-by-lap, bin-by-bin day 1 (novel env. B) activity projected onto day 1 principal components (left) and day 0 principal components (right) and then averaged per 5 laps for visualization. Top row: colored by reward-referenced position (cm). Bottom row: colored by lap number. This spatial-map concatenation was used solely for visualization; all quantitative measures of subspace alignment were computed from time-binned population Δf/f activity within individual mice as described in Methods. **(B)** As in (A), but for DCZ new PCs on day 0 that also had reliable PFs on day 1 (n = 343 cells). Note visually weaker ring structure when projected onto day 0 subspace (right column), indicating reduced alignment with the familiar representation compared to saline controls. **(C)** Z-scored subspace alignment between day 1 (novel env. B) and day 0 (familiar env. A) population activity. Here DCZ group used new PCs only (PCs on day 0 that were not PCs on day –1), whereas saline group used all PCs on day 0. DCZ (new PCs): 2.5±0.3, n = 6 mice; saline (all PCs): 5.2±0.6, n = 5 mice; p = 2.8e–3, unpaired *t*-test. Z-scores computed relative to 1000 shuffles of randomly selected 2 subspaces from day 0. Circles: individual mice; crosses: mean±s.e.m. **(D)** As in (C), but using all PCs for both groups. DCZ 4.1±0.52, saline 5.1±0.65, p = 0.25, unpaired t-test.

